# Placing very long branch taxa in the plant tree of life: a case study with Dioscoreales mycoheterotrophs

**DOI:** 10.1101/2025.09.22.677722

**Authors:** Marybel Soto Gomez, Qianshi Lin, Nathaniel J. Klimpert, Vivienne K. Y. Lam, Juan Viruel, Vincent S. F. T. Merckx, Paula J. Rudall, Sean W. Graham

## Abstract

**Premise:** The inclusion of heterotrophic plants in broadly focused systematic studies is challenging, reflecting substantial morphological modification, gene loss, and accelerated nucleotide substitution following photosynthesis loss. In particular, inclusion of very long branch (VLB) taxa with highly elevated rates may lead to phylogenomic misinference.

**Methods:** We explore how rapidly evolving plastomes behave in model-based phylogenetic inferences of the yam order Dioscoreales, which experienced convergent photosynthesis losses. The taxon sampling includes all photosynthetic genera, and mycoheterotrophic lineages with a broad range of substitution rates.

**Results:** When only moderately rapidly evolving heterotrophs are included, relationships within Dioscoreales are congruent with recent mitochondrial analysis, and generally strongly supported. Photosynthetic *Stenomeris* (represented by only three genes) is moderately supported as sister to Dioscoreaceae. Notably, Burmanniaceae and Thismiaceae, two mycoheterotrophic families including highly reduced plastomes, are strongly supported as distantly related within Dioscoreales. Constrained analysis rejects combining Thismiaceae with Burmanniaceae, conflicting with current angiosperm classification. Including VLB mycoheterotrophs in phylogenetic inference can lead to localized to broad reduction in branch support—substantially so for Afrothismiaceae, a family placing in disparate locations in Dioscoreales (and beyond) in variant analyses, with no ability to differentiate among its alternative placements.

**Conclusions:** Inclusion of highly rate-elevated taxa in analyses of Dioscoreales phylogeny can depress branch support. Inferred relationships in the order are otherwise congruent with studies based on mitochondrial data; the family-level classification needs updating. The extraordinarily rapidly evolving family Afrothismiaceae places inconsistently in diverse plastid analyses, pointing to probable analytical limits for plastid-based phylogenomic analyses of VLB heterotrophs.

## INTRODUCTION

Phylogenomic datasets based on the plastid (chloroplast) genome have facilitated well supported inferences of higher-level relationships both within and among most orders of monocots (e.g., Givnish et al., 2010, 2015, 2016, 2018; Steele et al., 2012; Barrett et al., 2013, 2014, 2016, 2024a; Henriquez et al., 2014; Logacheva et al., 2014; Comer et al., 2015; Mennes et al., 2015; Ross et al., 2015; Sass et al., 2016; Lam et al., 2018; Soto Gomez et al. 2020; Li et al., 2019, 2021; Zuntini et al., 2021). Phylogenomic approaches have not yet been attempted on an extensive scale for the yam order Dioscoreales, although Garrett et al. (2023) employed plastid genome data for a constituent family (Thismiaceae), and Givnish et al. (2018) and Lam et al. (2018) included several exemplar taxa in broader phylogenetic studies. Nonetheless, this order is perhaps the last major monocot clade in need of detailed attention using phylogenomic data (e.g., Trias-Blasi et al., 2015). A major complicating factor in addressing higher-order relationships in Dioscoreales is the repeated origin of fully mycoheterotrophic (FM), non-photosynthetic taxa in the order (Merckx et al., 2008a, 2010, 2013). These lineages, like heterotrophic plants in general, are often highly modified morphologically, and can display substantially elevated substitutional rates that can be problematic in phylogenetic or phylogenomic inference (e.g., Neyland and Hennigan, 2003; Nickrent et al., 2004; Barkman et al., 2007; Merckx et al., 2009a; Su et al., 2015; Bellot et al., 2016; Lam et al., 2016, 2018; Naumann et al., 2016; Shepeleva et al., 2020—and see Felsenstein, 1978; Hendy and Penny, 1989, on the general problem of long-branch attraction).

Fully mycoheterotrophic plants obtain fixed carbon from root-associated fungal partners rather than through photosynthesis (Leake, 2004; Merckx et al., 2009b). Therefore, the strong selective constraints that normally operate in genes associated with photosynthetic function can be completely lost, leading to rapid gene loss (e.g., Barrett et al., 2014; Graham et al., 2017). Other retained genes with non-photosynthetic functions can experience relaxed purifying selection (e.g., Garrett et al., 2023). This can contribute to elevated DNA substitution rates, but elevated substitution in general likely relates to higher overall rates of mutation reflecting reduction in the efficiency of plastid DNA replication and repair machinery in heterotrophs (e.g., Wicke et al. 2016). The mitochondrial genome is known to be less affected by substitutional rate elevation and gene loss in heterotrophic plants, but has occasionally been applied to phylogenetic inference in these lineages using mitochondrial genes (e.g., Merckx et al., 2009a; Mennes et al., 2013) or genomes (Soto Gomez et al., 2020; Lin et al., 2022; Barrett et al., 2024).

Plastid sequences still underlie much of our knowledge of higher-order plant relationships, highlighting their continuing importance for integrating previously unsequenced or problematic lineages into the plant tree of life. The photosynthesis-related genes *atp*B and *rbc*L are two of the three major loci that broadly underpin the Angiosperm Phylogeny Group classification systems (APG, 1998, 2003, 2009, 2016). However, both genes are generally missing from the plastid genomes of fully heterotrophic plants (e.g., Graham et al., 2017; Lam et al., 2018; Wicke and Naumann, 2018). Nonetheless, multiple phylogenomic studies have shown that retained plastid genes unrelated to photosynthesis in heterotrophic plants can be employed effectively in phylogenetic analyses, despite sometimes extensive gene loss, and often substantial rate-elevation in the included heterotrophic lineages (e.g., Delannoy et al., 2011; Logacheva et al., 2014; Givnish et al., 2015, 2018; Lam et al., 2015, 2018; Mennes et al., 2015; Bodin et al., 2016; Feng et al., 2016; Barrett et al., 2018, 2019, 2024a,b; Lee et al., 2020; Soto Gomez et al., 2020; Yudina et al., 2021; Klimpert et al., 2022; Garrett et al., 2023). Indeed, a broad survey by Lam et al. (2018) to place the major lineages of mycoheterotrophs on the angiosperm tree of life demonstrated that taxa with elevated rates—and often relatively few retained genes—can be successfully incorporated in plastid phylogenomic analyses. Nonetheless, their study and others also provided evidence of strong long-branch artefacts for a subset of mycoheterotrophic monocots displaying more extreme rates of DNA substitution, including members of Thismiaceae (Lam et al., 2018) and Burmanniaceae (Garrett et al., 2023) in Dioscoreales. Here we refer to these potentially problematic lineages as very long branch (VLB) taxa; which we define quantitatively below. It is still unclear how well VLB lineages can be accommodated in phylogenomic inference using plastid data. This is major challenge in phylogenetic inference, given the numerous losses of photosynthesis that have taken place in photosynthetic lineages across the tree of life (e.g., Merckx and Freudenstein, 2010; Figueroa-Martinez et al., 2015; Dorell at al., 2019; Nickrent, 2020; deShaw et al., 2022).

Family-level circumscription of Dioscoreales remains broadly unsettled, with three to seven families recognized in previous studies (Caddick et al., 2002a, b; Merckx et al., 2009a, 2010; Merckx and Smets, 2014; Lam et al., 2016, 2018; Givnish et al., 2018; Cheek et al., 2024). The most recent Angiosperm Phylogeny Group scheme (APG, 2016) did not update the classification of the order (recognizing only Burmanniaceae, Dioscoreaceae and Nartheciaceae), although they also acknowledged that there are problems with this *status quo*. One challenging classification issue is whether the mycoheterotrophic lineages in the order should be recognized under a single very broadly defined family (Burmanniaceae), as in recent angiosperm-wide classification schemes (APG 2003, 2009, 2016), or whether Thismiaceae should be recognized as a separate family (Merckx et al., 2009a, 2010; Hunt et al., 2014; Merckx and Smets, 2014; Lam et al., 2016, 2018; Givnish et al., 2018; Shepeleva et al., 2020). Recognizing both Thismiaceae and Burmanniaceae may also require acceptance of several photosynthetic lineages at the family level, and/or additional re-circumscription of existing families (see below). For example, fully mycoheterotrophic genus *Afrothismia* was placed in Thismiaceae until recently (Schlechter, 1921; Jonker, 1938; Maas et al., 1986; Maas-van de Kamer, 1998), but has not been included in plastid-based phylogenetic studies to date. However, few-gene phylogenetic inferences using mitochondrial and nuclear markers indicate that *Afrothismia* represents an additional independent loss of photosynthesis in Dioscoreales (Merckx and Bidartondo, 2008; Merckx et al., 2009a, 2010; Merckx and Smets, 2014; Shepeleva et al., 2020), a finding that is well supported in a recent mitochondrial phylogenomic study (Lin et al., 2022). Based on these phylogenetic findings and evidence of its morphological distinctness from Thismiaceae and related photosynthetic taxa, *Afrothismia* was recently recognized at the family level as Afrothismiaceae (Cheek et al. 2024).

The precise relationships of photosynthetic taxa within individual Dioscoreales families have also proved challenging to reconstruct. Most autotrophs have been assigned to the strictly photosynthetic families Dioscoreaceae and Nartheciaceae (e.g., APG 2003, 2009, 2016), although *Burmannia*, a member of Burmanniaceae, includes both photosynthetic and FM species. The composition of Nartheciaceae has been relatively stable in angiosperm classification (e.g., APG 2003, 2009, 2016) following the discovery that the genus *Isidrogalvia* (previously included in the family by Tamura, 1998 and Tamura et al., 2004), belongs in Alismatales, as part of Tofieldiaceae (Azuma and Tobe, 2005, 2011). However, relationships among the five genera that now comprise Nartheciaceae have varied in previous studies based on a few genes or phylogenomic studies (Merckx et al., 2008b; Fuse et al., 2012; Zhao et al., 2012; Garrett et al., 2023), and therefore need attention. The circumscription of the yam family Dioscoreaceae has also remained in flux, as two photosynthetic taxa, *Tacca* and *Trichopus*, have either been recognized as part of it (e.g., Caddick et al., 2002a,b; APG, 2003, 2009, 2016), or as two distinct monogeneric families, Taccaceae and Trichopodaceae, respectively (Huber, 1998; Kubitzki, 1998; Chase et al., 2006; Merckx et al., 2010; Merckx and Smets, 2014; Lam et al., 2016, 2018; Givnish et al., 2018; Lin et al., 2022). In addition, *Stenomeris* has been placed in Dioscoreaceae in recent angiosperm classification schemes (APG 1998, 2003, 2009, 2016), but has also been recognized in its own family, Stenomeridaceae (reviewed in Caddick et al., 2002a). Only three plastid genes are available for *Stenomeris*, which may have contributed to its inconsistent local placement in the few phylogenetic studies that have included the genus (Caddick et al., 2002a; Wilkin et al., 2005; Merckx et al., 2009a, 2010; Hsu et al., 2013; Viruel et al., 2016). A recent nuclear phylogenomic dataset recovered *Stenomeris* as sister to a clade comprising *Dioscorea, Tacca* and *Trichopus*, although with substantial gene-tree conflict for this arrangement (Zuntini et al., 2024).

Approximately 40% of the 22 genera in Dioscoreales have yet to be included in plastid phylogenomic analyses. A broadly representative plastid phylogenomic sampling would be useful to address unsettled higher-order relationships and family-level circumscriptions in the order. The degree of impact of VLB lineages on plastid-based phylogenetic inference in the order is also unclear. Here we present a plastid phylogenomic and exemplar-based survey of Dioscoreales to address its genus- and family-level relationships, with a special focus on placing its fully mycoheterotrophic lineages. We specifically address: (i) genus-level relationships in each family—our updated sampling now includes all of the photosynthetic genera and approximately half of the FM genera, or approximately three quarters of the genera; (ii) the validity of recent angiosperm classification schemes (e.g., APG 2016) in which all mycoheterotrophic taxa are lumped in Burmanniaceae; and (iii) the general impact of including VLB mycoheterotrophic lineages on phylogenomic inference, and whether strong negative effects can be ameliorated. As a backdrop to our study, we first present analyses of a “core” taxon set that includes most photosynthetic taxa and the least rapidly evolving mycoheterotrophic lineages (i.e., *Haplothismia* in Thismiaceae; *Burmannia* and *Campylosiphon* in Burmanniaceae); the latter taxa may be least affected by long-branch artefacts. We then perform a series of taxon-addition experiments that are mostly focused on different individual VLB mycoheterotrophic lineages, to reduce the chances of long-branch taxa attracting each other in phylogenetic inference (see Lam et al., 2018). We also explore several model-based approaches to attempt to reduce the effects of long-branch artefacts in phylogenomic inference.

## MATERIALS AND METHODS

### Taxon sampling and library preparation

We used genome skims (Straub et al., 2012) to generate a new phylogenomic dataset comprising the full complement of plastid protein-coding genes and rDNA genes for seven Dioscoreales species (Appendix S1; see Supplemental Data with this article). We added these data to published sequences for 29 other taxa in the order, along with 62 additional monocots as outgroups, to obtain a sampling that includes representatives of all 12 monocot orders (sensu Givnish et al., 2018; see Appendix S1 for source information for all sampled taxa). The 36 taxa sampled in Dioscoreales represent all nine strictly photosynthetic genera in the order and seven of 13 mycoheterotrophic genera for which we were able to retrieve plastid data. Six of the seven latter genera are fully mycoheterotrophic (FM), and *Burmannia* includes both photosynthetic and FM taxa (Table 1). We were unable to obtain sequenceable DNA for photosynthetic *Stenomeris* (which is currently placed in Dioscoreaceae; APG 1998, 2003, 2009, 2016), despite numerous attempts using a range of herbarium and fixed material, and so we represented this genus with three available plastid genes obtained from GenBank (Appendix S1). Genome skims for nearly all newly sequenced Dioscoreales taxa were produced following approaches outlined in Lam et al. (2015), except that we prepared genomic libraries using two additional library preparation kits (Bioo NEXTflex Rapid DNA Sequencing Kit, Bioo Scientific Corp., Austin, Texas, USA and NEBNext Ultra II DNA Library Prep Kit, New England Biolabs, Ipswich, Massachusetts, USA). We sequenced these libraries as 100-bp, 125-bp or 300-bp paired-end reads respectively on multiplexed lanes of HiSeq 2000, HiSeq 2500 or MiSeq platforms (Illumina, San Diego, California, USA). We obtained plastid data for *Trichopus sempervirens* as genome-skim “bycatch” from a nuclear target enrichment gene panel (Soto Gomez et al., 2019) following approaches outlined in Viruel et al. (2019), which we sequenced as 150-bp paired-end reads in a multiplexed lane of a HiSeq X platform.

**Table 1.** Nutritional status and approximate number of species in the 22 genera that comprise Dioscoreales. Sampled genera are shown in bold font. Note that *Burmannia* (*) includes both photosynthetic and fully mycoheterotrophic species, and that monotypic *Marthella* (†) may be extinct.

| Genus | Family | Nutritional status |  | Number of species |  |
| --- | --- | --- | --- | --- | --- |
|  |  | Photosynthetic | Fully<br>mycoheterotrophic | Sampled | Total |
| *Afrothismia* | Afrothismiaceae |  | X | 4 | 12<sup>a</sup> |
| *Apteria* | Burmanniaceae |  | X | 1 | 1<sup>a</sup> |
| *Burmannia* * | Burmanniaceae | X | X | 5 | 56<sup>a</sup> |
| *Campylosiphon* | Burmanniaceae |  | X | 1 | 2<sup>a</sup> |
| *Dictyostega* | Burmanniaceae |  | X | N/A | 1<sup>a</sup> |
| *Gymnosiphon* | Burmanniaceae |  | X | 1 | 32<sup>a</sup> |
| *Hexapterella* | Burmanniaceae |  | X | N/A | 2<sup>a</sup> |
| *Marthella*† | Burmanniaceae |  | X | N/A | 1<sup>a</sup> |
| *Miersiella* | Burmanniaceae |  | X | N/A | 1<sup>a</sup> |
| *Dioscorea* | Dioscoreaceae | X |  | 7 | 629<sup>b</sup> |
| *Stenomeris* | Dioscoreaceae | X |  | 1 | 2<sup>c</sup> |
| *Aletris* | Nartheciaceae | X |  | 3 | 24<sup>d</sup> |
| *Lophiola* | Nartheciaceae | X |  | 1 | 1<sup>d</sup> |
| *Metanartheicum* | Nartheciaceae | X |  | 1 | 1<sup>d</sup> |
| *Nartheicum* | Nartheciaceae | X |  | 1 | 8<sup>d</sup> |
| *Nietneria* | Nartheciaceae | X |  | 1 | 2<sup>d</sup> |
| *Tacca* | Taccaceae | X |  | 2 | 17<sup>b</sup> |
| *Haplothismia* | Thismiaceae |  | X | 1 | 1<sup>a</sup> |
| *Oxygyne* | Thismiaceae |  | X | N/A | 6<sup>e</sup> |
| *Thismia* | Thismiaceae |  | X | 2 | 45<sup>a</sup> |
| *Tiputinia* | Thismiaceae |  | X | N/A | 1<sup>a</sup> |
| *Trichopus* | Trichopodaceae | X |  | 2 | 2<sup>b</sup> |

Notes: <sup>a</sup>Merckx et al. (2013a); <sup>b</sup>Govaerts et al. (2021); <sup>c</sup>Caddick and Wilkin (1998); <sup>d</sup>Govaerts (2021); <sup>e</sup>Cheek et al. (2018).

### Contig assembly and gene annotation

We used methods described in Lam et al. (2015) to assemble contigs from genome-skim data, and to retrieve sets of plastid protein-coding and rDNA genes after filtering for possible mitochondrial and nuclear contigs, with minor modifications noted here. We used CLC Genomics Workbench (v.6.5.1; CLC Bio, Aarhus, Denmark) to assemble contigs for all but one taxon (*Trichopus sempervirens*, represented here using genome-skim “bycatch” data), for which we used NOVOPlasty v. 2.7.2 (Dierckxsens et al., 2017) with the *rbc*L plastid gene sequence of *Dioscorea elephantipes* (GenBank accession NC_009601.1) as a seed input, and otherwise default settings. All contigs assembled here have at least 30× average coverage regardless of assembly program. We identified plastid gene sets from *T. sempervirens* using GeSeq (Tillich et al., 2017) with default settings. For FM taxa with substantially elevated rates (i.e., *Afrothismia* sp.*, A. gesnerioides, A. hydra, A. winkleri, Apteria aphylla, Gymnosiphon longistylus, Haplothismia exannulata, Thismia rodwayi* and *T. tentaculata*), we additionally ran pairwise BLASTP searches (Altschul et al., 1990) on individual genes to retain regions with an expect (e)-value threshold of 10 relative to homologous genes of photosynthetic taxa. We used genes from photosynthetic *Tacca leontopetaloides* as a query sequence for searches of mycoheterotrophic *Afrothismia, Haplothismia* and *Thismia*, and used photosynthetic *Burmannia capitata* as a query for mycoheterotrophic *Apteria* and *Gymnosiphon*. Most of the recovered protein-coding genes for *Afrothismia* sp., *A. hydra* and *A. winkleri* contain internal stop codons at one to eight different positions across individual genes, and were slightly or moderately truncated compared to other sampled taxa. These internal stop codons may be undocumented RNA-edited sites (e.g., Takenaka et al., 2013; Bell et al., 2020) or instances of pseudogenization. By contrast, the retained genes of *A. gesnerioides* were completely uninterrupted, although most were either truncated or slightly longer relative to other sampled taxa. We included all of the *Afrothismia* sequences in phylogenetic analyses described below.

### Sequence alignment and concatenated matrix construction

We assembled a 98-taxon matrix by adding species that were newly sequenced here to published data retrieved from GenBank and two published matrices (Lam et al., 2018; Garrett et al., 2023). To do so, we generated 82 individual plastid gene files, representing 78 protein-coding genes and four rDNA genes, each with 98 taxon terminals. We produced DNA-based alignments for each gene, including manual adjustments, and prepared a concatenated matrix as described in Soto Gomez et al. (2020) for the plastid dataset in that study. To help with manual adjustments for aligning FM taxa, we additionally used the results from pairwise BLASTP searches described above. We applied procedures used by Lam et al. (2015) to check for matrix compilation errors; none were found. Photosynthetic taxa are represented by all or most genes found in plastid genomes of photosynthetic angiosperms, except *Stenomeris borneensis,* for which we only included three available protein-coding genes (i.e., *atp*B, *mat*K, *rbc*L; Appendix S1). Fully mycoheterotrophic taxa are represented by 6–24 protein-coding genes (counting putative pseudogenes in *Afrothismia*) and 2–4 rDNA genes, due to gene loss (Fig. 1). We coded missing genes (i.e., those lost in FM taxa or the few that were not recovered for photosynthetic taxa) as missing data. The gene sets retrieved for each of the FM taxa are likely complete or nearly so, based on preliminary plastid genome assemblies to be presented elsewhere. We retrieved the *ycf*1 locus for most photosynthetic taxa and for the FM species *Campylosiphon congestus* and *Burmannia itoana*, but did not include it in the final matrix due to difficulties aligning it across the full taxon set. The final DNA-based matrix is 80,390-bp long (derived from 67,156 bp of unaligned plastid sequence data in *Dioscorea membranacea*, for reference). We also translated this into a matrix comprising 24,980 amino-acid residues. Data matrices are available on figshare (https://figshare.com/s/8470257673242e6bf43f), and see Appendix S1 for GenBank numbers.

**Figure 1.**
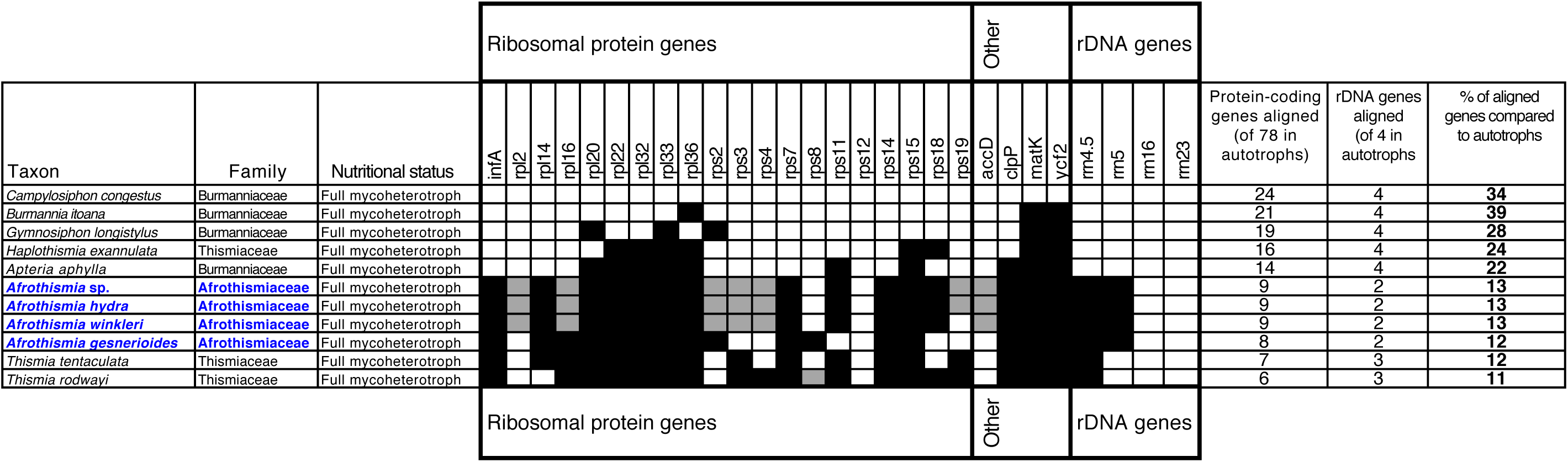
Fully mycoheterotrophic Dioscoreales sampled retain only a subset of plastid genes, summarized for loci included in alignments here (i.e., excluding *ycf*1 and tRNA genes). Species are ordered vertically by number of genes retained, and genes horizontally by functional complex; gene nomenclature follows Wicke et al., (2011). Newly sequenced taxa are shown in blue. White box = gene present, black = gene lost (or possibly not retrieved from taxa with preliminary plastid genome assemblies), grey = genes containing internal stop codons. Note that *ycf*1 was recovered in the Burmanniaceae species *C. congestus* and *B. itoana* (Lam, 2016) but not in Thismiaceae (Garrett et al., 2023) or Afrothismiaceae. A range of tRNA genes (excluding pseudogenized copies) were recovered from Burmanniaceae species (*C. congestus*: 23, *B. itoana*: 11, *G. longistylis*: 5, *A. aphylla*: 3; Lam, 2016), Thismiaceae species (*H. exannulata*: 7, *T. tentaculata*: 2, *T. rodwayi*: 1; Garrett et al., 2023) and Afrothismiaceae species (*Afrothismia* sp., *A. hydra*, *A. winkleri*, *A. gesnerioides*: 1).

### Characterization of substitution rate differences in Dioscoreales

We characterized relative differences in the substitution rates of mycoheterotrophic and photosynthetic taxa using a Bayesian framework implemented in BEAST v. 2.6.3 (Bouckaert et al., 2019). We prepared a reduced dataset to keep these computationally demanding analyses tractable in terms of the number of genes and taxa. We selected seven plastid genes (*acc*D, *rpl*2, *rps*12, *rps*4, *rps*8, *rrn*16, *rrn*23) that were available for a broad taxon sampling, which resulted in a 62-taxon dataset that includes at least one representative of each of the sampled mycoheterotrophic genera (Fig. 1). To focus on characterizing rate variation, we constructed an ultrametric input tree with a fixed topology based on the best likelihood tree recovered in the mitochondrial data inferences of Lin et al. (2022). Our analysis focuses on relative rates (e.g., Lam et al. 2018), and so we did not specify fossil constraints or attempt to date individual nodes, which may lead to artefactual results when highly rate-elevated taxa are included (e.g., Iles et al., 2015; Soto Gomez et al., 2020). We specified a GTR nucleotide substitution model with four categories, estimated substitution rates and empirical frequencies under a birth death model and a random local clock (Drummond and Suchard, 2010); we otherwise used default settings. We ran six independent analyses, each for 400 million generations, sampling every 2000 trees, and visually assessed stationarity and convergence amongst them using Tracer v. 1.7.1 (Rambaut et al., 2018). We considered posterior ESS values >200 to be acceptable, which was the case for all parameters. We used the LogCombiner program in BEAST to combine five analyses that converged, and the TreeAnnotator program to summarize the resulting sampling of trees after discarding 20% as burn-in and resampling at a lower frequency to obtain 10,000 trees that were used to make a single combined tree.

### Partitioned likelihood analyses

We conducted most phylogenetic analyses using a maximum likelihood approach with partitioned data as implemented in RAxML-NG (Kozlov et al., 2019). We ran multiple partitioned analyses using variant versions of the data matrix. However, the first two of these analyses were based on a core 89-taxon version of the matrix that includes (a) most photosynthetic taxa in Dioscoreales, including photosynthetic taxa in Burmanniaceae (i.e., *B. bicolor*, *B. capitata, B. coelestis* and *B. disticha*) and (b) three of the least rapidly evolving FM taxa (i.e., *Burmannia itoana* and *Campylosiphon congestus* in Burmanniaceae, and *Haplothismia exannulata* in Thismiaceae). Thus, this 89-taxon core taxon set <u>excludes</u> all VLB taxa (see below). It also excludes *Stenomeris borneensis*, a data-deficient photosynthetic taxon. We selected the three least rapidly evolving FM taxa after assessing rate elevation in an analysis of rate variation (see Fig. 2 and below) and by visual inspection of branch-length estimates in likelihood analyses here. We ran partitioned DNA- and AA-based analyses on this core 89-taxon set, and used the resulting trees as references to examine the effect of nine potentially problematic taxa on phylogenetic inference.

**Figure 2.**
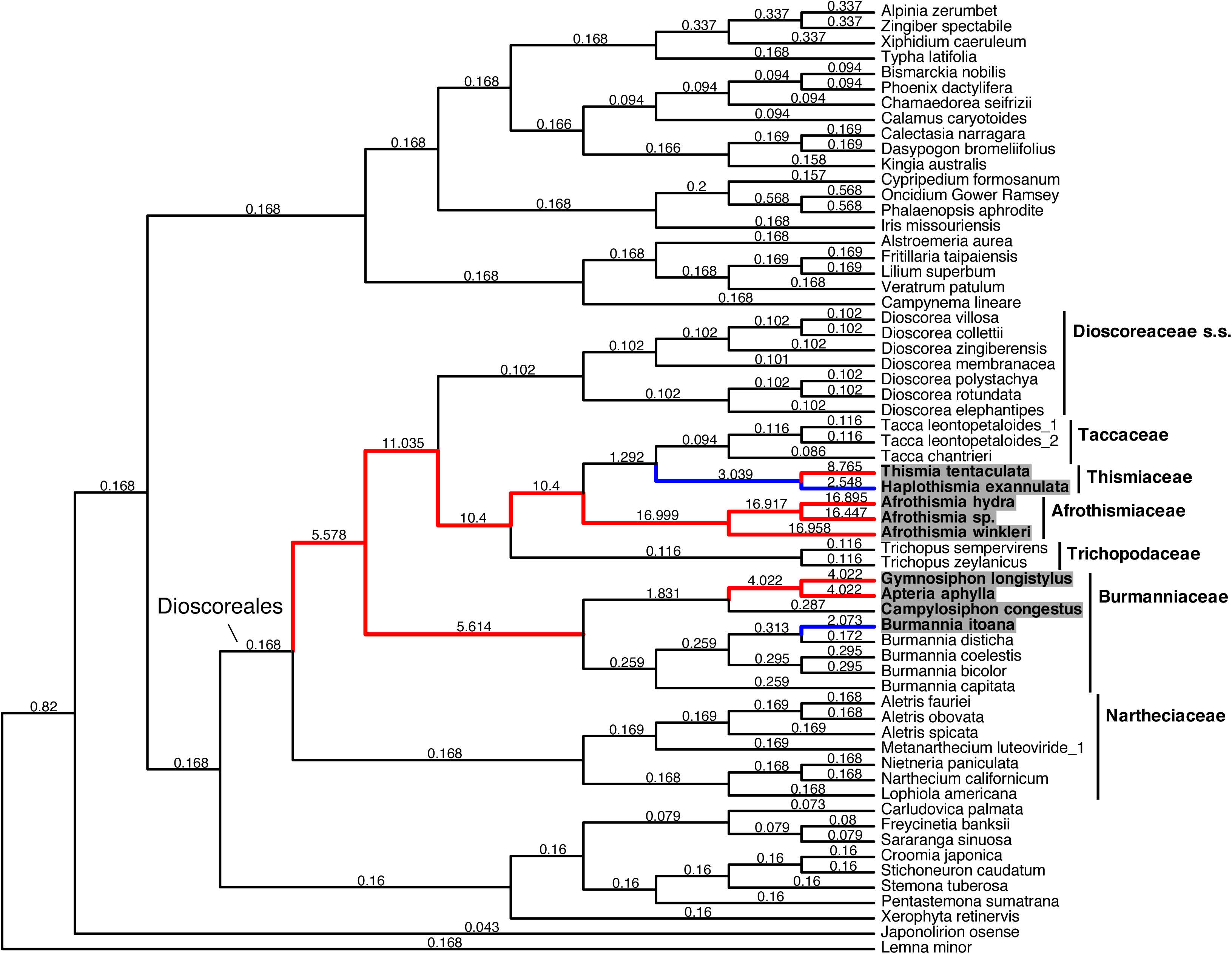
Fully mycoheterotrophic members of Dioscoreales display a range of elevated DNA substitutional rates in plastid genes, as determined in a Bayesian analyses of relative rates with a random local clock model, for a reduced dataset of monocots (seven plastid genes available for 62 taxa, using a fixed topology based on Fig. 4 in Lin et al., 2022). The estimated number of substitutions per site is shown for individual branches. Blue branches indicate intermediate rates that are two–four times faster; red branches indicate those that are over four times faster. We refer to the latter as very long branch (VLB) taxa here.

Eight of the nine problematic taxa are highly rate-elevated mycoheterotrophs in Afrothismiaceae, Burmanniaceae and Thismiaceae (see below); the ninth is the data-deficient photosynthetic taxon (*Stenomeris*), for which only three plastid genes are available. We assessed the placement of the nine potentially problematic taxa by adding one or more of these excluded species to the core 89-taxon dataset (in separate DNA-based partitioned likelihood analyses). The taxon changes to the core 89-taxon matrix included adding only (independently), in turn: (i) the data-deficient photosynthetic taxon, *S. borneensis*, resulting in a 90-taxon matrix; (ii) two VLB representatives of Burmanniaceae, *Apteria aphylla* and *Gymnosiphon longistylus*, resulting in a 91-taxon matrix; (iii) two VLB representatives of Thismiaceae, *Thismia rodwayi* and *T. tentaculata*, resulting in a second 91-taxon matrix; (iv) four VLB species of Afrothismiaceae—by far the most rate-elevated lineage that we surveyed—resulting in a 93-taxon matrix; (v) the only sampled member of Afrothismiaceae lacking internal stop codons in protein-coding genes, *A. gesnerioides*, resulting in a second 90-taxon matrix; and (vi) all Dioscoreales taxa with molecular data here, resulting in a “full” taxon set comprising 98 species. We additionally analyzed the full 98-taxon matrix in a partitioned analysis based on amino-acid data.

For these analyses, we partitioned DNA alignments by considering gene and codon position (a “G x C” scheme, e.g., Lam et al., 2015) for the 78 protein-coding genes, with additional starting partitions for each of the four rDNA genes, resulting in 238 initial partitions. The amino-acid (AA) matrix had 78 initial partitions, corresponding to one partition per protein-coding gene. We used the ModelFinder function (Kalyaanamoorthy et al., 2017) in IQ-TREE v. 1.6.12 (Nguyen et al., 2015) to combine partitions that did not have substantially different substitution models, employing the relaxed hierarchical clustering algorithm to examine the top 10% partition merging schemes (-rcluster 10), the corrected Akaike information criterion (AICc) to merge partitions, and considered invariant (I) and gamma (G) rate heterogeneity across sites (-m TESTMERGEONLY). We obtained a range of optimal DNA and AA substitution models for the resulting partitioning schemes (Appendix S2), which we used in subsequent likelihood analyses.

The DNA- and AA-based partitioned likelihood analyses performed using RAxML-NG included 40 independent searches for the best tree, each search starting from 20 parsimony-based and 20 random trees. We estimated branch support using bootstrap analysis (Felsenstein, 1985) by applying the extended majority rule (MRE) bootstrap stopping test (Pattengale et al., 2010) every 50 replicates. This test consists of randomly splitting the set of bootstrap replicates into two equal halves (with 1000 permutations per test), computing the majority rule consensus trees for each half, and using the weighted Robinson-Foulds (WRF) criterion to assess dissimilarity between the two resulting consensus trees. Bootstrapping is stopped when the relative WRF between sets is ≤1% for at least 990 permutations (--bs-cutoff 0.01). Here this resulted in 100–1000 standard/slow bootstrap replicates across the eight variant likelihood analyses. Each bootstrap analysis used the same partitioning schemes and substitution models as the corresponding searches for best trees. For all bootstrap analyses performed here, we considered weakly and strongly supported branches to respectively have <70% and ≥95% support (e.g., Soltis and Soltis, 2003).

### Phylogenetic analyses using site-heterogeneous models

We also analysed AA data using two model-based approaches thought to attempt to ameliorate the effects of long-branch artefacts (Lartillot et al., 2007; Quang et al., 2008; Wang et al., 2018). We implemented these using IQ-TREE v. 1.6.12 (Nguyen et al., 2015) for likelihood inference, and PhyloBayes MPI v. 1.9 (Lartillot et al., 2013) for Bayesian inference. For IQ-TREE analyses, we used a posterior mean site frequency (PMSF) model with the most complex empirical mixture model implemented in that program, which includes 60 AA mixture classes, empirical state frequencies and a discrete gamma distribution with four rate categories (-m LG+C60+F+G). In PMSF analyses, AA profiles are estimated for each site in an alignment based on a guide tree (estimated here using a simpler model in IQ-TREE, -m LG+F+G) and an input mixture model (estimated in the first phase of the analysis). The PMSF model has been shown to ameliorate the effects of long-branch artefacts in simulated and empirical phylogenomic analyses (Wang et al., 2018), while providing a rapid approximation of the more computationally intensive profile mixture models (Quang et al., 2008). We conducted 1000 ultrafast bootstrap replicates to estimate branch support for this approach.

For AA-based analyses using PhyloBayes MPI, we implemented the CAT-GTR model (using a discretized gamma distribution with four categories), which captures variation in AA preferences across sites using a Bayesian non-parametric approach (Lartillot, 2020). We ran two independent chains until they reached ∼10,200 points (i.e., iterations) and quantitatively assessed their convergence and mixing using the tracecomp and bpcomp commands, respectively, after setting a burn-in of 1000. We considered ESS values >100 and discrepancies among chains <0.3 to represent acceptable runs (Lartillot, 2020), which was the case for all model parameters. We obtained a consensus tree with posterior probabilities for branch support by pooling the trees produced by the two chains.

### Constraint tests of alternative phylogenetic hypotheses

We performed Shimodaira-Hasegawa (SH) and approximately unbiased (AU) tests (Shimodaira and Hasegawa, 2001) to assess variant placements for VLB mycoheterotrophs. Specifically, we tested whether several alternative arrangements of these mycoheterotrophs were significantly worse than optimal trees from analysis of the core 89-taxon set (i.e., alternative hypothesis i below; cf. Fig. 3, Appendices S3–S5), or from analysis of an expanded 93-taxon set that included Afrothismiaceae (i.e., alternative hypotheses ii–v below; cf. Appendix S6 and Appendices S7–S10). The minimal constraints used in each test are indicated with red branches in Appendices S7–S10. We used family-level boundaries based on narrow-sense circumscriptions (as shown in Table 1, following Merckx et al., 2010; Givnish et al., 2018; Lam et al., 2018; Cheek et al. 2024), and we follow these throughout the study, except where noted. We included *Haplothismia* as a slowly evolving mycoheterotroph to represent Thismiaceae in these tests, and treated Afrothismiaceae as a lineage distinct from the rest of Thismiaceae, when included (following Cheek et al., 2024). *Stenomeris*, a photosynthetic taxon with uncertain family placement that was represented here by only a few genes, was excluded from all constrained analyses.

**Figure 3.**
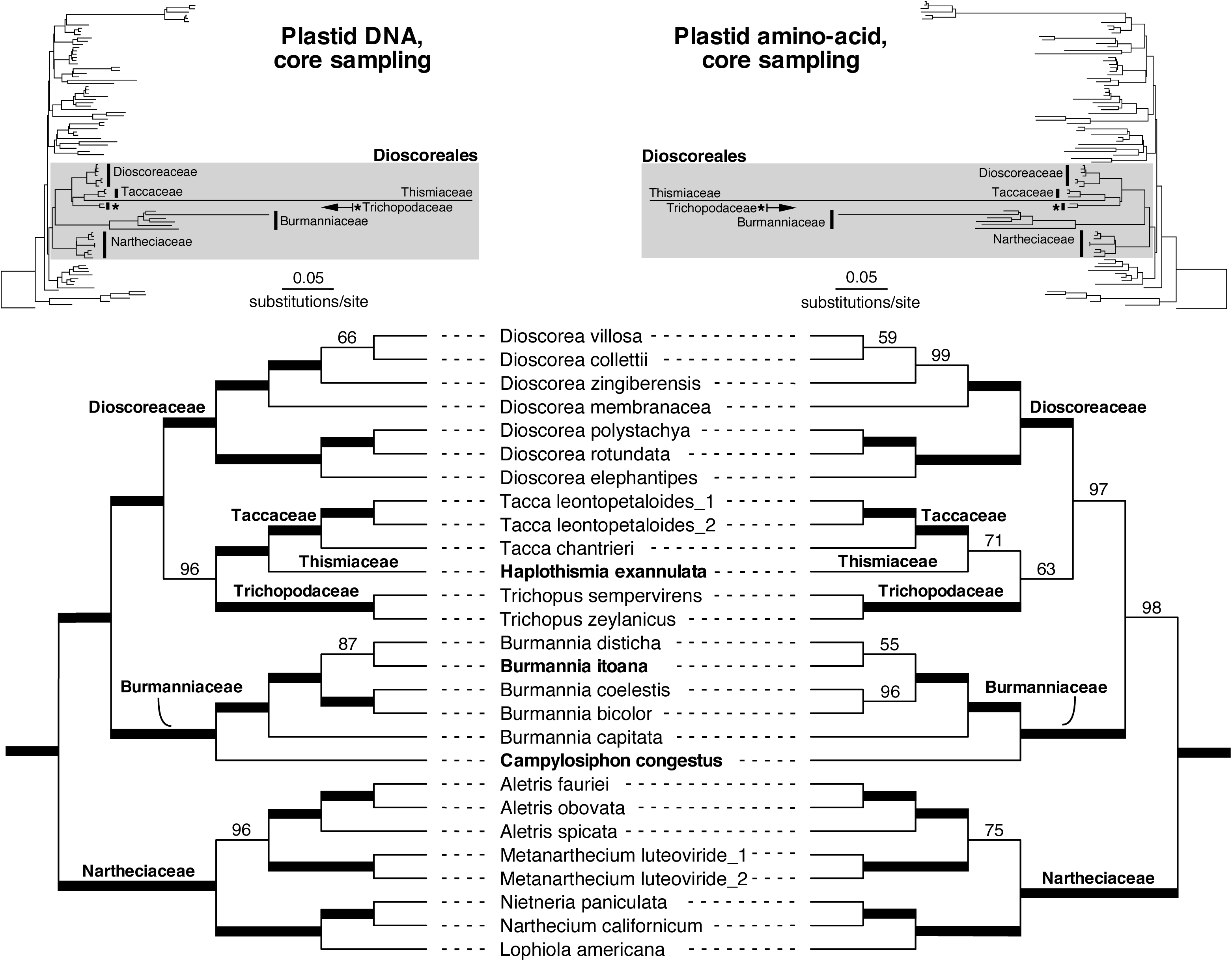
Highly congruent phylogenetic estimates recovered for Dioscoreales based on DNA vs. amino acid (AA)-based likelihood analyses of a “core” 89-taxon set (see text). Three relatively slowly evolving fully mycoheterotrophic taxa in Burmanniaceae and Thismiaceae were included in this core set (noted here in bold font; and see relative rates depicted in Fig. 2). Both analyses exclude very long branch (VLB) mycoheterotrophs and a data-deficient photosynthetic species (see text for details of genes included, and Appendices S3 and S4 for the full trees). Thick lines indicate 100% bootstrap support; values <100% are shown above branches. Upper phylograms depict full samplings of each analysis, with Dioscoreales shaded in grey; scale bars indicate estimated substitutions per site.

For alternative hypothesis (i), we constrained all members of Burmanniaceae and Thismiaceae to form a clade (we excluded Afrothismiaceae from consideration in this case, to remove the possible distorting effect of its extreme rate elevation, e.g., Appendices S6, S11). Thus, ignoring Afrothismiaceae, this constraint is consistent with the current broad circumscription of Burmanniaceae in recent angiosperm classifications that incorporates Thismiaceae (APG, 2003, 2009, 2016). We compared the resulting tree to a null hypothesis from the corresponding unconstrained search, in which Burmanniaceae and Thismiaceae are inferred to be distantly related clades in Dioscoreales (see Fig. 3, Appendices S3, S4). The remaining tests consider possible placements of Afrothismiaceae, by constraining clades that comprise: (ii) Afrothismiaceae and Dioscoreaceae; (iii) Afrothismiaceae and Burmanniaceae, the latter narrowly defined; (iv) Afrothismiaceae and Thismiaceae (= Thismiaceae s.l.); and (v) Afrothismiaceae, Thismiaceae and Taccaceae, with the latter two families forced to be sister taxa as in Lin et al. (2022). Hypotheses ii–v each assess different possible arrangements of Afrothismiaceae in Dioscoreales that we observed in phylogenetic analyses here (i.e., hypotheses ii–iii; see Fig. 4, Appendices S11–S14), or that are consistent with a possible broad circumscription of Thismiaceae to include Afrothismiaceae (i.e., hypothesis iv; see Maas-van de Kamer, 1998), or with a possible placement of Afrothismiaceae as a close relative of Thismiaceae and Taccaceae (i.e., hypothesis v; see Merckx et al., 2009a, 2010; Lin, 2022). We compared these four alternative hypotheses to a main hypothesis recovered in phylogenetic analyses here, where Afrothismiaceae forms a clade with the family Poaceae in the order Poales (see Appendix S6).

**Figure 4.**
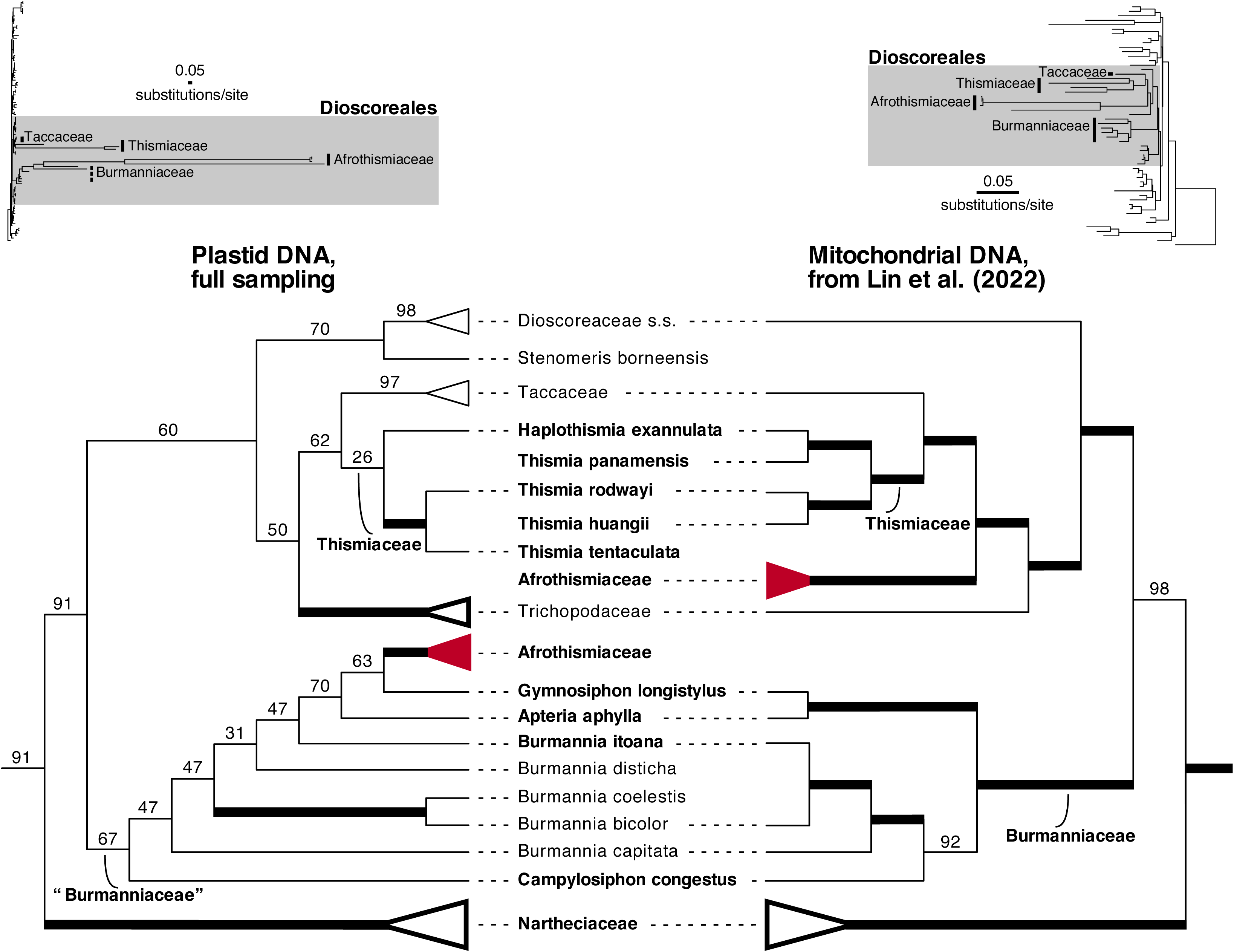
Highly divergent placements of Afrothismiaceae (red clade) in plastid-vs. mitochondrial phylogenomic analysis (left-vs. right-hand trees). Both trees were inferred using partitioned DNA-based maximum-likelihood analyses. The plastid tree was inferred using the “full” taxon sampling of Dioscoreales, which included very long branch (VLB) mycoheterotrophs and a data-deficient species (*Stenomeris*), based on 82 plastid genes (7–28 genes for fully mycoheterotrophic taxa, in bold font). The mitochondrial tree was inferred by Lin et al. (2022; Fig. 4 there) based on 37 mitochondrial genes. Nartheciaceae and all monogeneric families represented by more than one species (Afrothismiaceae, Dioscoreaceae, Taccaceae, Trichopodaceae) are depicted here as triangles. Thick lines indicate 100% bootstrap support; values <100% are shown above branches. The upper phylograms depict monocot phylograms for each analysis, with Dioscoreales shaded in grey, and non-monophyletic families indicated with dotted lines (see Appendix S12 here and Fig. 4 in Lin et al., 2022 for further detail). Scale bars indicate estimated substitutions per site.

In all cases, we tested whether we could reject trees recovered under constrained hypotheses compared to the unconstrained optimal trees for that taxon set. For tree searches using constrained branches, we excluded data-deficient and VLB taxa that were not relevant to the hypothesis. Thus, we employed the core 89-taxon set for test (i), and a modified version of the 93-taxon set including only two species of Afrothismiaceae for tests (ii)–(v). We did the latter reduction because three of the four Afrothismiaceae taxa are nearly identical at the sequence level. To find trees consistent with these five hypotheses, we made constraint trees for tests (i)–(iv) by constraining one branch to obtain two taxon partitions, one comprising the species of interest, and the other comprising the remaining taxa (highlighted branches in Appendices S5, S7–S9). For test (v) we constrained two branches to simultaneously obtain two taxon partitions, one comprising Thismiaceae-Taccaceae, and the other comprising Afrothismiaceae-Thismiaceae-Taccaceae (in this case effectively placing Afrothismiaceae close to the other two families, but not allowing it to be inside or sister to either family individually; highlighted branches in Appendix S10). We recovered the best likelihood trees that satisfied each of the five constraints using the -tree-constraint function in RAxML-NG, with default settings. We then performed the SH and AU tests in CONSEL v.0.20 (Shimodaira and Hasegawa, 2001), using site-likelihoods from partitioned likelihood analyses, to address whether the fit of the data was significantly different for unconstrained vs. constrained best trees.

## RESULTS

### Rate elevation in fully mycoheterotrophic lineages of Dioscoreales

The Bayesian analysis characterizing relative substitution rates found moderate to very high levels of rate elevation in the fully mycoheterotrophic (FM) taxa compared to photosynthetic relatives (Fig. 2; grey highlights indicate nine fully mycoheterotrophic taxa). Our initial phylogenetic analysis included three of nine tested FM species in the core taxon sampling (Fig. 3: *Campylosiphon congestus* and *Burmannia itoana* in Burmanniaceae, and *Haplothismia exannulata* in Thismiaceae). These three FM taxa exhibited moderate substitutional elevation, which we defined as rates up to around two to four times higher in their corresponding terminal lineages (or on nearby subtending branches) compared to photosynthetic relatives (Fig. 2). The six remaining mycoheterotrophs, which we define as very long branch taxa, all have substantially higher substitution rate elevation, with the very highest rates (>16-fold) in Afrothismiaceae taxa (Fig. 2). Curiously, several internal branches with high rates are inferred to occur near the origin of Dioscoreales. This may be because the algorithm wrongly inferred shared rate elevation in the common ancestors of lineages that independently evolved high rates; if so, this may imply that rates in some terminal branches are underestimated in the rates analysis (Fig. 2, and see also the plastid-based phylograms in Figs. 3, 4, Appendices S6, S11–S16). Regardless, based on the high rate elevation inferred here for multiple VLB taxa in Afrothismiaceae, Thismiaceae and Burmanniaceae (Fig. 2), we implemented taxon-addition phylogenetic experiments to sequentially (and independently) add these lineages to the core taxon set comprising photosynthetic species and more slowly evolving mycoheterotrophs (Appendices S6, S11, S15, S16).

### Phylogenetic analyses of the core taxon set (excluding VLB taxa and Stenomeris)

The trees inferred from DNA- and AA-based likelihood analyses of the core 89-taxon dataset are congruent with each other and are also generally very well supported regarding higher-level relationships in Dioscoreales (Fig. 3; see Appendices S3 and S4 for monocot-wide relationships from DNA and AA inferences, respectively). Well-supported relationships, which are consistent between DNA and AA analyses, include the placements of the three most slowly evolving FM lineages from Burmanniaceae (*Burmannia itoana* and *Campylosiphon congestus*) and Thismiaceae (*Haplothismia exannulata*) (Fig. 2). The FM family Thismiaceae—represented by *Haplothismia* alone— is part of a clade with Taccaceae, with both families in turn sister to Trichopodaceae among sampled taxa (Fig. 3, Appendices S3, S4). This arrangement has strong support from DNA data and moderate support from AA data: these three families (Taccaceae, Thismiaceae and Trichopodaceae) are strongly supported as part of a larger clade with Dioscoreaceae in both analyses. Thus, the family Thismiaceae is not associated with Burmanniaceae, and there is strong bootstrap support for their separation.

Burmanniaceae, represented in the core set by two FM and four photosynthetic taxa, is instead sister to all families of Dioscoreales except Nartheciaceae (Fig. 3, Appendices S3, S4). These family-level relationships in Dioscoreales all have moderate or strong support from DNA and AA analyses, as do most genus-level relationships (Fig. 3, Appendices S3, S4). Outside Dioscoreales, the monocot-wide relationships are also both congruent and generally well supported across analyses of the core taxon set (Appendices S3, S4). For example, Dioscoreales is consistently strongly supported as the sister group of Pandanales, with 100% bootstrap support for this relationship (Appendices S3, S4).

### Effect of including a data-deficient photosynthetic species

The placement of photosynthetic *Stenomeris borneensis* is consistent across analyses that included this species. It is represented here by only three available plastid genes from GenBank (Fig. 5, Appendices S12–S14, S17). We recovered *Stenomeris* as the sister group of *Dioscorea* (Dioscoreaceae) with poor to moderate support in DNA analyses when added individually to the core taxon set, or when included as part of the full 98-taxon set (67– 70%; Fig. 5, right-hand tree, and Appendices S12, S17). There was at best moderate support for its placement in AA analyses of the full taxon set using site heterogeneous models in both likelihood and Bayesian frameworks (80–81%; Appendices S13, S14).

**Figure 5.**
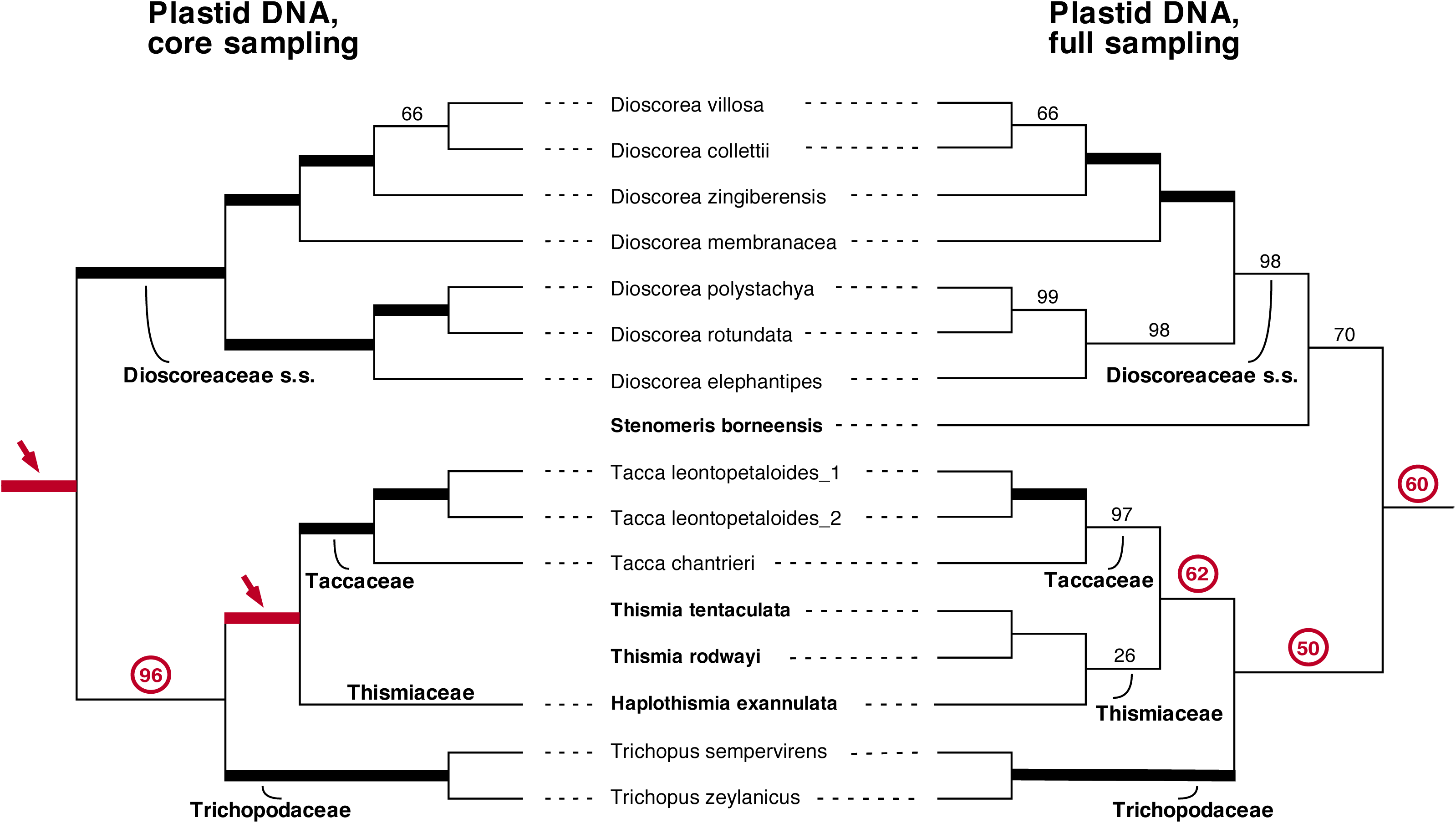
Strong depression of bootstrap support values, with consistent placement of Thismiaceae, in DNA-based maximum likelihood analysis, following inclusion of very long branch (VLB) mycoheterotrophs in Dioscoreales. Relationships were inferred in partitioned likelihood analyses of plastid DNA using either the core (89-taxon) vs. full (98-taxon) taxon samplings (left-vs. right-hand trees, respectively). Support values are indicated on branches (thick lines indicate 100% bootstrap support); red branches/arrows/circles indicate branches that are well supported in analysis of the core taxon set that have a substantial drop in support in the analysis of the full taxon set. Mycoheterotrophic taxa and the data-deficient photosynthetic taxon *Stenomeris* are shown in bold font. The full-taxon analysis includes all VLB mycoheterotrophs and *Stenomeris*, in addition to the taxa included in the core taxon set (see text). Two Dioscoreales families Burmanniaceae and Nartheciaceae are omitted for clarity, but were included in the analyses (see Appendices S3 and S12 for full relationships, and Fig. 3 for a view of the core taxon sample across Dioscoreales).

### Taxon-addition experiments with VLB mycoheterotrophic lineages

Adding individual (or related sets of) VLB mycoheterotrophs in Burmanniaceae and Thismiaceae to analyses yielded generally congruent relationships in Dioscoreales and across monocots, relative to the core taxon set (cf. Fig. 3, Appendices S3, S4 and Appendices S15, S16). Despite this, most branches neighboring points of attachment of VLB lineages to the tree—and often more distant ones—experienced reductions in bootstrap support compared to the DNA-based analysis of the core taxon set (see Table 2, cf. Fig. 3 and Appendices S15, S16). In a taxon-addition experiment in which only the two VLB mycoheterotrophs in Burmanniaceae (*Apteria* and *Gymnosiphon*) are added to the core set, these two taxa form a strongly supported clade that is in turn weakly placed as the sister group of *Campylosiphon* (52% support; Table 2, Appendix S15). Three of the five remaining branches in Burmanniaceae then exhibit substantial depression in bootstrap support of 41–47%, compared to DNA-based analysis of the core taxon set (Table 2, cf. Fig. 3 and Appendix S15). In a taxon-addition experiment in which only the two VLB mycoheterotrophs in Thismiaceae (*Thismia rodwayi* and *T. tentaculata*) are added to the core set, both *Thismia* and *Haplothismia* are inferred to be closely related to *Tacca* in a poorly supported grade (rather than a clade), with 44% bootstrap support (Table 2, Appendix S16). The addition of these two species did not depress branch support substantially (Table 2, cf. Fig. 3 and Appendix S16).

**Table 2.**
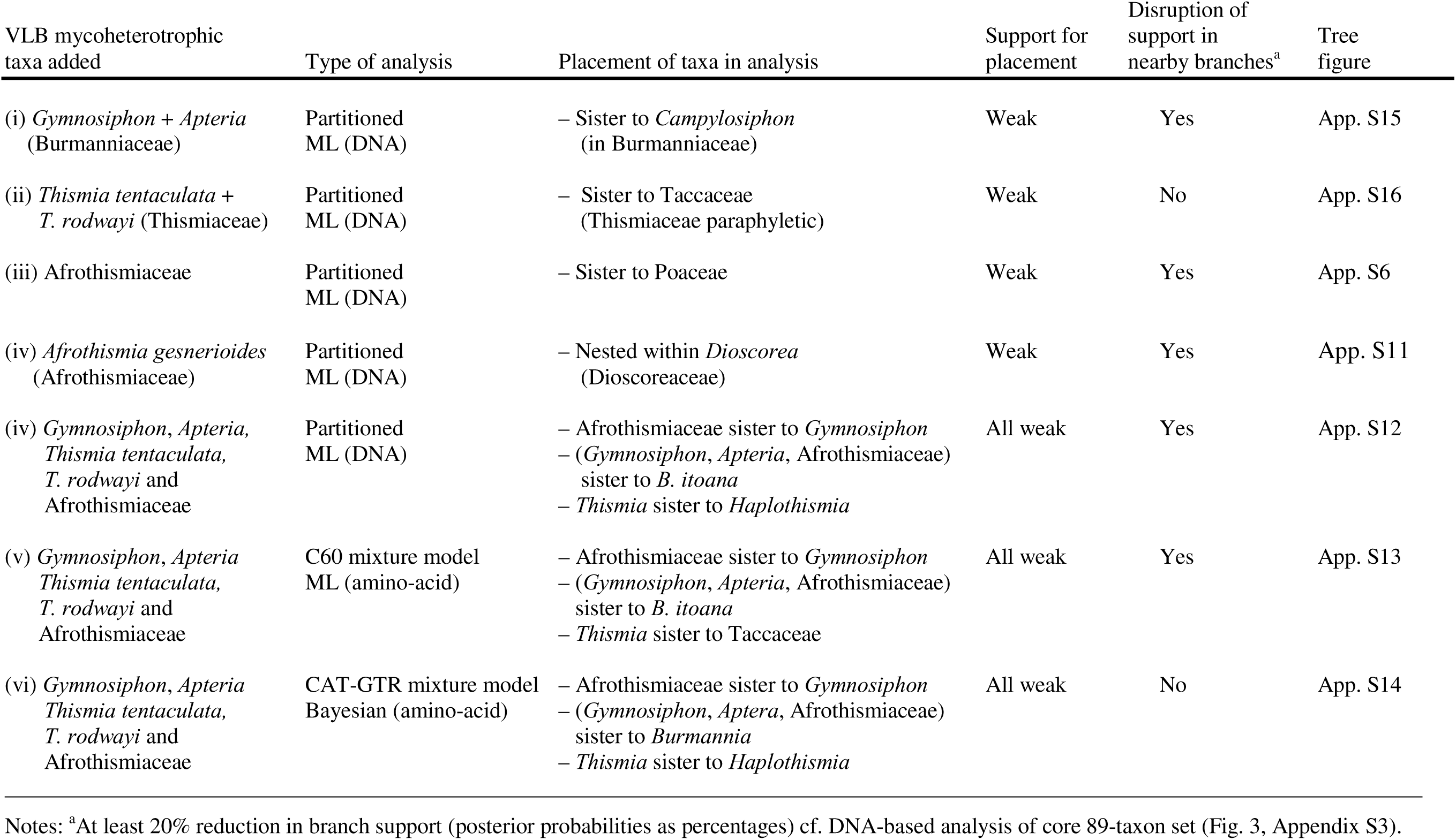
Summary of phylogenetic experiments to sequentially add very long branch (VLB) mycoheterotrophs to a core 89-taxon set which includes only moderately rapidly evolving mycoheterotrophs (i.e., *Haplothismia* in Thismiaceae, *Burmannia* and *Campylosiphon* in Burmanniaceae). The level of branch support for the placement of added taxa is indicated with respect to phylogenetic inferences documented in individual Appendices (“App.”).

| VLB mycoheterotrophic taxa added | Type of analysis | Placement of taxa in analysis | Support for placement | Disruption of support in nearby branches<sup>a</sup> | Tree figure |
| --- | --- | --- | --- | --- | --- |
| (i) *Gymnosiphon* + *Apteria* (Burmanniaceae) | Partitioned ML (DNA) | – Sister to *Campylosiphon* (in Burmanniaceae) | Weak | Yes | App. S15 |
| (ii) *Thismia tentaculata* + *T. rodwayi* (Thismiaceae) | Partitioned ML (DNA) | – Sister to Taccaceae (Thismiaceae paraphyletic) | Weak | No | App. S16 |
| (iii) Afrothismiaceae | Partitioned ML (DNA) | – Sister to Poaceae | Weak | Yes | App. S6 |
| (iv) *Afrothismia gesnerioides* (Afrothismiaceae) | Partitioned ML (DNA) | – Nested within *Dioscorea* (Dioscoreaceae) | Weak | Yes | App. S11 |
| (iv) *Gymnosiphon*, *Apteria*, *Thismia tentaculata*, *T. rodwayi* and Afrothismiaceae | Partitioned ML (DNA) | – Afrothismiaceae sister to *Gymnosiphon*<br>– (*Gymnosiphon*, *Apteria*, Afrothismiaceae) sister to *B. itoana*<br>– *Thismia* sister to *Haplothismia* | All weak | Yes | App. S12 |
| (v) *Gymnosiphon*, *Apteria*, *Thismia tentaculata*, *T. rodwayi* and Afrothismiaceae | C60 mixture model ML (amino-acid) | – Afrothismiaceae sister to *Gymnosiphon*<br>– (*Gymnosiphon*, *Apteria*, Afrothismiaceae) sister to *B. itoana*<br>– *Thismia* sister to Taccaceae | All weak | Yes | App. S13 |
| (vi) *Gymnosiphon*, *Apteria*, *Thismia tentaculata*, *T. rodwayi* and Afrothismiaceae | CAT-GTR mixture model Bayesian (amino-acid) | – Afrothismiaceae sister to *Gymnosiphon*<br>– (*Gymnosiphon*, *Apteria*, Afrothismiaceae) sister to *Burmannia*<br>– *Thismia* sister to *Haplothismia* | All weak | No | App. S14 |

Notes: <sup>a</sup>At least 20% reduction in branch support (posterior probabilities as percentages) cf. DNA-based analysis of core 89-taxon set (Fig. 3, Appendix S3).

When Afrothismiaceae are included, this lineage appears to be attracted to distantly related sets of taxa, and broadly reduces bootstrap support across the tree (Table 2, cf. Fig. 3, Appendices S3, S4 and Appendices S6, S11). First, in a taxon-addition experiment in which only the four sampled members of Afrothismiaceae are added to the core set, these taxa are inferred to be the sister group of Poaceae in the order Poales, a clade with one of the longest branches among photosynthetic lineages, but with very poor bootstrap support for this arrangement (14%; Table 2, Appendix S6). There are also substantial drops in bootstrap support along the entire “backbone” of monocot phylogeny, including within Burmanniaceae (cf. Fig. 3, Appendices S3, S4 and Appendix S6). Adding only *Afrothismia gesneriodes* to the core set appears to be less disruptive to support values across the monocot phylogeny, including within Burmanniaceae (cf. Fig. 3, Appendices S3, S4 and Appendix S11). In this taxon-addition experiment, Afrothismiaceae instead place within *Dioscorea*, again with poor support (35%; Table 2, Appendix S11).

### C60 and CAT-GTR approaches compared to partitioned ML analyses

Likelihood and Bayesian approaches designed to ameliorate composition-related effects for challenging long-branch taxa did not yield substantially different placements of problematic long-branch taxa, compared to a standard partitioned ML analysis that included all 98 taxa sampled here (Appendices S12–S14). In all of these analyses, members of Thismiaceae form either a clade (partitioned ML analysis and PhyloBayes analysis) or a grade that is closely related to Taccaceae (Fig. 5), with moderate to poor bootstrap support for these arrangements. Afrothismiaceae are moderately to weakly supported as nested in Burmanniaceae (sister to the longest branch taxon there, *Gymnosiphon*), in all three analyses.

### Constraint tests of alternative placements of Thismiaceae and Afrothismiaceae

We can strongly reject alternative hypothesis (i). This hypothesis includes a constrained clade comprising Burmanniaceae and Thismiaceae (Appendix S5), compared to the unconstrained (best) likelihood tree (Fig. 3), in which these two lineages are recovered as distant relatives in Dioscoreales (*P* = 0.001 for both AU and SH tests; Table 3). None of the other constrained hypotheses, (ii)–(v), which place Afrothismiaceae in various positions could be rejected, compared to its unconstrained (presumably artefactual) placement as the sister group of Poaceae (Appendix S6), using either AU or SH tests (Table 3, Appendices S7–S10). These included an alternative position of Afrothismiaceae (ii) within or as the sister group of Dioscoreaceae, as suggested in Appendix S7 (*P* = 0.483 and *P* = 0.477 for AU and SH tests, respectively); (iii) within or as the sister group of Burmanniaceae, as suggested in Appendix S8 (*P* = 0.446 and *P* = 0.442); (iv) within (or as the sister group of) Thismiaceae, represented by *Haplothismia* (see Appendix S9), as suggested in previous studies (e.g., Maas-van de Kamer, 1998; *P* = 0.126 and *P* = 0.145); and (v) as the sister-group of a clade comprising Taccaceae and Thismiaceae (see Appendix S10), as suggested in previous studies (Merckx et al., 2009a and Lin et al., 2022; *P* = 0.15 and *P* = 0.155); see Table 3. Thus, these tests indicate that plastid phylogenomic data lack power to place Afrothismiaceae in the monocot tree of life.

**Table 3.** Likelihood-based tests showing that an alternative arrangement of Thismiaceae with Burmanniaceae can be ruled out (hypothesis i), while multiple alternative placements of Afrothismiaceae cannot (hypotheses ii–v); see main text (and see Appendices noted below for the topological constraints used). Results (*P* values) from approximately unbiased (AU) and Shimodaira-Hasegawa (SH) tests compare constrained vs. unconstrained optimal trees for the five alternative hypotheses. Asterisks (*) indicate significant *P* values (“NS” = non-significant).

| Alternative hypothesis | Constraint used | Approximately unbiased (AU) *P*-value | Shimodaira-Hasegawa (SH) *P*-value |
| --- | --- | --- | --- |
| i (Appendix S5) | Burmanniaceae and Thismiaceae (ignoring *Afrothismia*) | 0.001* | 0.001* |
| ii (Appendix S7) | Dioscoreaceae and *Afrothismia* | 0.483 (NS) | 0.477 (NS) |
| iii (Appendix S8) | Burmanniaceae and *Afrothismia* | 0.446 (NS) | 0.442 (NS) |
| iv (Appendix S9) | Thismiaceae s.l. (including *Afrothismia*) | 0.126 (NS) | 0.145 (NS) |
| v (Appendix S10) | Thismiaceae and Taccaceae plus *Afrothismia* | 0.15 (NS) | 0.155 (NS) |

## DISCUSSION

Our study provides the first large-scale survey of relationships in Dioscoreales, based on full plastid genomes for photosynthetic taxa and reduced plastid genomes of fully mycoheterotrophic taxa. This approach has broadly worked well elsewhere for plastid-based studies including fully mycoheterotrophic lineages (e.g., Lam et al., 2018; and see Fig. 3 here, for example), although here we found that adding rapidly evolving lineages can lead to local depression in branch support and thus minor uncertainties in taxon placement—as shown in the taxon-addition experiments for VLB taxa in Thismiaceae and Burmanniaceae (e.g., Appendices 15, 16) compared to the core taxon sampling that included more slowly evolving mycoheterotrophic lineages (Fig. 3; rates of evolution summarized in Fig. 2). Our overall understanding of higher-order relationships in Dioscoreales based on plastid data is therefore now highly consistent with a comparable study based on mitochondrial data (Fig. 4; summarized in Lin et al., 2022). The main exception concerns Afrothismiaceae (e.g., Fig. 4), which is recovered in multiple weakly supported positions in monocot phylogeny in the different taxon addition analyses adding different sets of taxa from the family (Appendices 6, 11, 12), or using different likelihood and Bayesian approaches (e.g., Appendices S12–S14). Our results therefore demonstrate that most but not all heterotrophic/VLB lineages can be accommodated in phylogenetic analyses of plastid genome data, with Afrothismiaceae as the main remaining problem.

### Dioscoreales relationships are now largely well understood

Our taxon sampling is complete at the genus level for photosynthetic Dioscoreales, and most higher-order relationships among these taxa are now resolved here (Fig. 1). A residual uncertainty concerns the earliest splits in Nartheciaceae, specifically the relative positions of *Aletris* and *Metanarthecium*—both to each other and to a clade comprising the three other genera in the family. These relationships were also uncertain in previous few-gene studies of Nartheciaceae (Merckx et al., 2008b; Fuse et al., 2012; Zhao et al., 2012), and may be associated with short deep branches that reflect rapid early diversification (e.g., Appendices S3, S4). An *Aletris*-*Metanarthecium* clade is strongly supported in our DNA analyses of the core taxon set (Fig. 3, Appendices S3, S4), which is completely consistent with moderately supported inferences from mitochondrial phylogenomic data (Fig. 4; Lin et al., 2022). However, this arrangement conflicts with the strongly supported placement of *Metanarthecium* as sister to the remaining Nartheciaceae genera in a recent DNA-based analysis of similar plastid phylogenomic data to that employed here (Garrett et al., 2023). The reason for this clash is unclear, although the latter analysis included numerous VLB taxa in Thismiaceae that we did not include here.

A second uncertainty for the photosynthetic taxa concerns the placement of *Stenomeris*, for which plastid phylogenomic data are currently lacking. The genus is either placed in Dioscoreaceae (e.g., APG, 2016), or has been recognized as its own family, Stenomeridaceae (reviewed in Caddick et al., 2002a). Here *Stenomeris* is recovered as the sister group of *Dioscorea* (Dioscoreaceae) with poor to moderate support based on the inclusion of only three available plastid genes in our phylogenomic analyses (Appendices S12–S14, S17). The placement of *Stenomeris* was also uncertain in previous phylogenetic studies that used a few plastid markers or morphology (Caddick et al., 2002a; Wilkin et al., 2005; Hsu et al., 2013; Viruel et al., 2016). A specimen of *Stenomeris dioscoreifolia* that was nested within *Dioscorea* with strong support in Merckx et al. (2009a, 2010) likely was mislabeled or contaminated (V. Merckx, unpublished data). A recent study based on nuclear phylogenomic data recovered *Stenomeris* as the sister-group to a clade comprising *Dioscorea*, *Tacca* and *Trichopus* with strong support (Zuntini et al., 2024). However, 50% of the ∼353 genes used in the coalescence-based analysis conflict with this arrangement, and so its position may still be considered to be ambiguous.

### Using plastid data to place mycoheterotrophs with high to very high substitution rates

Analyses of a core 89-taxon set here, which includes the least rapidly evolving full mycoheterotrophs sampled in Dioscoreales (Fig. 2), provide well supported inferences across most branches in the order (Fig. 3, Appendices S3, S4). The placements of the three FM taxa included in this set (i.e., *Burmannia itoana* and *Campylosiphon congestus* in Burmanniaceae; *Haplothismia exannulata* in Thismiaceae) and the remaining higher-order relationships in Dioscoreales are not sensitive to the use of DNA or AA data, although the latter yielded lower bootstrap support for some branches (Fig. 3, Appendices S3, S4). We obtained these results despite extensive gene loss in the three FM taxa (Fig. 1), suggesting that their moderately high substitution rates and patchy gene recovery do not strongly interfere with phylogenetic inference. This finding is consistent with (i) a previous study showing that phylogenetic analyses can benefit from inclusion of taxa with highly incomplete gene sets, as even small amounts of data can help to subdivide long branches (e.g., Wiens and Tiu, 2012); and (ii) phylogenomic studies that successfully place other mycoheterotrophic lineages, including Corsiaceae (Bodin et al., 2016; Givnish et al., 2016; Lam et al., 2018), *Geosiris* (Iridaceae; Joyce et al., 2018; Lam et al., 2018), *Petrosavia* (Petrosaviaceae; Logacheva et al., 2014; Lam et al., 2018), Triuridaceae (Lam et al., 2015, 2018; Soto Gomez et al., 2020), multiple Orchidaceae taxa (e.g., *Aphyllorchis, Cephalanthera, Corallorhiza, Cymbidium, Cyrtosia, Danxiaorchis, Degranvillea, Dipodium, Hetaeria, Lecanorchis, Neottia*, *Risleya*; Delannoy et al., 2011; Givnish et al., 2015; Feng et al., 2016; Lam et al., 2018; Kim et al., 2020; Li et al., 2020; Barrett et al., 2024a), and the eudicot families Ericaceae, Gentianaceae and Polygalaceae (Braukmann and Stefanović, 2012; Braukmann et al., 2017; Lam et al., 2018; Petersen et al., 2019).

The monophyly of the mycoheterotrophic family Burmanniaceae remains strongly supported in this analysis (setting aside the problematic behavior of Afrothismiaceae, e.g., Table 2). However, we were unable to robustly place where the *Apteria*-*Gymnosiphon* clade fits within Burmanniaceae, as the simultaneous addition of these two genera results in substantial reductions (≥20%) in bootstrap support for most branches in the family compared to the core taxon set (Table 2, cf. Fig. 3 and Appendix S15). This support reduction may represent a mild long-branch artefact. Analyses using mitochondrial phylogenomic data (Lin et al., 2022) also inferred a strongly supported *Apteria*-*Gymnosiphon* clade, in turn moderately supported as the sister-group of *Burmannia* plus *Campylosiphon* (Fig. 4). The latter arrangement is at odds with an older parsimony analysis with three nuclear and mitochondrial loci, which obtained a strongly supported *Apteria*-*Burmannia* clade, in turn sister to a strongly supported *Gymnosiphon*-*Hexapterella* clade (Merckx et al., 2008a). Although *Hexapterella* was not included here, or in Lin et al. (2022), the contrasting results in Merckx et al. (2008a) may represent another long-branch artefact in our studies. Broader sampling of Burmanniaceae in future studies (both mitochondrial- and plastid-based, for the more slowly evolving taxa) are needed to continue to clarify genus-level relationships, and the exact number of origins of full mycoheterotrophy in this family, currently estimated to be around eight such transitions (Merckx et al., 2013).

We expected to observe strong long-branch artefacts in the taxon-addition experiments for Thismiaceae, as the two added VLB species, *Thismia rodwayi* and *T. tentaculata*, have higher substitution rates than those in Burmanniaceae (see Fig. 2 for *T. tentaculata* and Appendix S16 for both species). However, our results from this experiment are broadly consistent with those from analyses of the core taxon set that included the moderately rate-elevated *Haplothismia* as the only Thismiaceae representative. Although we recovered Thismiaceae as a grade in two analyses (Appendices S13, S16), members of this FM family still form a larger clade with Taccaceae that is in turn inferred to be sister to Trichopodaceae, with only a slight reduction (<20%) in bootstrap support (Table 2, cf. Fig. 3 and Appendix S16). Thus, the arrangement recovered here for these three families is also largely consistent with recent plastid- and mitochondrial-based phylogenomic inferences (Lin et al., 2022; Garrett et al., 2023).

### Addressing the challenge of placing VLB mycoheterotrophs using plastid data

In general, model-based methods like maximum likelihood may be expected to be less prone to long-branch artefacts than distance methods like parsimony (Felsenstein, 1981; Huelsenbeck, 1995; Swofford et al., 2001; for examples see Nickrent et al., 2004; Merckx et al., 2009a; Lam et al., 2015, 2016, 2018). However, model-based analysis is not immune to long-branch problems, and so mixture models implemented in likelihood or Bayesian frameworks may help ameliorate the effects of long branches (e.g., Lartillot et al., 2007; Quang et al., 2008; Wang et al., 2018).

However, we also show here that these alternative model-based approaches can still be misled with respect to VLB taxa. This appears to be the case for Afrothismiaceae, a family with very severe rate elevation in its retained plastid genes (Fig. 2, Appendices S6, S11). We recovered multiple highly disjunct and conflicting placements for Afrothismiaceae across the variant likelihood and Bayesian analyses performed here (Fig. 4, Appendices S6, S11–S14) (see also Caddick et al., 2002a; Merckx et al., 2006, 2009a, 2010; Merckx and Bidartondo, 2008; Yokoyama et al., 2008; Shepeleva et al., 2020; Lin et al., 2022). The optimal (and absurd) placement of Afrothismiaceae here as the sister group of Poaceae (grasses) in the taxon-addition experiment that included all sampled Afrothismiaceae taxa (Appendix S6) is especially peculiar, and is very likely a long-branch artefact, as grasses represent one of the faster evolving photosynthetic lineages of angiosperms (Bousquet et al., 1992; De La Torre et al., 2017, and see Appendices S3, S4 here). This placement of Afrothismiaceae also disagrees with previous studies based on a single nuclear locus and one to two relatively slowly evolving mitochondrial genes (Merckx and Bidartondo, 2008; Merckx et al., 2010; Merckx and Smets, 2014), and with analyses based on a full complement of the protein-coding mitochondrial genes (Lin et al., 2022, and see Fig. 4). The latter non-plastid-based studies instead support Afrothismiaceae as being deeply nested within Dioscoreales close to Taccaceae, Trichopodaceae and the rest of Thismiaceae, but consistently outside of these three clades, with varying levels of branch support. This mitochondrial phylogenomic result helped justify the recognition of Afrothismiaceae as a distinct family (Cheek et al., 2024)

The inclusion of Afrothismiaceae in plastid-based analyses here also has a strong depressive effect on support values for branches near its point of attachment in the monocot tree (Table 2, Appendices S6, S11–S14). This type of long-branch artefact occurred whether Afrothismiaceae was added individually in lineage-specific phylogenetic experiments using DNA-based likelihood analyses (Appendices S6, S11), or in experiments that include the full complement of photosynthetic and FM taxa sampled here using likelihood analyses of DNA and AA data, and using Bayesian analyses of AA data (Appendices S12–S14). Finally, none of the placements of Afrothismiaceae observed here for plastid data can be rejected by AU or SH tests (Table 3, Appendices S7–S10). The position of Afrothismiaceae in Dioscoreales can therefore be considered as unresolved by using plastid genome data alone.

This may point to a limit to plastid-based phylogenetic analyses that incorporate extremely long branch taxa; in addition, methods that should reduce composition-related effects for long-branch taxa (Lartillot et al., 2007; Quang et al., 2008; Wang et al., 2018) do not appear to help here (Appendices S12–S14). Barrett et al. (2024a) recently showed that analyses that take account of heterotachy (lineage-specific variation in evolutionary rates) for long-branch lineages can yield well-supported inferences, based on a study of a rapidly evolving FM orchid (*Degranvillea dermaptera*). In a study of *Wullschlaegelia*, another rapidly evolving FM orchid lineage (Barrett et al., 2024b), they also pointed to possible limits to this approach related to model parameterization, considering retained genes and taxon-sampling. While we did not explore the issue of heterotachy here, we agree with Barrett et al. (2024a) that additional research is needed to compare the efficacy of different approaches—including comparisons with partitioned analyses and approaches addressing composition-related effects—for handling exceptionally rapidly evolving lineages such as Afrothismiaceae.

### Options for updating the higher-order classification of Dioscoreales

The current circumscription of photosynthetic Nartheciaceae is unproblematic, and the removal of Thismiaceae from Burmanniaceae leaves the latter as a well-defined family of autotrophic and mycoheterotrophic lineages. Separating Thismiaceae from Burmanniaceae in angiosperm classification schemes is strongly supported by our plastid-based analysis considering: (i) AU and SH tests, which reject constraining members of Burmanniaceae and Thismiaceae together (Table 3, Appendix S5), and (ii) the strongly supported intervening branches between these two lineages in analyses of the core taxon set (Fig. 3). Moreover, the Taccaceae-Thismiaceae clade that we inferred was also recovered in previous analyses based on plastid datasets comprising fewer Dioscoreales taxa (Givnish et al. 2018; Lam et al., 2018) or genes (Lam et al., 2016) or focusing on specific lineages in the order (i.e., Thismiaceae; Garrett et al., 2023). It is also consistent with inferences based on mitochondrial and nuclear datasets (Yokoyama et al., 2008; Merckx et al., 2009a, 2010; Merckx and Smets, 2014; Shepeleva et al., 2020; Lin et al. 2022).

There are several options for future adjustment to the family-level classification of Dioscoreales to accommodate these results. First, based on consideration of mitochondrial data and morphology, Afrothismiaceae were recently formalized as a new family in the order (Cheek et al., 2024). Alternative solutions that recognize fewer families ought to also be considered. One option would be to lump all lineages outside both Nartheciaceae and Burmanniaceae (defined narrowly to exclude Thismiaceae) into a single large family: Dioscoreaceae would have priority, and this would result in a reorganized three-family classification. However, this approach is undesirable in our view, as it sweeps considerable morphological diversity (e.g., Caddick et al., 2002a) into a single taxon, making it morphologically highly heterogeneous and hard to diagnose. A second solution would be to recognize multiple smaller families, including Taccaceae, Trichopodaceae, Thismiaceae, and the recently described Afrothismiaceae. This is in practice what some recent workers have done (Merckx et al., 2010; Lin et al., 2022), and is the scheme we employed here. A third solution would be to recognize Dioscoreaceae as including only *Dioscorea* (and possibly *Stenomeris*), and to recognize an expanded Taccaceae that incorporates additional families. For example, Afrothismiaceae and Thismiaceae could be lumped in Taccaceae, and perhaps Trichopodaceae too. This solution would yield a four- or five-family classification scheme for the order (depending on whether Trichophodaceae is recognized as a separate family).

All three of these options are consistent with a more narrowly defined Burmanniaceae (i.e., excluding members of Thismiaceae and Afrothismiaceae), and also maintain recognition of Nartheciaceae and Dioscoreaceae. However, an adequate classification of Dioscoreales depends in part on resolving the placement of photosynthetic *Stenomeris* within the order, a crucial priority for future research on the order. Deciding among these competing classification schemes also depends on additional considerations, such as the need to (i) minimize redundancy (e.g., lumping monotypic families that are sister to larger families), (ii) maximize support for monophyly, and (iii) facilitate straightforward identification by botanists (e.g., Backlund and Bremer, 1998). Regarding the latter criteria, we are currently engaged in a study to update our understanding of morphological evolution in Dioscoreales as a whole.

### Some conclusions, future work, and advice for other studies

Our study provides a roadmap for researchers attempting to place very rapidly evolving lineages as part of phylogenomic analysis. Here we used plastid genomic data to addresses the placement of VLB taxa within the yam order Dioscoreales, which is relevant to studies of its classification and evolution. To do this, we first characterized the range of rates in mycoheterotrophic lineages in a Bayesian relative rates analysis (Fig. 2). We used this to select three of the less rapidly evolving FM lineages in initial analyses with photosynthetic Dioscoreales (i.e., *Burmannia itoana* and *Campylosiphon* in Burmanniaceae, and *Haplothismia* in Thismiaceae). Our phylogenetic inferences of this “core” taxon set, which comprises all photosynthetic taxa in Dioscoreales, including several species of *Burmannia*—along with the three moderately rapidly evolving FM lineages—are in general (i) well supported, and (ii) highly congruent with studies based on mitochondrial data (Figs. 3, 4). The more slowly evolving mycoheterotrophic taxa may provide “anchors” that help to place the more rapidly evolving VLB taxa in our subsequent taxon-addition experiments. While the inclusion of VLB taxa leads to moderate to substantial reductions in branch support, this phenomenon is only localized in most cases (e.g., Fig. 5; Table 2). In addition, all added VLB taxa (setting aside Afrothismiaceae) are recovered in positions consistent with earlier studies (e.g., Garrett at al., 2023). Our study unambiguously confirms that Thismiaceae and Burmanniaceae evolved in distant parts of Dioscoreales, and thus experienced distantly convergent losses of photosynthesis. This result is inconsistent with current classification (APG, 2016), which should be updated. We review several possible solutions for an update classification, which we recommend should be postponed pending clarification of a careful re-examination of the morphological support for different options for family reorganization.

We also document how inclusion of members of, Afrothismiaceae, a VLB taxon, substantially disrupts phylogenetic inference using plastid data (Fig. 5). Fortunately, the position of Afrothismiaceae in Dioscoreales phylogeny is now well understood, thanks to phylogenetic analysis of the much more slowly evolving mitochondrial genome (e.g., Lin et al., 2022). Afrothismiaceae may be a “challenge too far” for plastid-based phylogenomic analyses at this point. This is so even after using likelihood and Bayesian approaches designed to ameliorate composition-related effects for challenging long-branch taxa (e.g., C60 and CAT-GTR; Lartillot et al., 2007; Quang et al., 2008; Wang et al., 2018). We recover this recently described family (Cheek et al., 2024) in highly disjunct positions across the monocot tree of life, based on a range of analyses using essentially the same plastid data set, and found no ability to distinguish among these placements, using likelihood-based tests of alternative trees. The inclusion of members of Afrothismiaceae also leads to widespread and severe reduction in branch support in plastid-based analyses. Plastid data sets that include Afrothismiaceae thus represent a potentially useful test case for the development of analytical approaches to deal with phylogenetic inference artefacts associated with very long branch taxa (see also Barrett et al., 2024).

## Acknowledgements

The authors thank the Royal Botanic Gardens, Kew, and the United States National Herbarium for providing material of *Stenomeris*, Sangjin Jo and Ki-Joong Kim (Korea University) for making plastome data for two sampled species of *Burmannia* available on GenBank, previous reviewers of an earlier version of this study, and Peter Stevens for discussion of Dioscoreales classification. This research was enabled in part by WestGrid (www.westgrid.ca), Compute Canada (www.computecanada.ca) and Research Computing at the James Hutton Institute (“UK’s Crop Diversity Bioinformatics HPC”; BBSRC grants BB/S019669/1 and BB/X019683/1). This research was supported by an NSERC Postgraduate Fellowship, UBC Four-Year Fellowship and Kew Future Leader Fellowship to MSG, and an NSERC Discovery Grant to SWG.

## Author Contributions

M.S.G., Q.L., V.K.Y.L, J.V., V.S.F.T.M. performed DNA extractions and DNA sequencing. M.S.G., N.J.K., V.K.Y.L. assembled genome skim data and extracted gene sets. M.S.G. performed analyses. M.S.G and S.W.G. conceived the study and wrote the paper, with contributions from all authors.

## Data Availability Statement

All data matrices are available in figshare (https://figshare.com/s/8470257673242e6bf43f).

## Supporting Information

Additional supporting information may be found online in the Supporting Information section at the end of the article.

## APPENDICES

Appendix S1. Specimen source information; herbarium abbreviations follow Thiers (continuously updated).

Appendix S2. Optimal substitution models for final partitioning schemes inferred using the ModelFinder function in IQ-TREE (see text). DNA analyses were partitioned using a gene-by-codon (“G x C”) scheme for protein-coding genes and individual partitions for rDNA genes; amino-acid (AA) analyses were partitioned by gene. (A) G x C partitioning scheme for the core 89-taxon DNA matrix; (B) gene-based partitioning scheme for the amino-acid version of the same core matrix; (C) G x C partitioning scheme for a 90-taxon DNA matrix after adding three genes for photosynthetic *Stenomeris* to the core matrix; (D) G x C partitioning scheme for a 91-taxon DNA matrix after adding two very long branch taxa of fully mycoheterotrophic Burmanniaceae (*Apteria aphylla*, *Gymnosiphon longistylis*) to the core matrix; (E) G x C partitioning scheme for a second 91-taxon DNA matrix after adding two very long branch taxa of fully mycoheterotrophic Thismiaceae (*Thismia rodwayi, T. tentaculata*) to the core matrix; (F) G x C partitioning scheme for a second 90-taxon DNA matrix after adding one of four sampled species of very long branch and fully mycoheterotrophic Afrothismiaceae (*Afrothismia gesnerioides*) to the core matrix; (G) G x C partitioning scheme for a 93-taxon DNA matrix after adding all four sampled species of very long branch and fully mycoheterotrophic Afrothismiaceae to the core matrix; and (H) G x C partitioning scheme for the full 98-taxon DNA matrix including all Dioscoreales taxa with available molecular data here. Genes are shown before the underscore and the ‘pos’ term after the underscore indicates the codon position (not applicable for rrn genes).

Appendix S3. DNA-based maximum-likelihood phylogeny of monocots inferred for a core taxon sampling in Dioscoreales that excludes very long branch (VLB) mycoheterotrophs and a data-deficient photosynthetic taxon (*Stenomeris*) in Dioscoreales. Relationships were inferred using partitioned likelihood analysis of 82 plastid genes (20–28 genes in fully mycoheterotrophic taxa; highlighted in grey). Bootstrap support values are indicated beside branches if <100%. The scale bar indicates estimated substitutions per site. The main left-hand trees in Figs. 3 and 5 are a subset of this tree.

Appendix S4. Amino acid (AA)-based maximum-likelihood phylogeny of monocots inferred for a core taxon sampling in Dioscoreales that excludes very long branch (VLB) mycoheterotrophs and a data-deficient photosynthetic taxon (*Stenomeris*) in Dioscoreales (this is amino-acid counterpart to Appendix S3). Relationships were inferred using partitioned likelihood analysis of 78 plastid genes (16–24 in fully mycoheterotrophic taxa; highlighted in grey). Bootstrap support values are indicated beside branches if <100%. The scale bar indicates estimated substitutions per site. The main right-hand tree in Fig. 3 is a subset of this tree.

Appendix S5. Likelihood-based phylogenetic inference of an alternative hypothesis in Dioscoreales in which Burmanniaceae and Thismiaceae are constrained to form a clade (i.e., hypothesis i in the main text). The single fixed branch (thick red line) acts as a constraint for two taxon partitions: one partition comprises Burmanniaceae and Thismiaceae (ignoring Afrothismiaceae here), and the second all remaining taxa. The “constrained” phylogeny was inferred using this single constraint in partitioned DNA-based likelihood analysis of 82 plastid genes (20–28 genes in fully mycoheterotrophic taxa; highlighted in grey). This resulting constrained tree was then tested against an unconstrained one which recovered a Thismiaceae-Taccaceae clade (see Fig. 3, left-hand tree, Appendix S3, and Table 3).

Appendix S6. DNA-based maximum-likelihood phylogeny of monocots inferred by expanding a core taxon sampling in Dioscoreales (taxa in black font, see Fig. 3, Appendix S3) to include four long-branch mycoheterotrophs from Afrothismiaceae (taxa in blue font). Relationships were inferred using partitioned likelihood analysis of 82 plastid genes (10–28 genes in fully mycoheterotrophic taxa; highlighted in grey), including sequences with internal stop codons for three of the added taxa (see text). Bootstrap support values are indicated beside branches if <100%; because of compression of the phylogram backbone, an inset figure is used to show support values for major clades. Non-monophyletic families are indicated with dotted lines. The scale bar indicates estimated substitutions per site.

Appendix S7. Likelihood-based phylogenetic inference of an alternative hypothesis in Dioscoreales in which Dioscoreaceae and Afrothismiaceae are constrained to form a clade (i.e., hypothesis ii in the main text). The fixed branch (thick red line) acts as a constraint for two taxon partitions; one partition comprises Dioscoreaceae and Afrothismiaceae (the latter represented by two species), and the second all remaining taxa. The “constrained” phylogeny was inferred using this single constraint in partitioned DNA-based likelihood analysis of 82 plastid genes (10–28 genes in fully mycoheterotrophic taxa; highlighted in grey). This resulting constrained tree was then tested against an unconstrained one which recovered an Afrothismiaceae-Poaceae clade (see Appendix S6 and Table 3).

Appendix S8. Likelihood-based phylogenetic of an alternative hypothesis in Dioscoreales in which Burmanniaceae and Afrothismiaceae are constrained to form a clade (i.e., hypothesis iii in the main text). The fixed branch (thick red line) acts as a constraint for two taxon partitions; one partition comprises Burmanniaceae and Afrothismiaceae (the latter represented by two species), and the second all remaining taxa. The “constrained” phylogeny was inferred using this single constraint in partitioned DNA-based likelihood analysis of 82 plastid genes (10–28 genes in fully mycoheterotrophic taxa; highlighted in grey). This resulting tree was then tested against an unconstrained one which recovered an Afrothismiaceae-Poaceae clade (see Appendix S6 and Table 3).

Appendix S9. Likelihood-based phylogenetic of an alternative hypothesis in Dioscoreales in which Thismiaceae and Afrothismiaceae are constrained to form a clade (i.e., hypothesis iv in the main text). The fixed branch (thick red line) acts as a constraint for two taxon partitions; one partition comprises Thismiaceae and Afrothismiaceae (the latter represented by two species), and the second all remaining taxa. The “constrained” phylogeny was inferred using this single constraint in partitioned DNA-based likelihood analysis of 82 plastid genes (10–28 genes in fully mycoheterotrophic taxa; highlighted in grey). This resulting tree was then tested against an unconstrained one which recovered an Afrothismiaceae-Poaceae clade (see Appendix S6 and Table 3).

Appendix S10. Likelihood-based phylogenetic inference of an alternative arrangement in Dioscoreales in which Thismiaceae and Taccaceae are constrained to be sister taxa, with this clade in turn constrained to be sister to Afrothismiaceae (i.e., hypothesis v in the main text). Afrothismiaceae are represented here by two taxa. Two fixed branches (thick red lines) constrain two nested sets of taxon partitions. In the first constraint, one taxon partition comprises Taccaceae and Thismiaceae, and the second the remaining taxa. In the second, one taxon partition comprises Afrothismiaceae, Taccaceae and Thismiaceae, and the second all remaining taxa. The “constrained” phylogeny was inferred using these nested constraints in partitioned DNA-based likelihood analysis of 82 plastid genes (10– 28 genes in fully mycoheterotrophic taxa; highlighted in grey). This resulting tree was then tested against an unconstrained one which recovered an Afrothismiaceae-Poaceae clade (see Appendix S6 and Table 3).

Appendix S11. DNA-based maximum-likelihood phylogeny of monocots inferred by expanding a core taxon sampling in Dioscoreales (taxa in black font) to include *Afrothismia gesnerioides* (species in blue font). The latter is a very long branch (VLB) mycoheterotroph in Afrothismiaceae that has retained plastid genes uninterrupted by internal stop codons (a contrasting condition with other members of Afrothismiaceae, see main text). Relationships were inferred using partitioned likelihood analysis of 82 plastid genes (10–28 genes in fully mycoheterotrophic taxa; highlighted in grey). Bootstrap support values are indicated beside branches if <100%; because of compression of the phylogram backbone, an inset figure is used to depict support values for major clades. Non-monophyletic families are indicated with dotted lines. The scale bar indicates estimated substitutions per site.

Appendix S12. DNA-based maximum-likelihood phylogeny of monocots inferred by expanding a core taxon sample (taxa in black font) to the full taxon sampling here, including all very long branch (VLB) mycoheterotrophs, and a data deficient photosynthetic taxon (*Stenomeris*) added simultaneously (taxa in blue font). Relationships were inferred using partitioned likelihood analysis of 82 plastid genes (7– 28 genes in fully mycoheterotrophic taxa; highlighted in grey). Bootstrap support values are indicated beside branches if <100%; because of compression of the phylogram backbone, an inset figure is used to show support values for major clades. The scale bar indicates estimated substitutions per site. The left-hand tree in Fig. 4 and the right-hand tree in Fig. 5 depict a subset of this tree.

Appendix S13. Amino acid (AA)-based monocot phylogeny inferred using a site-heterogeneous mixture model (C60) for the full taxon sampling (as in Appendix S11), which includes all very long branch (VLB) mycoheterotrophs, and a data deficient photosynthetic taxon (*Stenomeris*) added simultaneously (added taxa in blue font). Relationships were inferred using likelihood analysis of 78 plastid genes (16–24 in fully mycoheterotrophic taxa; highlighted in grey) under the site-heterogeneous mixture model. Bootstrap support values are indicated beside branches if <100%; because of compression of the phylogram backbone, an inset figure is used to show support values for major clades. The scale bar indicates estimated substitutions per site.

Appendix S14. Amino acid (AA)-based monocot phylogeny inferred using for a full taxon sampling after adding long-branch mycoheterotrophs and few-gene taxa in Dioscoreales (blue font) to the core set (black font). Relationships were inferred using Bayesian analysis of 78 plastid genes (16–24 in fully mycoheterotrophic taxa; highlighted in grey) under a site-heterogeneous mixture model (CAT-GTR). Bootstrap support values are indicated beside branches if <100%; due to compression of the backbone an inset of support values for major clades is included. The scale bar indicates estimated substitutions per site.

Appendix S15. DNA-based maximum-likelihood phylogeny of monocots inferred by expanding a core taxon sampling (taxa in black font) to include two long-branch mycoheterotrophs in Burmanniaceae (species in blue font). Relationships were inferred using partitioned likelihood analysis of 82 plastid genes (18–28 genes in fully mycoheterotrophic taxa; highlighted in grey). Bootstrap support values are indicated beside branches if <100%. The scale bar indicates estimated substitutions per site.

Appendix S16. DNA-based maximum-likelihood phylogeny of monocots inferred by expanding a core taxon sampling (taxa in black font) to include two long-branch mycoheterotrophs in Thismiaceae (species in blue font). Relationships were inferred using partitioned likelihood analysis of 82 plastid genes (7–28 genes in fully mycoheterotrophic taxa; highlighted in grey). Bootstrap support values are indicated beside branches if <100%; because of compression of the phylogram backbone, an inset figure is used to show support values for major clades. The scale bar indicates estimated substitutions per site.

Appendix S17. DNA-based maximum-likelihood phylogeny of monocots inferred by expanding a core taxon sampling (taxa in black font) to include a data-deficient photosynthetic taxon, *Stenomeris borneensis* (the species in blue font). Relationships were inferred using partitioned likelihood analysis of 82 plastid genes for most taxa, but included only three genes for *Stenomeris*, *atp*B, *mat*K, *rbc*L). Bootstrap support values are indicated beside branches if <100%. The scale bar indicates estimated substitutions per site.

