## Appendix S1 for "Placing very long branch taxa in the plant tree of life: a case study with Dioscoreales mycoheterotrophs"

Appendix S1. Specimen source information; herbarium abbreviations follow Thiers (continuously updated).

_________________________________________________________________________________________________________________________________

Species^a^	Family	Voucher number

[Collector number, herbarium]

_________________________________________________________________________________________________________________________________

**Dioscoreales**

*Metanarthecium luteoviride* Maxim.	Nartheciaceae	IV Taratenko s.n., MW

*Tacca leontopetaloides* (L.) Kuntze	Taccaceae	Wilkin 817, K

*Afrothismia gesnerioides* H.Maas	Afrothismiaceae	Sainge M. 2756, YA

*Afrothismia hydra* Sainge & T.Franke	Afrothismiaceae	Sainge 2624, YA

*Afrothismia* Schltr. sp.	Afrothismiaceae	Sainge M. 2625, YA

*Afrothismia winkleri* (Engl.) Schltr.	Afrothismiaceae	Sainge 2637, YA

*Trichopus sempervirens* (H.Perrier) Caddick & Wilkin	Trichopodaceae	Ranaivojaona, R. 790, K

_________________________________________________________________________________________________________________________________

^a^Additional sequences: *Aletris fauriei* (= *A. foliata* [Maxim.] Makino & Nemoto; NC_033412.1), *A. spicata* (Thunb.) Franch. (NC_033411.1), *Burmannia coelestis* D.Don (KT734618.1), *B. disticha* L. (NC_036661.1), *Dioscorea collettii* Hook.f. (NC_037717.1), *D. polystachya* Turcz. (NC_037716.1), *D. rotundata* (= *D. cayenensis* subsp. *rotundata* [Poir.] J.Miège; NC_024170.1), *D. villosa* L. (NC_034686.1), *D. zingiberensis* C.H.Wright (NC_027090.1), *Metanarthecium luteoviride* (NC_029214.1), *Stenomeris borneensis* (AF308018.1, AY973836.1, AF307475.1), *Tacca leontopetaloides* (NC_036658.1). See Garrett et al., (2023) for *Aletris obovata* Nash, *Dioscora membranacea* Pierre ex Prain & Burkill, *Haplothismia exannulata* Airy Shaw, *Narthecium californicum* Baker, *Nietneria paniculata* Steyerm., *Thismia rodwayi* F.Muell., and *Trichopus zeylanicus* Gaertn; see Lam et al., (2018) for the remaining taxa.
