## Appendix S2 for "Placing very long branch taxa in the plant tree of life: a case study with Dioscoreales mycoheterotrophs"

Appendix S2. Optimal substitution models for final partitioning schemes inferred using the ModelFinder function in IQ-TREE (see text). DNA analyses were partitioned using a gene-by-codon (“G x C”) scheme for protein-coding genes and individual partitions for rDNA genes; amino-acid (AA) analyses were partitioned by gene. (A) G x C partitioning scheme for the core 89-taxon DNA matrix; (B) gene-based partitioning scheme for the AA version of the same core matrix; (C) G x C partitioning scheme for a 90-taxon DNA matrix after adding three genes for photosynthetic *Stenomeris* to the core matrix; (D) G x C partitioning scheme for a 91-taxon DNA matrix after adding two very long branch taxa of fully mycoheterotrophic Burmanniaceae (*Apteria aphylla*, *Gymnosiphon longistylis*) to the core matrix; (E) G x C partitioning scheme for a second 91-taxon DNA matrix after adding two very long branch taxa of fully mycoheterotrophic Thismiaceae (*Thismia rodwayi, T. tentaculata*) to the core matrix; (F) G x C partitioning scheme for a second 90-taxon DNA matrix after adding one of four sampled species of very long branch and fully mycoheterotrophic *Afrothismia* (*A.* *gesnerioides*) to the core matrix; (G) G x C partitioning scheme for a 93-taxon DNA matrix after adding all four sampled species of very long branch and fully mycoheterotrophic *Afrothismia* to the core matrix; and (H) G x C partitioning scheme for the full 98-taxon DNA matrix including all Dioscoreales taxa with available molecular data here. Genes are shown before the underscore and the ‘pos’ term after the underscore indicates the codon position (not applicable for rrn genes).

_____________________________________________________________________________

No. of

partitions	Best model	Partition subsets

_____________________________________________________________________________

**(A)**

1	GTR+F+I+G	ACCD_pos1

2	TVM+F+G	ACCD_pos2, RPL22_pos2, RPS12_pos3

3	GTR+F+I+G	ACCD_pos3

4	GTR+F+I+G	ATPA_pos1, PSBJ_pos1

5	TVM+F+I+G	ATPA_pos2, ATPB_pos2

6	TVM+F+I+G	ATPA_pos3, PSBI_pos3

7	GTR+F+I+G	ATPB_pos1

8	TVM+F+G	ATPB_pos3, PETA_pos3

9	GTR+F+I+G	ATPE_pos1, RPOB_pos1

10	TVM+F+I+G	ATPE_pos2, NDHK_pos2, RPOC1_pos2

11	GTR+F+I+G	ATPE_pos3, RPL33_pos3, RPOB_pos3, RPOC1_pos3,

YCF4_pos3

12	TVM+F+I+G	ATPF_pos1, NDHK_pos1

13	HKY+F+I+G	ATPF_pos2, RPS16_pos2

14	TVM+F+I+G	ATPF_pos3, ATPH_pos3

15	GTR+F+I+G	ATPH_pos1, PSBC_pos1, PSBD_pos1

16	F81+F	ATPH_pos2

17	TVM+F+I+G	ATPI_pos1, PSBF_pos1, YCF3_pos1

18	TVM+F+I+G	ATPI_pos2, PETB_pos2, PSBI_pos2, PSBL_pos1

19	TVM+F+I+G	ATPI_pos3

20	TVM+F+I+G	CCSA_pos1, PSBF_pos3

21	GTR+F+I+G	CCSA_pos2, NDHF_pos2

22	GTR+F+I+G	CCSA_pos3, NDHF_pos3

23	GTR+F+I+G	CEMA_pos1, RPOA_pos1, YCF4_pos1

24	TVM+F+G	CEMA_pos2, PSBL_pos3, RPL23_pos3

25	TVM+F+G	CEMA_pos3

26	TVMef+G	CLPP_pos1

27	TVM+F+I+G	CLPP_pos2, RPS2_pos2

28	GTR+F+I+G	CLPP_pos3, RPS18_pos3, RPS2_pos3

29	TVM+F+G	INFA_pos1, INFA_pos2, RPS14_pos1

30	TVM+F+I+G	INFA_pos3, RPS19_pos3

31	GTR+F+I+G	MATK_pos1

32	TVM+F+I+G	MATK_pos2

33	TVM+F+I+G	MATK_pos3

34	GTR+F+I+G	NDHA_pos1, NDHD_pos1

35	TVM+F+I+G	NDHA_pos2, NDHD_pos2, NDHG_pos2

36	TVM+F+I+G	NDHA_pos3, NDHE_pos3

37	TIM3+F+I+G	NDHB_pos1, RPL23_pos2, RPS12_pos2

38	TIM2+F+I+G	NDHB_pos2, NDHC_pos2, PSBM_pos2, PSBN_pos2

39	TVM+F+G	NDHB_pos3

40	TVM+F+I+G	NDHC_pos1, NDHJ_pos1, PSBN_pos1, YCF4_pos2

41	GTR+F+I+G	NDHC_pos3, PSBK_pos3

42	GTR+F+I+G	NDHD_pos3

43	TVM+F+G	NDHE_pos1, NDHI_pos1, RPL33_pos1, RPL36_pos2

44	GTR+F+G	NDHE_pos2, PSBZ_pos2

45	TVM+F+I+G	NDHF_pos1

46	GTR+F+G	NDHG_pos1, PSBJ_pos3

47	GTR+F+I+G	NDHG_pos3

48	GTR+F+I+G	NDHH_pos1, RPOC1_pos1

49	TVM+F+I+G	NDHH_pos2

50	GTR+F+I+G	NDHH_pos3, PETD_pos3

51	TIM1+F+I+G	NDHI_pos2, NDHJ_pos2

52	GTR+F+I+G	NDHI_pos3

53	GTR+F+I+G	NDHJ_pos3, PSAA_pos3, PSBM_pos3

54	TVM+F+I+G	NDHK_pos3, PSBH_pos3

55	K81uf+F+I+G	PETA_pos1, RPS16_pos1

56	TVM+F+I+G	PETA_pos2, RPOB_pos2, YCF3_pos2

57	TIM1+F+I+G	PETB_pos1, PSBA_pos1

58	TVM+F+I+G	PETB_pos3, PSBB_pos3

59	TVMef+I+G	PETD_pos1

60	K81uf+F+I+G	PETD_pos2

61	TVMef+G	PETG_pos1, PETN_pos1, PSBE_pos1

62	TN93+F+I+G	PETG_pos2, PSBE_pos2, PSBL_pos2

63	TVM+F+I+G	PETG_pos3, RPL20_pos2, YCF3_pos3

64	TN93+F+I+G	PETL_pos1, RPS14_pos2

65	TVM+F+G	PETL_pos2, PSAJ_pos1, PSBK_pos2, PSBZ_pos1

66	GTR+F+G	PETL_pos3, PETN_pos3, PSAI_pos3, PSBE_pos3

67	HKY+F	PETN_pos2

68	K81uf+F+I+G	PSAA_pos1

69	GTR+F+I+G	PSAA_pos2, PSAB_pos2, PSBB_pos2

70	TVM+F+I+G	PSAB_pos1

71	GTR+F+I+G	PSAB_pos3

72	TVM+F+I+G	PSAC_pos1, PSBB_pos1

73	SYM+I	PSAC_pos2

74	GTR+F+I+G	PSAC_pos3, RBCL_pos3

75	TN93ef+G	PSAI_pos1

76	HKY+F+G	PSAI_pos2

77	SYM+G	PSAJ_pos2

78	K81uf+F+G	PSAJ_pos3

79	TIM3+F+I+G	PSBA_pos2, PSBF_pos2

80	GTR+F+I+G	PSBA_pos3

81	TVM+F+I+G	PSBC_pos2, PSBD_pos2

82	GTR+F+I+G	PSBC_pos3, PSBD_pos3

83	TIM3ef+I+G	PSBH_pos1, PSBH_pos2

84	TPM3+F+G	PSBI_pos1, PSBT_pos1, RPL23_pos1

85	HKY+F+I+G	PSBJ_pos2

86	TIM1ef+I+G	PSBK_pos1

87	TIM1+F+G	PSBM_pos1, RPL36_pos1

88	TN93+F+G	PSBN_pos3

89	TN93+F+I+G	PSBT_pos2

90	GTR+F+G	PSBT_pos3, RPS16_pos3

91	TIM1+F+I	PSBZ_pos3

92	GTR+F+I+G	RBCL_pos1

93	GTR+F+I+G	RBCL_pos2

94	GTR+F+I+G	RPL14_pos1

95	TN93+F+G	RPL14_pos2, RPS7_pos2

96	TVM+F+I+G	RPL14_pos3, RPS8_pos3

97	GTR+F+I+G	RPL16_pos1

98	SYM+I+G	RPL16_pos2

99	GTR+F+I+G	RPL16_pos3

100	GTR+F+I+G	RPL2_pos1, RPS12_pos1

101	SYM+I+G	RPL2_pos2, RRN23

102	K81uf+F+G	RPL2_pos3

103	K81uf+F+G	RPL20_pos1

104	TVM+F+I+G	RPL20_pos3, RPOC2_pos3

105	TVM+F+G	RPL22_pos1, RPL32_pos1, RPS15_pos1

106	TVM+F+I+G	RPL22_pos3, RPS11_pos3, RPS3_pos3

107	GTR+F+G	RPL32_pos2

108	GTR+F+I+G	RPL32_pos3

109	TVM+F+I+G	RPL33_pos2, RPOA_pos2, RPOC2_pos2

110	TIM3+F+G	RPL36_pos3

111	TVM+F+G	RPOA_pos3

112	GTR+F+I+G	RPOC2_pos1

113	TIM1+F+G	RPS11_pos1, RPS8_pos1

114	TIM3ef+G	RPS11_pos2

115	GTR+F+G	RPS14_pos3

116	TVM+F+I+G	RPS15_pos2

117	TVM+F+G	RPS15_pos3

118	GTR+F+G	RPS18_pos1

119	GTR+F+I+G	RPS18_pos2, RPS19_pos2, RPS4_pos1, RPS4_pos2

120	K81uf+F+G	RPS19_pos1, RPS7_pos1

121	GTR+F+G	RPS2_pos1

122	GTR+F+G	RPS3_pos1

123	GTR+F+I+G	RPS3_pos2

124	GTR+F+G	RPS4_pos3

125	GTR+F+G	RPS7_pos3

126	GTR+F+G	RPS8_pos2

127	TVM+F+G	YCF2_pos1

128	TVM+F+G	YCF2_pos2

129	TVM+F+G	YCF2_pos3

130	SYM+I+G	RRN16

131	K81+G	RRN4, RRN5

**(B)**

1	HIVw+F	ACCD

2	HIVw+F	ATPA

3	HIVb+F	ATPB, PSBT

4	HIVb	ATPE, RPOC1

5	HIVb	ATPF, RPL33

6	cpREV	ATPH

7	HIVw+F	ATPI, PSBF, PSBM, PSBN, YCF3

8	HIVw+F	CCSA, RPL20, RPS15

9	HIVw+F	CEMA, NDHG

10	FLU	CLPP

11	JTT	INFA, RPS11

12	HIVw+F	MATK

13	HIVw+F	NDHA, NDHI, NDHK

14	HIVw+F	NDHB, PETG, PSBI, PSBL

15	HIVw+F	NDHC, NDHJ

16	HIVb+F	NDHD, PSBK

17	mtREV+F	NDHE

18	HIVw+F	NDHF, RPS3

19	HIVb	NDHH

20	JTT-DCMut	PETA

21	FLU+F	PETB, PSBA, PSBB, PSBC, PSBD

22	HIVb	PETD

23	HIVw+F	PETL, RPOA, RPS19

24	mtZOA	PETN

25	FLU+F	PSAA, PSAB

26	JTT-DCMut	PSAC

27	HIVw	PSAI

28	cpREV	PSAJ

29	FLU	PSBE

30	FLU	PSBH

31	mtREV	PSBJ

32	mtZOA	PSBZ

33	LG	RBCL

34	HIVw	RPL14

35	cpREV	RPL16

36	HIVw+F	RPL2

37	FLU	RPL22

38	HIVw	RPL23

39	JTT	RPL32

40	HIVb+F	RPL36, RPOB

41	HIVw+F	RPOC2, RPS14

42	HIVb	RPS12

43	HIVb	RPS16

44	JTT+F	RPS18, RPS4, RPS8

45	HIVw	RPS2

46	cpREV	RPS7

47	HIVw+F	YCF2

48	HIVb+F	YCF4

**(C)**

1	GTR+F+I+G	ACCD_pos1

2	TVM+F+I+G	ACCD_pos2, RPL32_pos1

3	GTR+F+I+G	ACCD_pos3

4	GTR+F+I+G	ATPA_pos1

5	TVM+F+I+G	ATPA_pos2, ATPB_pos2

6	TVM+F+I+G	ATPA_pos3, PSBI_pos3

7	GTR+F+I+G	ATPB_pos1

8	TVM+F+G	ATPB_pos3, PETA_pos3

9	GTR+F+I+G	ATPE_pos1, RPOB_pos1

10	GTR+F+I+G	ATPE_pos2, NDHK_pos2

11	TVM+F+I+G	ATPE_pos3, YCF4_pos3

12	TVM+F+I+G	ATPF_pos1, NDHK_pos1

13	HKY+F+I+G	ATPF_pos2, RPS16_pos2

14	GTR+F+I+G	ATPF_pos3, RPOB_pos3

15	TIM1+F+I+G	ATPH_pos1

16	TPM3+F+I+G	ATPH_pos2, PSBA_pos2, PSBF_pos2

17	TVM+F+I+G	ATPH_pos3

18	TVM+F+I+G	ATPI_pos1, PSBN_pos1

19	TVM+F+I+G	ATPI_pos2, PETB_pos2

20	TVM+F+I+G	ATPI_pos3, NDHK_pos3

21	TVM+F+I+G	CCSA_pos1, PSBF_pos3

22	GTR+F+I+G	CCSA_pos2, NDHF_pos2

23	TVM+F+I+G	CCSA_pos3

24	TVM+F+G	CEMA_pos1, YCF4_pos1

25	TVM+F+G	CEMA_pos2, RPL23_pos3

26	TVM+F+I+G	CEMA_pos3, PSBH_pos3

27	TVMef+G	CLPP_pos1

28	TVM+F+I+G	CLPP_pos2, RPS2_pos2

29	TVM+F+I+G	CLPP_pos3, RPS18_pos3

30	TVM+F+G	INFA_pos1, RPS15_pos2

31	TVM+F+G	INFA_pos2, RPS14_pos1

32	TVM+F+I+G	INFA_pos3, RPS19_pos3

33	GTR+F+G	MATK_pos1

34	TVM+F+I+G	MATK_pos2

35	TVM+F+I+G	MATK_pos3

36	GTR+F+I+G	NDHA_pos1, NDHD_pos1

37	TPM3uf+F+I+G	NDHA_pos2, NDHG_pos2

38	TVM+F+I+G	NDHA_pos3, NDHE_pos3

39	TIM3+F+I+G	NDHB_pos1, YCF3_pos1

40	TN93+F+I+G	NDHB_pos2, NDHC_pos2

41	TVM+F+G	NDHB_pos3

42	TVM+F+G	NDHC_pos1, NDHJ_pos1

43	GTR+F+I+G	NDHC_pos3, PSBK_pos3

44	TVM+F+I+G	NDHD_pos2, PSAJ_pos1

45	GTR+F+I+G	NDHD_pos3

46	TVM+F+G	NDHE_pos1, NDHI_pos1

47	GTR+F+I+G	NDHE_pos2, PSBZ_pos2

48	TVM+F+I+G	NDHF_pos1

49	GTR+F+I+G	NDHF_pos3

50	GTR+F+G	NDHG_pos1, PSBJ_pos3

51	GTR+F+I+G	NDHG_pos3

52	GTR+F+I+G	NDHH_pos1, RPOC1_pos1

53	TVM+F+I+G	NDHH_pos2

54	TVM+F+I+G	NDHH_pos3

55	TIM1+F+I+G	NDHI_pos2, NDHJ_pos2

56	GTR+F+I+G	NDHI_pos3

57	TVM+F+I+G	NDHJ_pos3, PSBM_pos3

58	K81uf+F+I+G	PETA_pos1, RPS16_pos1

59	TVM+F+I+G	PETA_pos2, RPOB_pos2

60	TIM1+F+I+G	PETB_pos1, PSBA_pos1

61	TVM+F+I+G	PETB_pos3, PSBB_pos3

62	TVMef+I+G	PETD_pos1

63	GTR+F+I+G	PETD_pos2, PSAJ_pos2

64	GTR+F+G	PETD_pos3, PSBT_pos3

65	TVMef+I	PETG_pos1, PSBE_pos1

66	TN93+F+I+G	PETG_pos2, PSBE_pos2, PSBL_pos2

67	TVM+F+I+G	PETG_pos3, YCF3_pos3

68	TN93+F+I+G	PETL_pos1, RPS14_pos2

69	TPM3uf+F+G	PETL_pos2, PSBZ_pos1

70	TPM3uf+F+I+G	PETL_pos3, PETN_pos3

71	TIM2ef	PETN_pos1

72	TPM3uf+F	PETN_pos2, PSBI_pos2

73	K81uf+F+I+G	PSAA_pos1

74	GTR+F+I+G	PSAA_pos2, PSBB_pos2

75	GTR+F+I+G	PSAA_pos3

76	TVM+F+I+G	PSAB_pos1

77	TVM+F+I+G	PSAB_pos2, PSBC_pos2, PSBD_pos2

78	GTR+F+I+G	PSAB_pos3

79	TVM+F+I+G	PSAC_pos1, PSBB_pos1

80	SYM+I	PSAC_pos2

81	GTR+F+I+G	PSAC_pos3, RBCL_pos3

82	TN93ef+G	PSAI_pos1

83	HKY+F+G	PSAI_pos2

84	TIM3+F+G	PSAI_pos3, RPL22_pos2

85	K81uf+F+G	PSAJ_pos3

86	GTR+F+I+G	PSBA_pos3

87	GTR+F+I+G	PSBC_pos1, PSBD_pos1

88	GTR+F+I+G	PSBC_pos3, PSBD_pos3

89	TPM2uf+F+G	PSBE_pos3, PSBN_pos3

90	K81+G	PSBF_pos1, RPL23_pos2

91	TIM3ef+I+G	PSBH_pos1, PSBH_pos2

92	TIM3ef	PSBI_pos1, PSBT_pos1

93	GTR+F+G	PSBJ_pos1, RPL36_pos2

94	HKY+F+I+G	PSBJ_pos2

95	TIM1ef+I+G	PSBK_pos1

96	TIM3+F+I+G	PSBK_pos2

97	TIM2+F+I+G	PSBL_pos1, PSBN_pos2

98	TVM+F+I+G	PSBL_pos3, RPOA_pos2

99	TPM2uf+F+I+G	PSBM_pos1, YCF3_pos2

100	TN93+F+I	PSBM_pos2

101	TN93+F+I+G	PSBT_pos2

102	TIM1+F+I	PSBZ_pos3

103	GTR+F+I+G	RBCL_pos1

104	GTR+F+I+G	RBCL_pos2

105	GTR+F+I+G	RPL14_pos1

106	TN93+F+G	RPL14_pos2, RPS7_pos2

107	TVM+F+I+G	RPL14_pos3, RPS8_pos3

108	GTR+F+I+G	RPL16_pos1

109	SYM+I+G	RPL16_pos2

110	GTR+F+I+G	RPL16_pos3

111	GTR+F+I+G	RPL2_pos1

112	SYM+I+G	RPL2_pos2, RRN23

113	K81uf+F+G	RPL2_pos3

114	K81uf+F+G	RPL20_pos1

115	TIM1+F+G	RPL20_pos2, RPS12_pos3

116	TVM+F+I+G	RPL20_pos3

117	TVM+F+G	RPL22_pos1, RPS15_pos1

118	TVM+F+I+G	RPL22_pos3, RPS3_pos3

119	TPM3+F+G	RPL23_pos1

120	GTR+F+G	RPL32_pos2

121	GTR+F+I+G	RPL32_pos3

122	GTR+F+G	RPL33_pos1, RPOA_pos1

123	TVM+F+I+G	RPL33_pos2, RPOC2_pos2

124	GTR+F+G	RPL33_pos3, RPOA_pos3

125	TPM2uf+F+G	RPL36_pos1

126	TIM3+F+G	RPL36_pos3

127	TVM+F+I+G	RPOC1_pos2, YCF4_pos2

128	TVM+F+I+G	RPOC1_pos3

129	GTR+F+I+G	RPOC2_pos1

130	TVM+F+I+G	RPOC2_pos3

131	TIM1+F+G	RPS11_pos1, RPS8_pos1

132	TIM3ef+G	RPS11_pos2

133	TVM+F+G	RPS11_pos3

134	K81uf+F+G	RPS12_pos1

135	SYM+I+G	RPS12_pos2, RRN16

136	GTR+F+G	RPS14_pos3

137	TVM+F+G	RPS15_pos3

138	TVM+F+G	RPS16_pos3

139	GTR+F+G	RPS18_pos1

140	TIM1+F+G	RPS18_pos2, RPS4_pos1

141	GTR+F+G	RPS19_pos1, RPS2_pos1

142	TVM+F+I+G	RPS19_pos2, RPS4_pos2

143	GTR+F+G	RPS2_pos3

144	GTR+F+G	RPS3_pos1

145	GTR+F+I+G	RPS3_pos2

146	GTR+F+G	RPS4_pos3

147	K81uf+F+G	RPS7_pos1

148	GTR+F+G	RPS7_pos3

149	GTR+F+G	RPS8_pos2

150	TVM+F+G	YCF2_pos1

151	TVM+F+G	YCF2_pos2

152	TVM+F+G	YCF2_pos3

153	K81+G	RRN4, RRN5

**(D)**

1	GTR+F+I+G	ACCD_pos1

2	TVM+F+G	ACCD_pos2

3	GTR+F+I+G	ACCD_pos3

4	GTR+F+I+G	ATPA_pos1, PSBJ_pos1

5	TVM+F+I+G	ATPA_pos2, ATPB_pos2

6	TVM+F+I+G	ATPA_pos3, NDHH_pos3, PETD_pos3, PSBI_pos3

7	GTR+F+I+G	ATPB_pos1

8	TVM+F+G	ATPB_pos3, PETA_pos3

9	GTR+F+I+G	ATPE_pos1, RPOB_pos1

10	TVM+F+I+G	ATPE_pos2, NDHK_pos2, RPOC1_pos2

11	GTR+F+I+G	ATPE_pos3, ATPF_pos3, RPOB_pos3, RPOC1_pos3,

YCF4_pos3

12	TVM+F+I+G	ATPF_pos1, NDHK_pos1

13	HKY+F+I+G	ATPF_pos2, RPS16_pos2

14	GTR+F+I+G	ATPH_pos1, PSBC_pos1, PSBD_pos1

15	F81+F	ATPH_pos2

16	TVM+F+I+G	ATPH_pos3

17	TVM+F+I+G	ATPI_pos1, PSBF_pos1, YCF3_pos1

18	TVM+F+I+G	ATPI_pos2, PETB_pos2, PSBL_pos1

19	TVM+F+I+G	ATPI_pos3, PSAI_pos3

20	TVM+F+I+G	CCSA_pos1, PSBF_pos3

21	GTR+F+I+G	CCSA_pos2, NDHF_pos2

22	GTR+F+I+G	CCSA_pos3, NDHF_pos3

23	GTR+F+I+G	CEMA_pos1, RPOA_pos1, YCF4_pos1

24	TVM+F+G	CEMA_pos2, PSBL_pos3, RPL23_pos3

25	TVM+F+G	CEMA_pos3

26	TVMef+G	CLPP_pos1

27	TIM3+F+I+G	CLPP_pos2

28	GTR+F+I+G	CLPP_pos3, RPS14_pos3

29	TVM+F+G	INFA_pos1, INFA_pos2, RPS14_pos1

30	TVM+F+I+G	INFA_pos3, RPS19_pos3

31	GTR+F+I+G	MATK_pos1

32	TVM+F+I+G	MATK_pos2

33	TVM+F+I+G	MATK_pos3

34	GTR+F+I+G	NDHA_pos1, NDHD_pos1

35	TVM+F+I+G	NDHA_pos2, NDHD_pos2, NDHG_pos2

36	TVM+F+I+G	NDHA_pos3, NDHE_pos3

37	TIM3+F+I+G	NDHB_pos1, PSBT_pos1, RPL23_pos2

38	TIM2+F+I+G	NDHB_pos2, NDHC_pos2, PSBM_pos2, PSBN_pos2

39	TVM+F+G	NDHB_pos3

40	TVM+F+I+G	NDHC_pos1, NDHJ_pos1, PSBN_pos1, YCF4_pos2

41	GTR+F+I+G	NDHC_pos3, PSBK_pos3

42	GTR+F+I+G	NDHD_pos3

43	TVM+F+G	NDHE_pos1, NDHI_pos1, RPL33_pos1, RPL36_pos2

44	GTR+F+I+G	NDHE_pos2, PSBZ_pos2

45	TVM+F+I+G	NDHF_pos1

46	GTR+F+G	NDHG_pos1, PSBJ_pos3

47	GTR+F+I+G	NDHG_pos3, RPL33_pos3, RPOA_pos3

48	GTR+F+I+G	NDHH_pos1, RPOC1_pos1

49	TVM+F+I+G	NDHH_pos2

50	K81uf+F+I+G	NDHI_pos2, NDHJ_pos2, PSBM_pos1

51	GTR+F+I+G	NDHI_pos3

52	TVM+F+I+G	NDHJ_pos3, PETB_pos3, PSBM_pos3

53	TVM+F+I+G	NDHK_pos3, PSBH_pos3

54	K81uf+F+I+G	PETA_pos1, RPS16_pos1

55	TVM+F+I+G	PETA_pos2, RPOB_pos2, YCF3_pos2

56	TIM1+F+I+G	PETB_pos1, PSBA_pos1

57	SYM+I+G	PETD_pos1

58	K81uf+F+I+G	PETD_pos2

59	TVMef+G	PETG_pos1, PETN_pos1, PSBE_pos1, PSBI_pos1

60	TN93+F+I+G	PETG_pos2, PSBE_pos2, PSBL_pos2

61	TVM+F+I+G	PETG_pos3, RPL20_pos2, YCF3_pos3

62	TN93+F+I+G	PETL_pos1, RPS14_pos2

63	TVM+F+G	PETL_pos2, PSAJ_pos1, PSBK_pos2, PSBZ_pos1

64	TPM2uf+F+G	PETL_pos3, PSBE_pos3, PSBN_pos3

65	TPM3+F	PETN_pos2, PSBI_pos2

66	GTR+F+I+G	PETN_pos3, RPL22_pos2, RPS15_pos2, RPS3_pos2

67	K81uf+F+I+G	PSAA_pos1

68	GTR+F+I+G	PSAA_pos2, PSAB_pos2, PSBB_pos2

69	GTR+F+I+G	PSAA_pos3, PSBC_pos3, PSBD_pos3

70	TVM+F+I+G	PSAB_pos1

71	GTR+F+I+G	PSAB_pos3

72	TVM+F+I+G	PSAC_pos1, PSBB_pos1

73	SYM+I	PSAC_pos2

74	GTR+F+I+G	PSAC_pos3, RBCL_pos3

75	TN93ef+G	PSAI_pos1

76	HKY+F+G	PSAI_pos2

77	SYM+G	PSAJ_pos2

78	K81uf+F+G	PSAJ_pos3

79	TIM3+F+I+G	PSBA_pos2, PSBF_pos2

80	GTR+F+I+G	PSBA_pos3

81	TVM+F+I+G	PSBB_pos3

82	TVM+F+I+G	PSBC_pos2, PSBD_pos2

83	TIM3ef+I+G	PSBH_pos1, PSBH_pos2

84	HKY+F+I+G	PSBJ_pos2

85	TIM1ef+I+G	PSBK_pos1

86	TN93+F+I+G	PSBT_pos2

87	GTR+F+G	PSBT_pos3, RPS16_pos3

88	TIM1+F+I	PSBZ_pos3

89	GTR+F+I+G	RBCL_pos1

90	GTR+F+I+G	RBCL_pos2

91	GTR+F+I+G	RPL14_pos1

92	GTR+F+I+G	RPL14_pos2, RPL2_pos2, RPS12_pos2, RPS7_pos2

93	TVM+F+I+G	RPL14_pos3

94	GTR+F+G	RPL16_pos1

95	SYM+I+G	RPL16_pos2

96	TVM+F+I+G	RPL16_pos3, RPL22_pos3, RPS3_pos3, RPS8_pos3

97	GTR+F+G	RPL2_pos1

98	K81uf+F+G	RPL2_pos3

99	K81uf+F+G	RPL20_pos1

100	GTR+F+I+G	RPL20_pos3, RPOC2_pos3

101	TVM+F+G	RPL22_pos1, RPL32_pos1, RPS15_pos1

102	TPM3+F+G	RPL23_pos1

103	GTR+F+G	RPL32_pos2

104	GTR+F+I+G	RPL32_pos3

105	TVM+F+I+G	RPL33_pos2, RPOA_pos2, RPOC2_pos2

106	TPM2uf+F+G	RPL36_pos1

107	TIM3+F+G	RPL36_pos3

108	GTR+F+I+G	RPOC2_pos1

109	TVM+F+G	RPS11_pos1

110	TIM3ef+G	RPS11_pos2

111	TVM+F+G	RPS11_pos3

112	TVM+F+G	RPS12_pos1

113	K81uf+F+G	RPS12_pos3

114	TVM+F+G	RPS15_pos3

115	TVM+F+G	RPS18_pos1

116	TVM+F+G	RPS18_pos2, RPS4_pos1, RPS4_pos2

117	GTR+F+I+G	RPS18_pos3

118	K81uf+F+G	RPS19_pos1, RPS7_pos1

119	TPM3uf+F+G	RPS19_pos2, RPS2_pos2

120	GTR+F+G	RPS2_pos1

121	GTR+F+G	RPS2_pos3

122	GTR+F+G	RPS3_pos1, RPS8_pos1

123	GTR+F+G	RPS4_pos3

124	GTR+F+G	RPS7_pos3

125	GTR+F+G	RPS8_pos2

126	TVM+F+G	YCF2_pos1

127	TVM+F+G	YCF2_pos2

128	TVM+F+G	YCF2_pos3

129	SYM+I+G	RRN16, RRN4

130	SYM+I+G	RRN23

131	K80+G	RRN5

**(E)**

1	GTR+F+I+G	ACCD_pos1

2	TVM+F+I+G	ACCD_pos2

3	GTR+F+I+G	ACCD_pos3

4	GTR+F+I+G	ATPA_pos1, PSBJ_pos1

5	TVM+F+I+G	ATPA_pos2, ATPB_pos2

6	TVM+F+I+G	ATPA_pos3, NDHH_pos3, PETD_pos3, PSBI_pos3

7	GTR+F+I+G	ATPB_pos1

8	TVM+F+G	ATPB_pos3, PETA_pos3

9	GTR+F+I+G	ATPE_pos1, RPOB_pos1

10	TVM+F+I+G	ATPE_pos2, NDHK_pos2, RPOC1_pos2

11	GTR+F+I+G	ATPE_pos3, ATPF_pos3, RPOB_pos3, RPOC1_pos3,

YCF4_pos3

12	TVM+F+I+G	ATPF_pos1, NDHK_pos1

13	HKY+F+I+G	ATPF_pos2, RPS16_pos2

14	GTR+F+I+G	ATPH_pos1, PSBC_pos1, PSBD_pos1

15	F81+F	ATPH_pos2

16	TVM+F+I+G	ATPH_pos3

17	TVM+F+I+G	ATPI_pos1, PSBF_pos1, YCF3_pos1

18	GTR+F+I+G	ATPI_pos2, NDHE_pos2, PETB_pos2, PSBI_pos2,

PSBL_pos1

19	TVM+F+I+G	ATPI_pos3, NDHK_pos3, PSAI_pos3, PSBH_pos3

20	TVM+F+I+G	CCSA_pos1, PSBF_pos3

21	GTR+F+I+G	CCSA_pos2, NDHF_pos2

22	GTR+F+I+G	CCSA_pos3, NDHF_pos3

23	GTR+F+I+G	CEMA_pos1, RPOA_pos1, YCF4_pos1

24	TVM+F+G	CEMA_pos2, PSBL_pos3, RPL23_pos3

25	TVM+F+G	CEMA_pos3

26	TVMef+G	CLPP_pos1

27	TIM3+F+I+G	CLPP_pos2

28	GTR+F+I+G	CLPP_pos3, RPS14_pos3

29	TVM+F+G	INFA_pos1, INFA_pos2, RPS14_pos1

30	TVM+F+I+G	INFA_pos3, RPL22_pos3, RPL32_pos3

31	GTR+F+I+G	MATK_pos1

32	TVM+F+I+G	MATK_pos2

33	TVM+F+I+G	MATK_pos3

34	GTR+F+I+G	NDHA_pos1, NDHD_pos1

35	TVM+F+I+G	NDHA_pos2, NDHD_pos2, NDHG_pos2

36	TVM+F+I+G	NDHA_pos3, NDHE_pos3

37	TIM3+F+I+G	NDHB_pos1, PSBT_pos1, RPL23_pos2

38	TIM2+F+I+G	NDHB_pos2, NDHC_pos2, PSBM_pos2, PSBN_pos2

39	TVM+F+G	NDHB_pos3

40	TVM+F+I+G	NDHC_pos1, NDHJ_pos1, PSBN_pos1, YCF4_pos2

41	GTR+F+I+G	NDHC_pos3, PSBK_pos3

42	GTR+F+I+G	NDHD_pos3

43	TVM+F+G	NDHE_pos1, NDHI_pos1, RPL33_pos1, RPL36_pos2

44	TVM+F+I+G	NDHF_pos1

45	GTR+F+G	NDHG_pos1, PSBJ_pos3

46	GTR+F+I+G	NDHG_pos3, RPL33_pos3, RPOA_pos3

47	GTR+F+I+G	NDHH_pos1, RPOC1_pos1

48	TVM+F+I+G	NDHH_pos2

49	K81uf+F+I+G	NDHI_pos2, NDHJ_pos2, PSBM_pos1

50	GTR+F+I+G	NDHI_pos3

51	TVM+F+I+G	NDHJ_pos3, PETB_pos3, PSBM_pos3

52	K81uf+F+I+G	PETA_pos1, RPS16_pos1

53	TVM+F+I+G	PETA_pos2, RPOB_pos2, YCF3_pos2

54	TIM1+F+I+G	PETB_pos1, PSBA_pos1

55	TVMef+I+G	PETD_pos1

56	K81uf+F+I+G	PETD_pos2

57	TVMef+I	PETG_pos1, PSBE_pos1, PSBI_pos1

58	TN93+F+I+G	PETG_pos2, PSBE_pos2, PSBL_pos2

59	TVM+F+I+G	PETG_pos3, RPL20_pos2, YCF3_pos3

60	TN93+F+I+G	PETL_pos1, PSAI_pos1, RPS14_pos2

61	GTR+F+G	PETL_pos2, PSAJ_pos1, PSBZ_pos1

62	TPM2uf+F+G	PETL_pos3, PSBE_pos3, PSBN_pos3

63	K81+G	PETN_pos1, RRN4, RRN5

64	TIM3+F+G	PETN_pos2, PSBZ_pos2

65	TIM3+F+I+G	PETN_pos3, RPL22_pos2, RPS15_pos2

66	K81uf+F+I+G	PSAA_pos1

67	GTR+F+I+G	PSAA_pos2, PSAB_pos2, PSBB_pos2

68	GTR+F+I+G	PSAA_pos3, PSBC_pos3, PSBD_pos3

69	TVM+F+I+G	PSAB_pos1

70	GTR+F+I+G	PSAB_pos3

71	TVM+F+I+G	PSAC_pos1, PSBB_pos1

72	SYM+I	PSAC_pos2

73	GTR+F+I+G	PSAC_pos3, PSBB_pos3, RBCL_pos3

74	HKY+F+G	PSAI_pos2

75	GTR+F+I+G	PSAJ_pos2, PSBK_pos2

76	K81uf+F+G	PSAJ_pos3

77	TIM3+F+I+G	PSBA_pos2, PSBF_pos2

78	GTR+F+I+G	PSBA_pos3

79	TVM+F+I+G	PSBC_pos2, PSBD_pos2

80	TIM3ef+I+G	PSBH_pos1, PSBH_pos2

81	HKY+F+I+G	PSBJ_pos2

82	TIM1ef+I+G	PSBK_pos1

83	TN93+F+I+G	PSBT_pos2

84	GTR+F+G	PSBT_pos3, RPS16_pos3

85	TIM1+F+I	PSBZ_pos3

86	GTR+F+I+G	RBCL_pos1

87	GTR+F+I+G	RBCL_pos2

88	GTR+F+I+G	RPL14_pos1

89	TIM1+F+G	RPL14_pos2, RPS7_pos2

90	TVM+F+I+G	RPL14_pos3, RPL36_pos3

91	GTR+F+G	RPL16_pos1

92	SYM+G	RPL16_pos2

93	GTR+F+I+G	RPL16_pos3

94	GTR+F+G	RPL2_pos1, RPL2_pos2

95	K81uf+F+I+G	RPL2_pos3

96	TVM+F+G	RPL20_pos1, RPL22_pos1, RPL32_pos1, RPS15_pos1

97	GTR+F+I+G	RPL20_pos3, RPOC2_pos3

98	TPM3+F+G	RPL23_pos1

99	GTR+F+G	RPL32_pos2

100	TVM+F+I+G	RPL33_pos2, RPOA_pos2, RPOC2_pos2

101	K81uf+F+G	RPL36_pos1, RPS7_pos1

102	GTR+F+I+G	RPOC2_pos1

103	TIM1+F+G	RPS11_pos1, RPS8_pos1

104	TIM3ef+G	RPS11_pos2

105	TVM+F+G	RPS11_pos3

106	TVM+F+G	RPS12_pos1

107	SYM+G	RPS12_pos2, RRN16

108	K81uf+F+G	RPS12_pos3

109	TVM+F+G	RPS15_pos3

110	GTR+F+G	RPS18_pos1

111	TIM1+F+G	RPS18_pos2

112	GTR+F+I+G	RPS18_pos3

113	K81uf+F+G	RPS19_pos1

114	TIM3+F+G	RPS19_pos2

115	GTR+F+G	RPS19_pos3

116	GTR+F+G	RPS2_pos1

117	TVM+F+G	RPS2_pos2

118	GTR+F+G	RPS2_pos3

119	GTR+F+G	RPS3_pos1

120	GTR+F+I+G	RPS3_pos2

121	TVM+F+I+G	RPS3_pos3

122	TVM+F+G	RPS4_pos1, RPS4_pos2

123	GTR+F+G	RPS4_pos3

124	GTR+F+G	RPS7_pos3

125	GTR+F+G	RPS8_pos2

126	TVM+F+G	RPS8_pos3

127	TVM+F+G	YCF2_pos1

128	TVM+F+G	YCF2_pos2

129	TVM+F+G	YCF2_pos3

130	SYM+G	RRN23

**(F)**

1	GTR+F+I+G	ACCD_pos1

2	TVM+F+I+G	ACCD_pos2

3	GTR+F+I+G	ACCD_pos3

4	GTR+F+I+G	ATPA_pos1

5	TVM+F+I+G	ATPA_pos2, ATPB_pos2

6	TVM+F+I+G	ATPA_pos3, NDHH_pos3, PETD_pos3, PSBI_pos3

7	GTR+F+I+G	ATPB_pos1

8	TVM+F+G	ATPB_pos3, PETA_pos3

9	GTR+F+I+G	ATPE_pos1, RPL36_pos2, RPOB_pos1

10	GTR+F+I+G	ATPE_pos2, NDHK_pos2

11	TVM+F+I+G	ATPE_pos3, RPOC1_pos3, YCF4_pos3

12	TVM+F+I+G	ATPF_pos1, NDHK_pos1, PSBJ_pos1

13	HKY+F+I+G	ATPF_pos2, RPS16_pos2

14	GTR+F+I+G	ATPF_pos3, CLPP_pos3, RPOB_pos3, RPS14_pos3

15	GTR+F+I+G	ATPH_pos1, PSBC_pos1, PSBD_pos1

16	F81+F	ATPH_pos2

17	TVM+F+I+G	ATPH_pos3

18	TVM+F+I+G	ATPI_pos1, PSBF_pos1, YCF3_pos1

19	GTR+F+I+G	ATPI_pos2, NDHE_pos2, PETB_pos2, PSBI_pos2,

PSBL_pos1

20	TVM+F+I+G	ATPI_pos3, CEMA_pos3

21	TVM+F+I+G	CCSA_pos1

22	GTR+F+I+G	CCSA_pos2, NDHF_pos2

23	GTR+F+I+G	CCSA_pos3, NDHF_pos3

24	GTR+F+I+G	CEMA_pos1, RPOA_pos1, YCF4_pos1

25	TVM+F+G	CEMA_pos2, PSBL_pos3

26	GTR+F+G	CLPP_pos1

27	TVM+F+I+G	CLPP_pos2, RPOC1_pos2, YCF4_pos2

28	TVM+F+G	INFA_pos1, RPL33_pos1, RPS8_pos1

29	GTR+F+G	INFA_pos2, NDHE_pos1, NDHI_pos1, RPS14_pos1,

RPS2_pos1

30	TVM+F+I+G	INFA_pos3, RPS15_pos3

31	GTR+F+I+G	MATK_pos1

32	TVM+F+I+G	MATK_pos2, PETL_pos3, PSAI_pos3

33	TVM+F+I+G	MATK_pos3

34	GTR+F+I+G	NDHA_pos1, NDHD_pos1

35	TVM+F+I+G	NDHA_pos2, NDHD_pos2, NDHG_pos2

36	TVM+F+I+G	NDHA_pos3, NDHE_pos3

37	TIM3+F+I+G	NDHB_pos1, PSBT_pos1

38	TIM2+F+I+G	NDHB_pos2, NDHC_pos2, PSBM_pos2, PSBN_pos2

39	TVM+F+G	NDHB_pos3, RPL36_pos1

40	TVM+F+I+G	NDHC_pos1, NDHJ_pos1, PSBN_pos1

41	GTR+F+I+G	NDHC_pos3, PSBK_pos3

42	GTR+F+I+G	NDHD_pos3

43	TVM+F+I+G	NDHF_pos1

44	GTR+F+G	NDHG_pos1, PSBJ_pos3

45	GTR+F+I+G	NDHG_pos3, RPL33_pos3, RPOA_pos3

46	GTR+F+I+G	NDHH_pos1, RPOC1_pos1

47	TVM+F+I+G	NDHH_pos2

48	K81uf+F+I+G	NDHI_pos2, NDHJ_pos2, PSBM_pos1

49	GTR+F+I+G	NDHI_pos3

50	GTR+F+I+G	NDHJ_pos3, PSAA_pos3, PSBM_pos3

51	TVM+F+I+G	NDHK_pos3, PSBH_pos3

52	K81uf+F+I+G	PETA_pos1, RPS16_pos1

53	TVM+F+I+G	PETA_pos2, RPOB_pos2, YCF3_pos2

54	TIM1+F+I+G	PETB_pos1, PSBA_pos1

55	TVM+F+I+G	PETB_pos3, PSBB_pos3

56	TVMef+I+G	PETD_pos1

57	K81uf+F+I+G	PETD_pos2

58	TVMef+G	PETG_pos1, PETN_pos1, PSBE_pos1, PSBI_pos1,

RRN5

59	TN93+F+I+G	PETG_pos2, PSBE_pos2, PSBL_pos2

60	TVM+F+I+G	PETG_pos3, RPL20_pos2, YCF3_pos3

61	TIM2ef+G	PETL_pos1

62	TVM+F+G	PETL_pos2, PSAJ_pos1, PSBK_pos2, PSBZ_pos1

63	HKY+F	PETN_pos2

64	TPM2uf+F+G	PETN_pos3, PSBE_pos3, PSBN_pos3

65	K81uf+F+I+G	PSAA_pos1

66	GTR+F+I+G	PSAA_pos2, PSAB_pos2, PSBB_pos2

67	TVM+F+I+G	PSAB_pos1

68	GTR+F+I+G	PSAB_pos3

69	TVM+F+I+G	PSAC_pos1, PSBB_pos1

70	SYM+I	PSAC_pos2

71	GTR+F+I+G	PSAC_pos3, RPL14_pos3, RPL36_pos3

72	TN93ef+G	PSAI_pos1

73	HKY+F+G	PSAI_pos2

74	SYM+G	PSAJ_pos2

75	K81uf+F+G	PSAJ_pos3

76	TIM3+F+I+G	PSBA_pos2, PSBF_pos2

77	GTR+F+I+G	PSBA_pos3

78	TVM+F+I+G	PSBC_pos2, PSBD_pos2

79	GTR+F+I+G	PSBC_pos3, PSBD_pos3

80	TIM3+F+G	PSBF_pos3, PSBZ_pos3, RPL22_pos2

81	TIM3ef+I+G	PSBH_pos1, PSBH_pos2

82	HKY+F+I+G	PSBJ_pos2

83	TIM1ef+I+G	PSBK_pos1

84	TN93+F+I+G	PSBT_pos2

85	TVM+F+G	PSBT_pos3, RPS8_pos3

86	TIM3+F+G	PSBZ_pos2

87	GTR+F+I+G	RBCL_pos1

88	GTR+F+I+G	RBCL_pos2

89	GTR+F+I+G	RBCL_pos3

90	GTR+F+G	RPL14_pos1

91	TIM1+F+G	RPL14_pos2, RPS7_pos2

92	GTR+F+I+G	RPL16_pos1

93	SYM+I+G	RPL16_pos2

94	GTR+F+I+G	RPL16_pos3, RPL32_pos3

95	TIM2+F+G	RPL2_pos1, RPL2_pos2

96	K81uf+F+G	RPL2_pos3

97	GTR+F+G	RPL20_pos1, RPL32_pos2

98	GTR+F+I+G	RPL20_pos3, RPOC2_pos3

99	TVM+F+G	RPL22_pos1, RPL32_pos1, RPS15_pos1

100	TVM+F+I+G	RPL22_pos3

101	TPM3+F+G	RPL23_pos1

102	TPM2uf+F+G	RPL23_pos2, RRN4.5

103	GTR+F+G	RPL23_pos3, RPS2_pos2, RPS7_pos3

104	TVM+F+I+G	RPL33_pos2, RPOA_pos2, RPOC2_pos2

105	GTR+F+I+G	RPOC2_pos1

106	SYM+G	RPS11_pos1

107	TIM3ef+G	RPS11_pos2

108	TVM+F+I+G	RPS11_pos3, RPS3_pos3

109	TIM1+F+G	RPS12_pos1

110	TIM1ef+G	RPS12_pos2, RRN16

111	GTR+F+G	RPS12_pos3

112	TN93+F+G	RPS14_pos2

113	GTR+F+I+G	RPS15_pos2, RPS8_pos2

114	TVM+F+G	RPS16_pos3

115	GTR+F+G	RPS18_pos1

116	TVM+F+G	RPS18_pos2, RPS4_pos1, RPS4_pos2

117	GTR+F+G	RPS18_pos3

118	K81uf+F+G	RPS19_pos1

119	TPM3+F+G	RPS19_pos2

120	GTR+F+G	RPS19_pos3

121	GTR+F+I+G	RPS2_pos3

122	GTR+F+G	RPS3_pos1

123	GTR+F+I+G	RPS3_pos2

124	GTR+F+G	RPS4_pos3

125	K81uf+F+G	RPS7_pos1

126	TVM+F+G	YCF2_pos1

127	TVM+F+G	YCF2_pos2

128	TVM+F+G	YCF2_pos3

129	GTR+F+G	RRN23

**(G)**

1	TVM+F+I+G	ACCD_pos1, RPL22_pos1

2	TVM+F+I+G	ACCD_pos2, RPL20_pos1, RPL32_pos1, RPS15_pos1

3	GTR+F+I+G	ACCD_pos3

4	GTR+F+I+G	ATPA_pos1, NDHH_pos1, PSBJ_pos1

5	TVM+F+I+G	ATPA_pos2, ATPB_pos2

6	TVM+F+I+G	ATPA_pos3, PSBI_pos3

7	GTR+F+I+G	ATPB_pos1, ATPI_pos1, PSBN_pos1

8	TVM+F+I+G	ATPB_pos3, CLPP_pos3, PETA_pos3, RPS2_pos3

9	GTR+F+I+G	ATPE_pos1, RPL36_pos2, RPOB_pos1

10	TVM+F+I+G	ATPE_pos2, NDHK_pos2, RPOC1_pos2

11	GTR+F+I+G	ATPE_pos3, RPL33_pos3, RPOB_pos3, RPOC1_pos3,

YCF4_pos3

12	TVM+F+I+G	ATPF_pos1, NDHK_pos1

13	HKY+F+I+G	ATPF_pos2, RPS16_pos2

14	GTR+F+G	ATPF_pos3, RPS14_pos3, RPS18_pos3, RPS4_pos3

15	GTR+F+I+G	ATPH_pos1, PSBC_pos1, PSBD_pos1

16	F81+F	ATPH_pos2

17	TVM+F+I+G	ATPH_pos3

18	TVM+F+I+G	ATPI_pos2, PETB_pos2, PSBL_pos1

19	TVM+F+I+G	ATPI_pos3

20	TVM+F+I+G	CCSA_pos1

21	GTR+F+I+G	CCSA_pos2, NDHF_pos2

22	GTR+F+I+G	CCSA_pos3, NDHF_pos3

23	GTR+F+I+G	CEMA_pos1, RPOA_pos1, YCF4_pos1

24	TVM+F+G	CEMA_pos2

25	TVM+F+G	CEMA_pos3

26	GTR+F+G	CLPP_pos1

27	TPM3uf+F+I+G	CLPP_pos2

28	TVM+F+G	INFA_pos1, INFA_pos2, RPL2_pos3, RPS14_pos1

29	TVM+F+I+G	INFA_pos3, RPS15_pos3

30	GTR+F+I+G	MATK_pos1

31	TVM+F+I+G	MATK_pos2

32	GTR+F+I+G	MATK_pos3

33	GTR+F+I+G	NDHA_pos1, NDHD_pos1

34	TVM+F+I+G	NDHA_pos2, NDHD_pos2, NDHG_pos2

35	TVM+F+I+G	NDHA_pos3, NDHE_pos3

36	TIM3+F+I+G	NDHB_pos1, PSBT_pos1

37	TIM2+F+I+G	NDHB_pos2, NDHC_pos2, PSBM_pos2, PSBN_pos2

38	TVM+F+G	NDHB_pos3

39	TVM+F+I+G	NDHC_pos1, NDHJ_pos1, YCF4_pos2

40	GTR+F+I+G	NDHC_pos3, PSBK_pos3

41	GTR+F+I+G	NDHD_pos3

42	GTR+F+G	NDHE_pos1, RPS2_pos1

43	GTR+F+G	NDHE_pos2, PSBZ_pos2

44	TVM+F+I+G	NDHF_pos1

45	GTR+F+I+G	NDHG_pos1, RPS12_pos3

46	GTR+F+I+G	NDHG_pos3

47	TVM+F+I+G	NDHH_pos2

48	GTR+F+I+G	NDHH_pos3, PETD_pos3

49	TVM+F+G	NDHI_pos1, RPS18_pos2

50	TIM1+F+I+G	NDHI_pos2, NDHJ_pos2

51	TVM+F+G	NDHI_pos3, RPS8_pos3

52	TVM+F+I+G	NDHJ_pos3, NDHK_pos3, PSBH_pos3, PSBM_pos3

53	K81uf+F+I+G	PETA_pos1, RPS16_pos1

54	TVM+F+I+G	PETA_pos2, RPOB_pos2, YCF3_pos2

55	TIM1+F+I+G	PETB_pos1, PSBA_pos1

56	TVM+F+I+G	PETB_pos3, PSBB_pos3

57	TVMef+I+G	PETD_pos1

58	K81uf+F+I+G	PETD_pos2

59	TVMef+G	PETG_pos1, PETN_pos1, PSBE_pos1, PSBI_pos1,

RPS12_pos2, RRN5

60	TN93+F+I+G	PETG_pos2, PSBE_pos2, PSBL_pos2

61	TVM+F+I+G	PETG_pos3, RPL20_pos2, YCF3_pos3

62	TVM+F+I+G	PETL_pos1, PSBL_pos3, RPOA_pos2

63	TVM+F+G	PETL_pos2, PSAJ_pos1, PSBK_pos2, PSBZ_pos1

64	TPM3uf+F+G	PETL_pos3, PSAI_pos3

65	TPM3+F	PETN_pos2, PSBI_pos2

66	TPM3uf+F+G	PETN_pos3, PSBF_pos3, RPL22_pos2

67	K81uf+F+I+G	PSAA_pos1

68	GTR+F+I+G	PSAA_pos2, PSAB_pos2, PSBB_pos2

69	GTR+F+I+G	PSAA_pos3, PSBC_pos3, PSBD_pos3

70	TVM+F+I+G	PSAB_pos1

71	GTR+F+I+G	PSAB_pos3

72	TVM+F+I+G	PSAC_pos1, PSBB_pos1

73	SYM+I	PSAC_pos2

74	GTR+F+I+G	PSAC_pos3, RPL14_pos3, RPL36_pos3

75	TN93ef+G	PSAI_pos1

76	HKY+F+G	PSAI_pos2

77	SYM+G	PSAJ_pos2

78	K81uf+F+G	PSAJ_pos3

79	TIM3+F+I+G	PSBA_pos2, PSBF_pos2

80	GTR+F+I+G	PSBA_pos3

81	TVM+F+I+G	PSBC_pos2, PSBD_pos2

82	TPM2uf+F+G	PSBE_pos3, PSBN_pos3

83	GTR+F+I+G	PSBF_pos1, RPL2_pos2, YCF3_pos1, RRN23, RRN4

84	TIM3ef+I+G	PSBH_pos1, PSBH_pos2

85	HKY+F+I+G	PSBJ_pos2

86	TPM3uf+F+G	PSBJ_pos3, PSBZ_pos3

87	TIM1ef+I+G	PSBK_pos1

88	TIM1+F+G	PSBM_pos1, RPL36_pos1

89	TN93+F+I+G	PSBT_pos2

90	GTR+F+G	PSBT_pos3, RPS19_pos3

91	GTR+F+I+G	RBCL_pos1

92	GTR+F+I+G	RBCL_pos2

93	GTR+F+I+G	RBCL_pos3

94	GTR+F+G	RPL14_pos1

95	TIM1+F+G	RPL14_pos2, RPL23_pos2, RPS7_pos2

96	GTR+F+I+G	RPL16_pos1

97	SYM+I+G	RPL16_pos2

98	TVM+F+I+G	RPL16_pos3

99	TVM+F+I+G	RPL2_pos1, RPS12_pos1, RPS7_pos1

100	GTR+F+I+G	RPL20_pos3, RPOC2_pos3

101	TVM+F+I+G	RPL22_pos3

102	TPM3+F+G	RPL23_pos1

103	TVM+F+G	RPL23_pos3, RPS19_pos2, RPS2_pos2, RPS7_pos3

104	TIM3+F+G	RPL32_pos2

105	TVM+F+I+G	RPL32_pos3, RPS3_pos3

106	TIM1+F+G	RPL33_pos1, RPS11_pos1, RPS8_pos1

107	TVM+F+I+G	RPL33_pos2, RPOC2_pos2

108	TVM+F+G	RPOA_pos3

109	TVM+F+I+G	RPOC1_pos1, RPS4_pos1, RPS4_pos2

110	GTR+F+I+G	RPOC2_pos1

111	TIM3ef+G	RPS11_pos2

112	TVM+F+G	RPS11_pos3

113	TN93+F+G	RPS14_pos2

114	GTR+F+I+G	RPS15_pos2, RPS3_pos2

115	TVM+F+G	RPS16_pos3

116	GTR+F+G	RPS18_pos1

117	TIM1+F+G	RPS19_pos1

118	GTR+F+G	RPS3_pos1

119	GTR+F+G	RPS8_pos2

120	TVM+F+G	YCF2_pos1

121	TVM+F+G	YCF2_pos2

122	TVM+F+G	YCF2_pos3

123	TIM1+F+I+G	RRN16

**(H)**

1	GTR+F+I+G	ACCD_pos1

2	TVM+F+I+G	ACCD_pos2

3	GTR+F+I+G	ACCD_pos3

4	GTR+F+I+G	ATPA_pos1, PSBJ_pos1

5	TVM+F+I+G	ATPA_pos2, ATPB_pos2

6	TVM+F+I+G	ATPA_pos3, NDHH_pos3, PETD_pos3, PSBI_pos3

7	GTR+F+I+G	ATPB_pos1

8	TVM+F+G	ATPB_pos3, PETA_pos3

9	GTR+F+I+G	ATPE_pos1, RPOB_pos1

10	TVM+F+I+G	ATPE_pos2, NDHK_pos2, RPOC1_pos2

11	GTR+F+I+G	ATPE_pos3, ATPF_pos3, RPOB_pos3, RPOC1_pos3,

YCF4_pos3

12	TVM+F+I+G	ATPF_pos1, NDHK_pos1

13	HKY+F+I+G	ATPF_pos2, RPS16_pos2

14	GTR+F+I+G	ATPH_pos1, PSBC_pos1, PSBD_pos1

15	F81+F	ATPH_pos2

16	TVM+F+I+G	ATPH_pos3

17	TVM+F+I+G	ATPI_pos1, PSBF_pos1, YCF3_pos1

18	GTR+F+I+G	ATPI_pos2, NDHE_pos2, PETB_pos2, PSBI_pos2,

PSBL_pos1

19	TVM+F+I+G	ATPI_pos3, NDHK_pos3, PSAI_pos3, PSBH_pos3

20	TVM+F+I+G	CCSA_pos1, PSBF_pos3

21	GTR+F+I+G	CCSA_pos2, NDHF_pos2

22	GTR+F+I+G	CCSA_pos3, NDHF_pos3

23	GTR+F+I+G	CEMA_pos1, RPOA_pos1, YCF4_pos1

24	TVM+F+G	CEMA_pos2, PSBL_pos3, RPL23_pos3

25	TVM+F+G	CEMA_pos3

26	TVMef+G	CLPP_pos1

27	GTR+F+I+G	CLPP_pos2

28	GTR+F+I+G	CLPP_pos3, RPS14_pos3

29	TVM+F+G	INFA_pos1, INFA_pos2, RPS14_pos1

30	TVM+F+I+G	INFA_pos3, RPL22_pos3, RPL32_pos3

31	GTR+F+G	MATK_pos1

32	TVM+F+I+G	MATK_pos2

33	TVM+F+I+G	MATK_pos3

34	GTR+F+I+G	NDHA_pos1, NDHD_pos1

35	TVM+F+I+G	NDHA_pos2, NDHD_pos2, NDHG_pos2

36	TVM+F+I+G	NDHA_pos3, NDHE_pos3

37	TIM3+F+I+G	NDHB_pos1, PSBT_pos1, RPL23_pos2

38	TIM2+F+I+G	NDHB_pos2, NDHC_pos2, PSBM_pos2, PSBN_pos2

39	TVM+F+G	NDHB_pos3

40	TVM+F+I+G	NDHC_pos1, NDHJ_pos1, PSBN_pos1, YCF4_pos2

41	GTR+F+I+G	NDHC_pos3, PSBK_pos3

42	GTR+F+I+G	NDHD_pos3

43	TVM+F+G	NDHE_pos1, NDHI_pos1, RPL33_pos1, RPL36_pos2

44	TVM+F+I+G	NDHF_pos1

45	GTR+F+G	NDHG_pos1, PSBJ_pos3

46	GTR+F+I+G	NDHG_pos3, RPL33_pos3, RPOA_pos3

47	GTR+F+I+G	NDHH_pos1, RPOC1_pos1

48	TVM+F+I+G	NDHH_pos2

49	K81uf+F+I+G	NDHI_pos2, NDHJ_pos2, PSBM_pos1

50	GTR+F+I+G	NDHI_pos3

51	TVM+F+I+G	NDHJ_pos3, PETB_pos3, PSBM_pos3

52	K81uf+F+I+G	PETA_pos1, RPS16_pos1

53	TVM+F+I+G	PETA_pos2, RPOB_pos2, YCF3_pos2

54	TIM1+F+I+G	PETB_pos1, PSBA_pos1

55	SYM+I+G	PETD_pos1

56	K81uf+F+I+G	PETD_pos2

57	TVMef+G	PETG_pos1, PETN_pos1, PSBE_pos1, PSBI_pos1

58	TN93+F+I+G	PETG_pos2, PSBE_pos2, PSBL_pos2

59	TVM+F+I+G	PETG_pos3, PETN_pos3, RPL20_pos2, YCF3_pos3

60	TN93+F+I+G	PETL_pos1, PSAI_pos1, RPS14_pos2

61	GTR+F+G	PETL_pos2, PSAJ_pos1, PSBZ_pos1

62	TPM2uf+F+G	PETL_pos3, PSBE_pos3, PSBN_pos3

63	TIM3+F+G	PETN_pos2, PSBZ_pos2

64	K81uf+F+I+G	PSAA_pos1

65	GTR+F+I+G	PSAA_pos2, PSAB_pos2, PSBB_pos2

66	GTR+F+I+G	PSAA_pos3, PSBC_pos3, PSBD_pos3

67	TVM+F+I+G	PSAB_pos1

68	GTR+F+I+G	PSAB_pos3

69	TVM+F+I+G	PSAC_pos1, PSBB_pos1

70	SYM+I	PSAC_pos2

71	GTR+F+I+G	PSAC_pos3, RBCL_pos3

72	HKY+F+G	PSAI_pos2

73	GTR+F+I+G	PSAJ_pos2, PSBK_pos2

74	K81uf+F+G	PSAJ_pos3

75	TIM3+F+I+G	PSBA_pos2, PSBF_pos2

76	GTR+F+I+G	PSBA_pos3

77	TVM+F+I+G	PSBB_pos3

78	TVM+F+I+G	PSBC_pos2, PSBD_pos2

79	TIM3ef+I+G	PSBH_pos1, PSBH_pos2

80	HKY+F+I+G	PSBJ_pos2

81	TIM1ef+I+G	PSBK_pos1

82	TN93+F+I+G	PSBT_pos2

83	GTR+F+G	PSBT_pos3, RPS16_pos3

84	TIM1+F+I	PSBZ_pos3

85	GTR+F+I+G	RBCL_pos1

86	GTR+F+I+G	RBCL_pos2

87	GTR+F+I+G	RPL14_pos1

88	TN93+F+I+G	RPL14_pos2

89	TVM+F+I+G	RPL14_pos3

90	GTR+F+G	RPL16_pos1

91	SYM+G	RPL16_pos2

92	TVM+F+I+G	RPL16_pos3, RPS11_pos3

93	GTR+F+G	RPL2_pos1, RPL2_pos2

94	K81uf+F+I+G	RPL2_pos3

95	TVM+F+G	RPL20_pos1, RPL22_pos1, RPL32_pos1, RPS15_pos1

96	GTR+F+I+G	RPL20_pos3, RPOC2_pos3

97	TN93+F+G	RPL22_pos2

98	TPM3+F+G	RPL23_pos1

99	GTR+F+G	RPL32_pos2

100	TVM+F+I+G	RPL33_pos2, RPOA_pos2, RPOC2_pos2

101	K81uf+F+G	RPL36_pos1, RPS7_pos1

102	TIM3+F+G	RPL36_pos3

103	GTR+F+I+G	RPOC2_pos1

104	GTR+F+G	RPS11_pos1, RPS18_pos2, RPS8_pos1

105	TIM3ef+G	RPS11_pos2

106	TVM+F+G	RPS12_pos1

107	K80+G	RPS12_pos2

108	K81uf+F+G	RPS12_pos3

109	TVM+F+I+G	RPS15_pos2

110	TVM+F+G	RPS15_pos3

111	K81uf+F+G	RPS18_pos1

112	GTR+F+I+G	RPS18_pos3

113	K81uf+F+G	RPS19_pos1

114	GTR+F+G	RPS19_pos2, RPS8_pos2

115	GTR+F+G	RPS19_pos3

116	GTR+F+G	RPS2_pos1

117	TPM3+F+G	RPS2_pos2

118	GTR+F+G	RPS2_pos3, RPS8_pos3

119	GTR+F+G	RPS3_pos1

120	GTR+F+I+G	RPS3_pos2

121	TVM+F+I+G	RPS3_pos3

122	K81uf+F+G	RPS4_pos1

123	TVM+F+G	RPS4_pos2

124	GTR+F+G	RPS4_pos3

125	K81uf+F+I	RPS7_pos2, RRN4

126	GTR+F+G	RPS7_pos3

127	TVM+F+G	YCF2_pos1

128	TVM+F+G	YCF2_pos2

129	TVM+F+G	YCF2_pos3

130	SYM+G	RRN16

131	SYM+G	RRN23

132	K80+G	RRN5
