## Supplementary figures and images for "Placing very long branch taxa in the plant tree of life: a case study with Dioscoreales mycoheterotrophs"

### Appendix S3

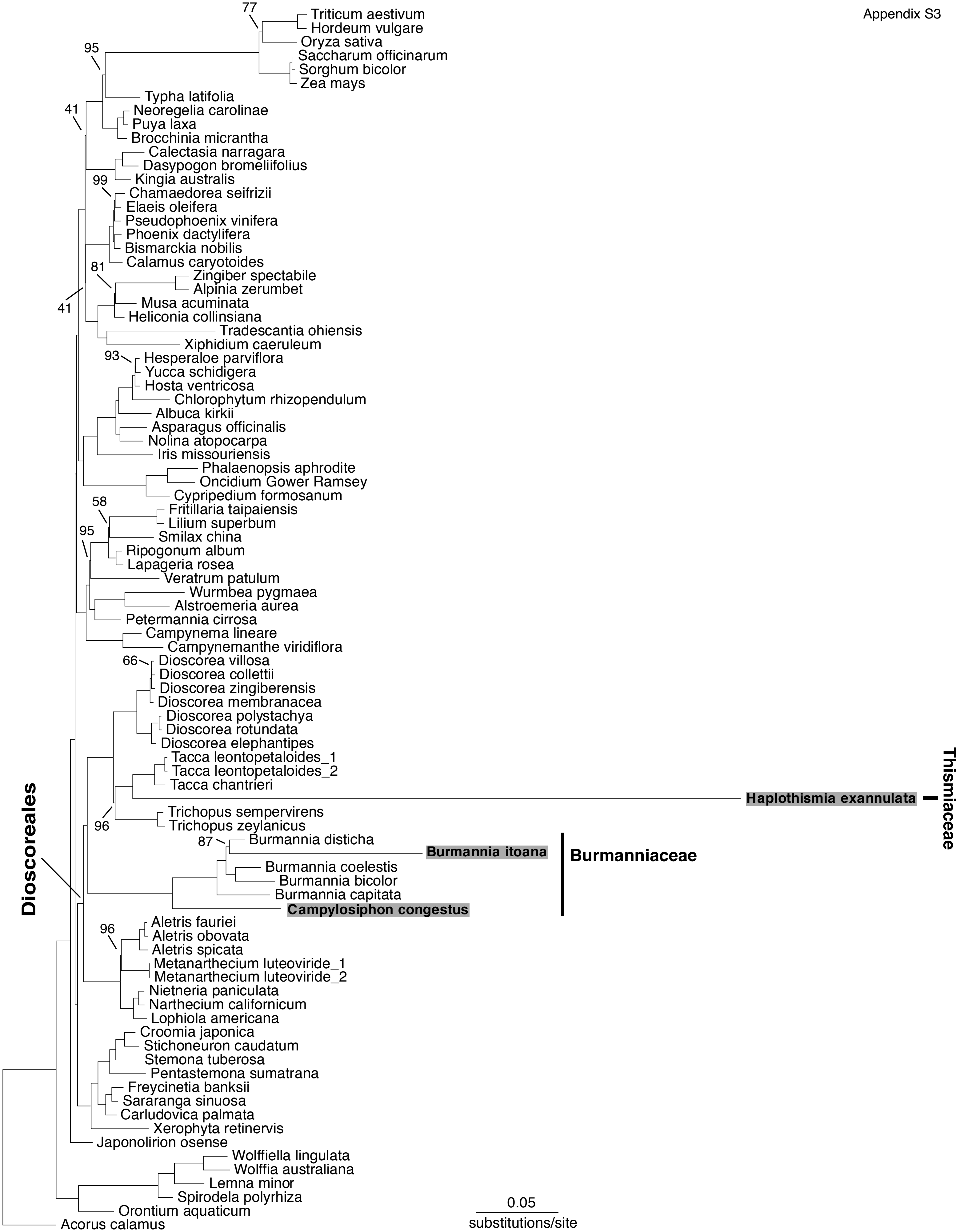

### Appendix S4

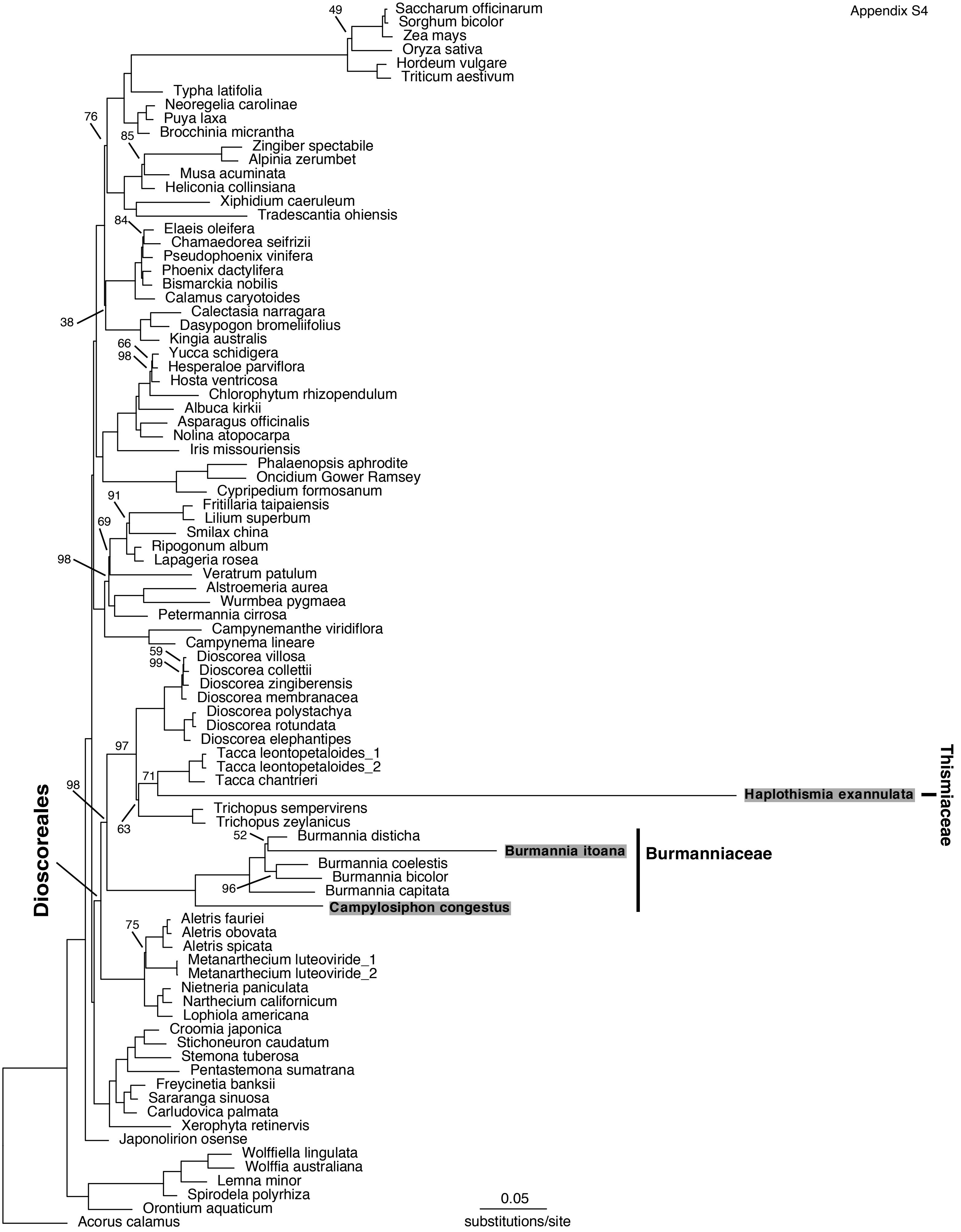

### Appendix S5

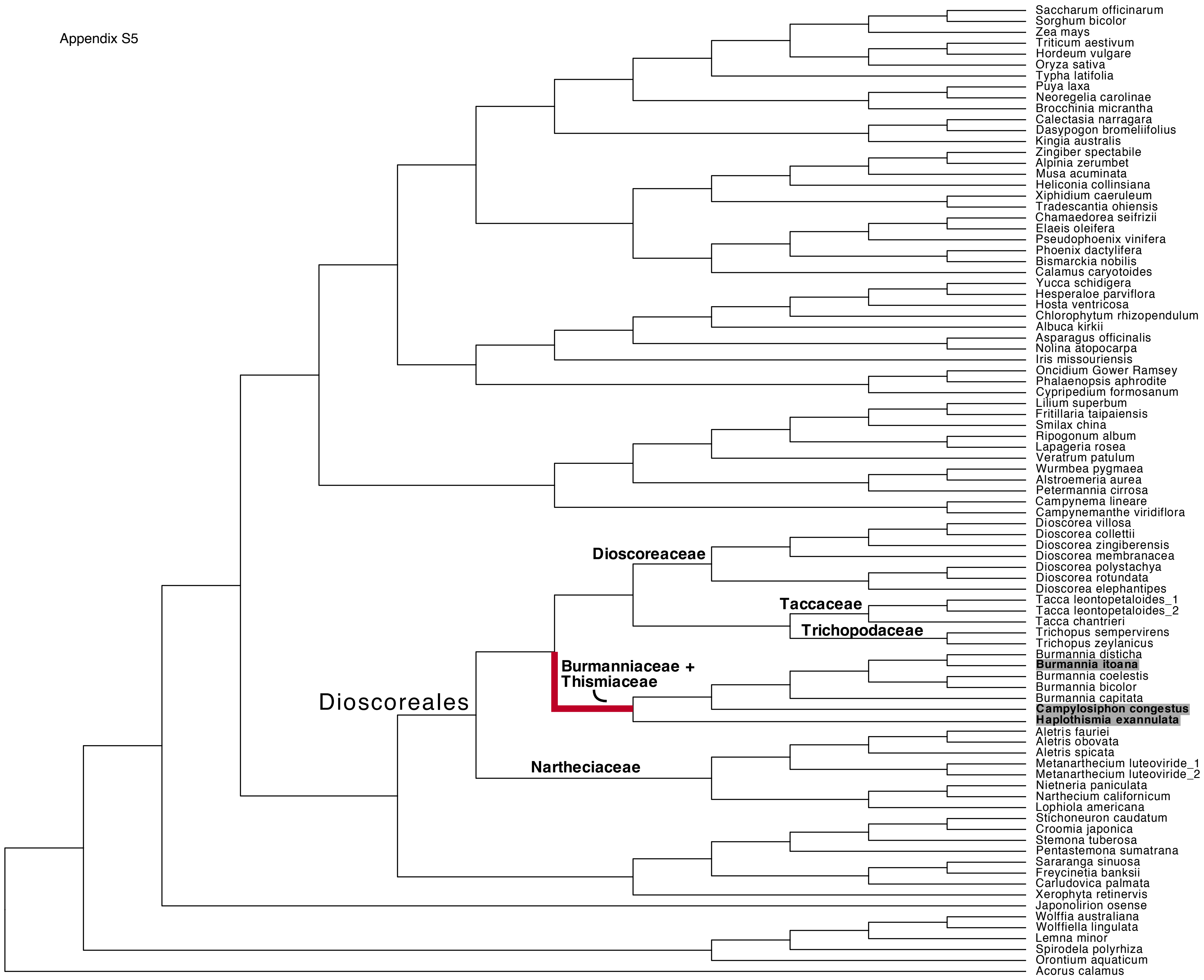

### Appendix S6

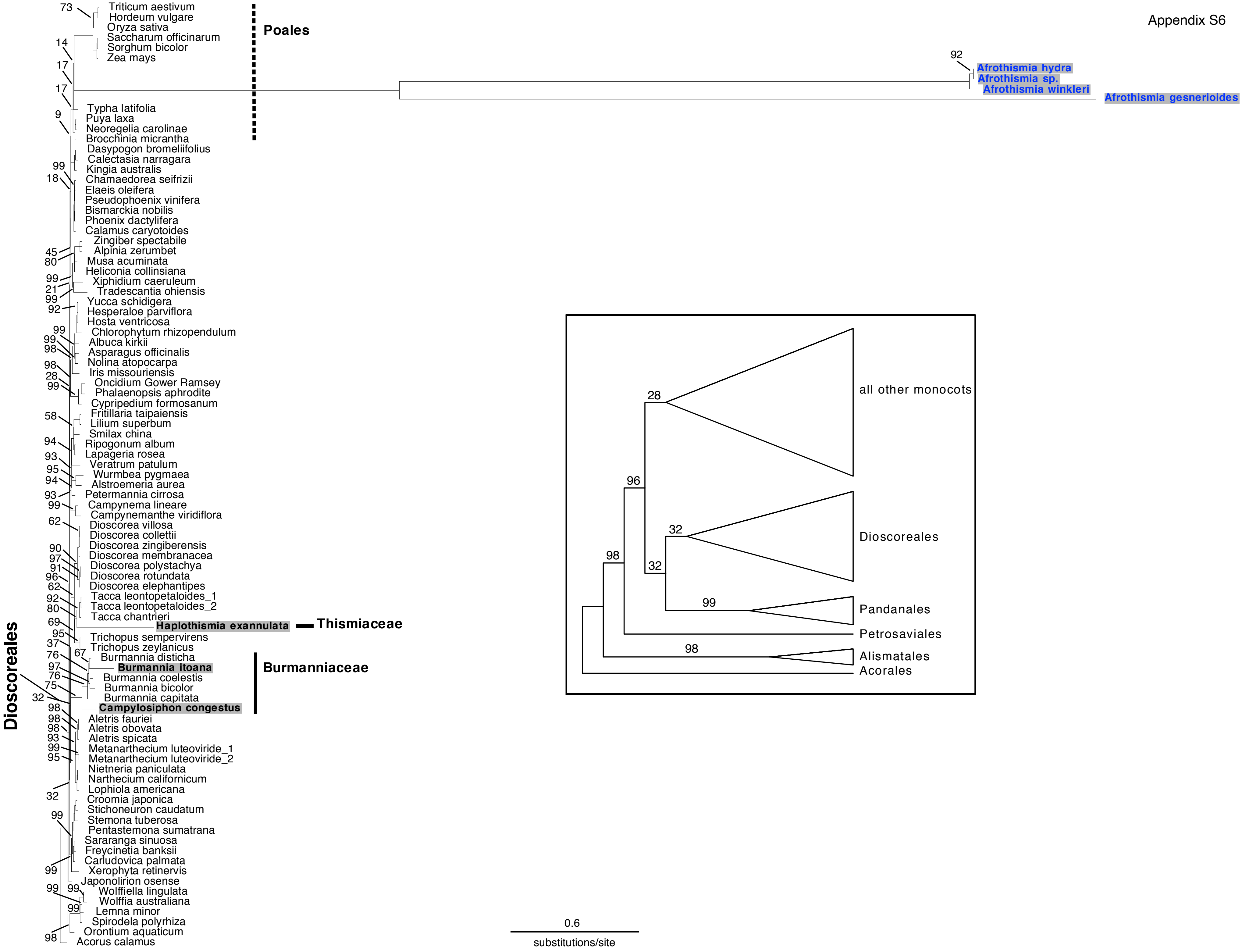

### Appendix S7

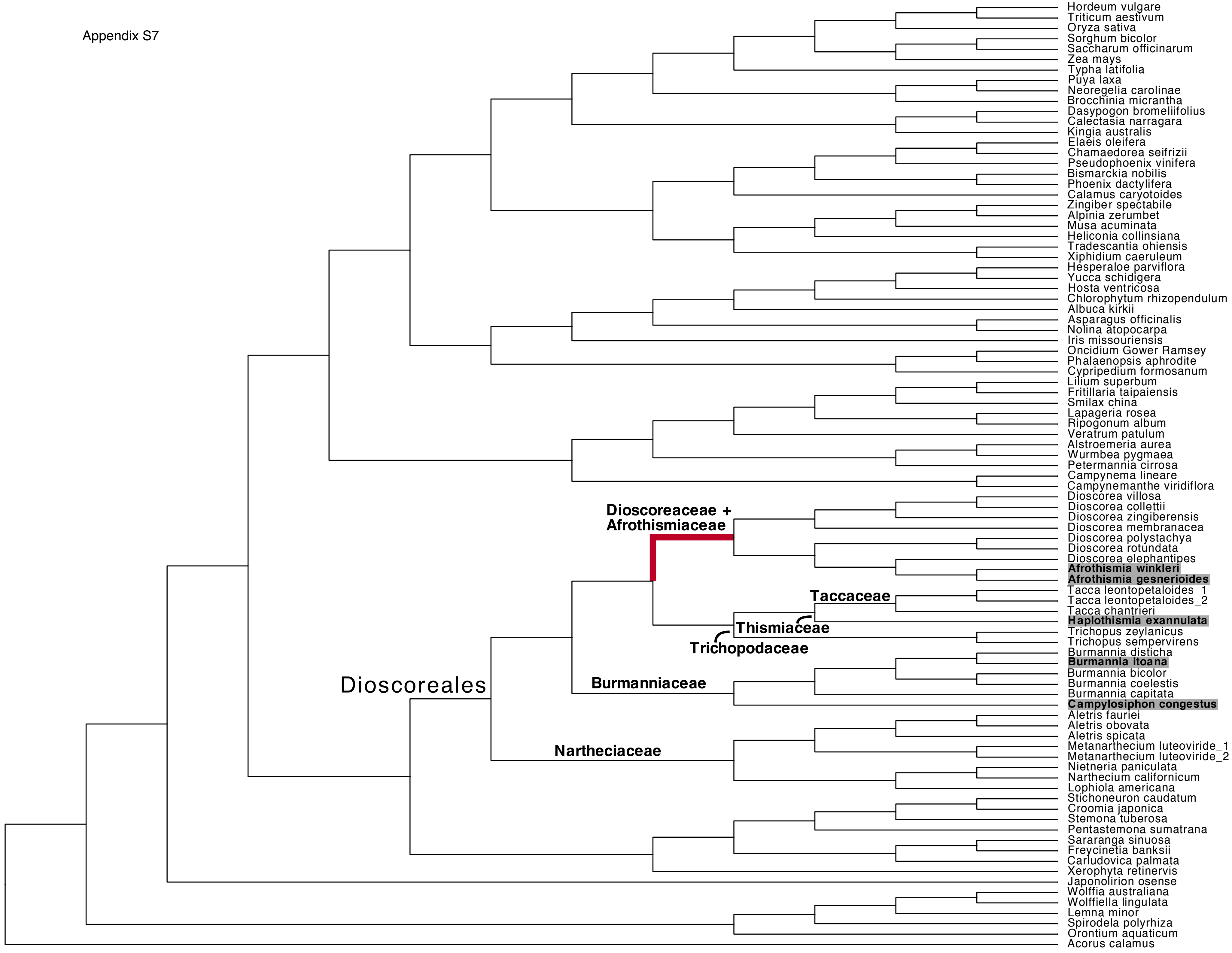

### Appendix S8

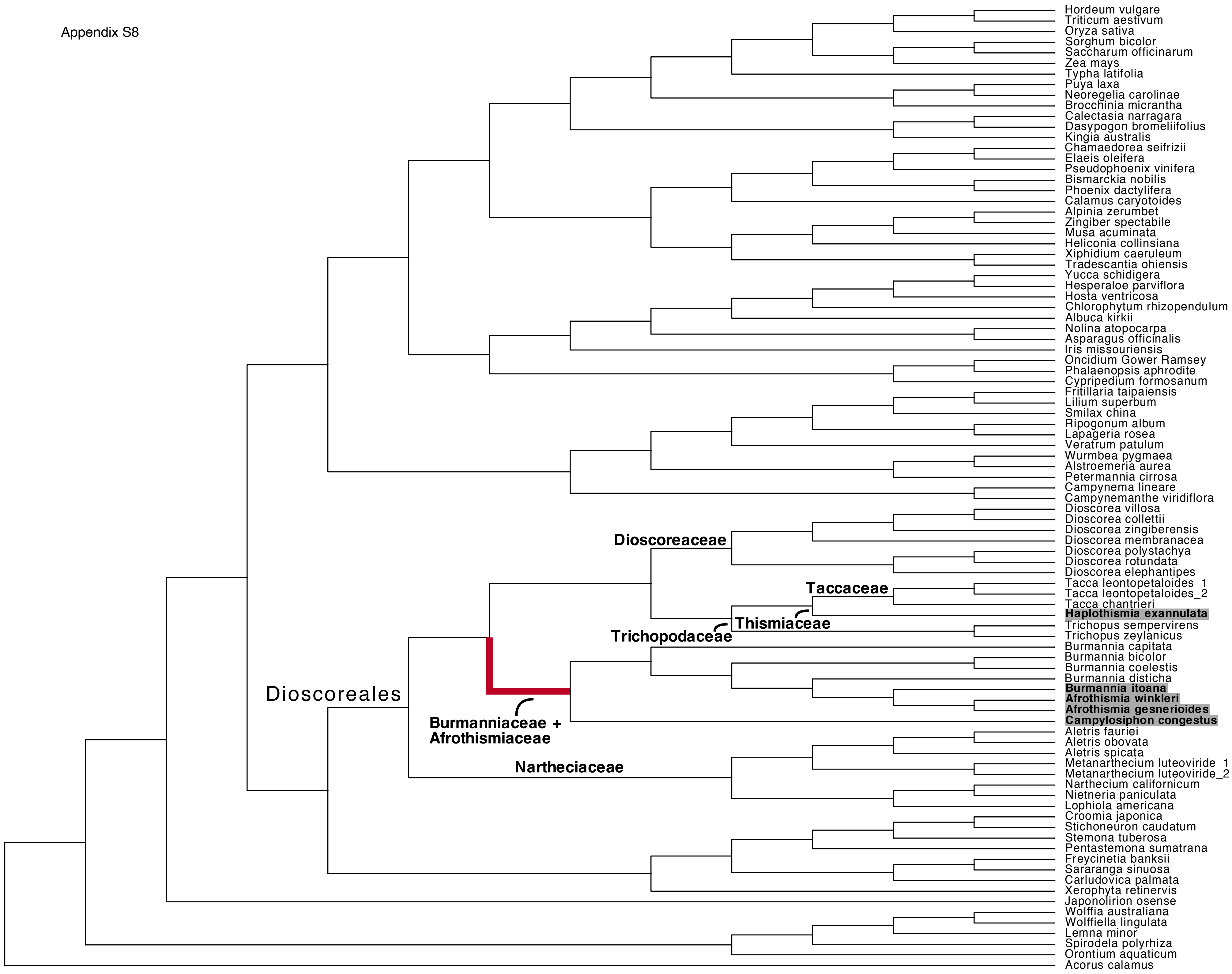

### Appendix S9

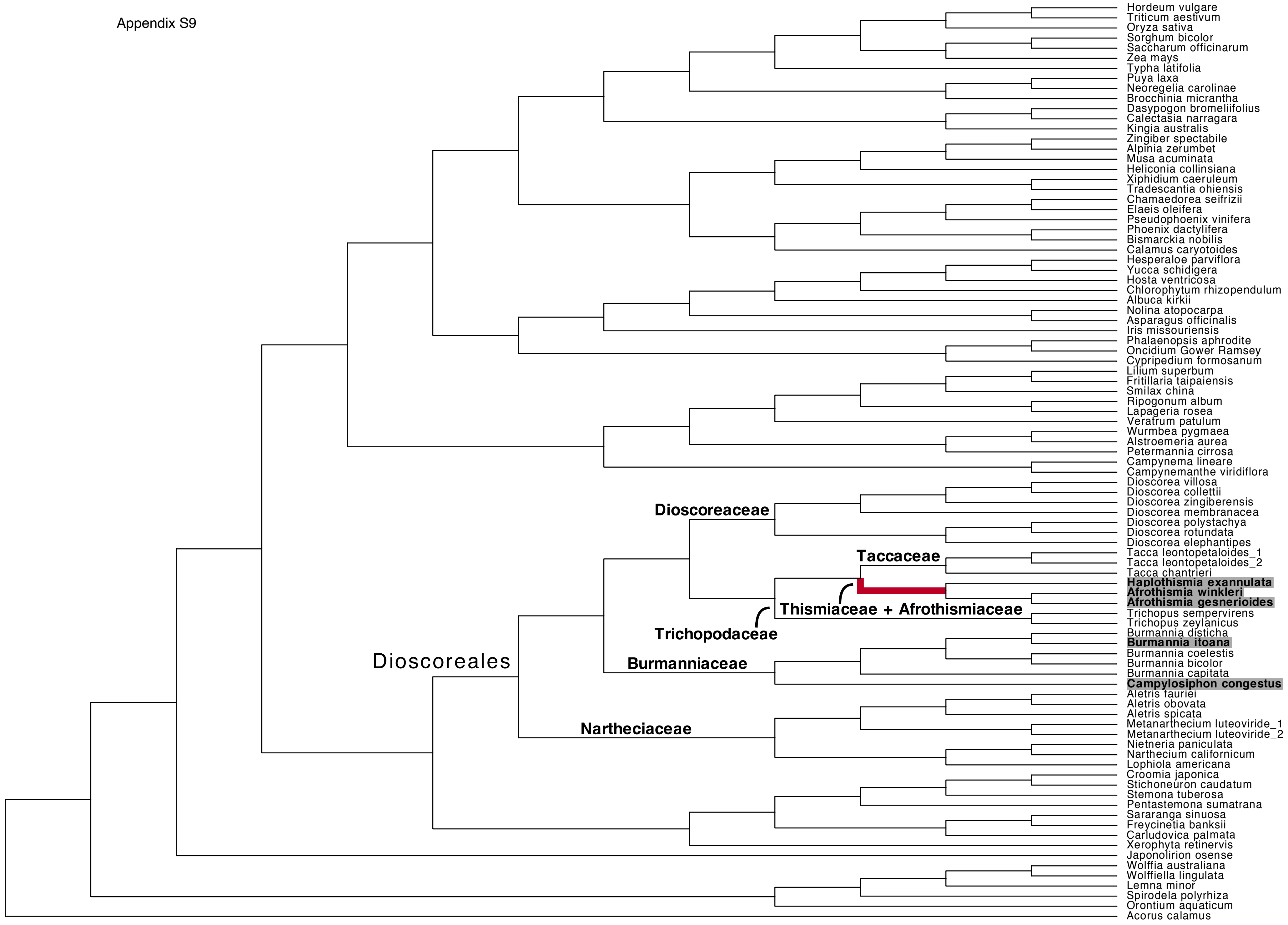

### Appendix S10

Appendix S10

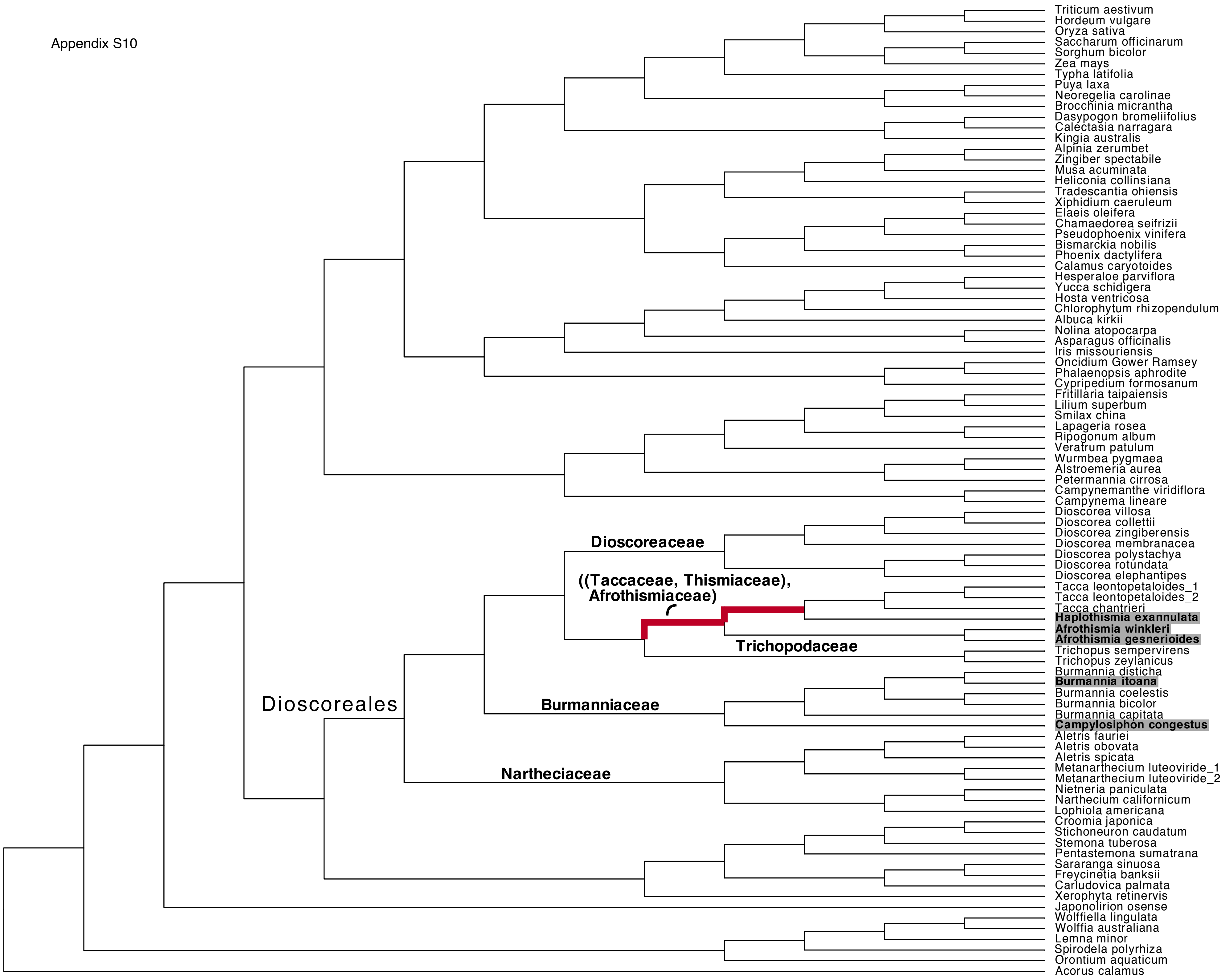

### Appendix S11

Appendix S11

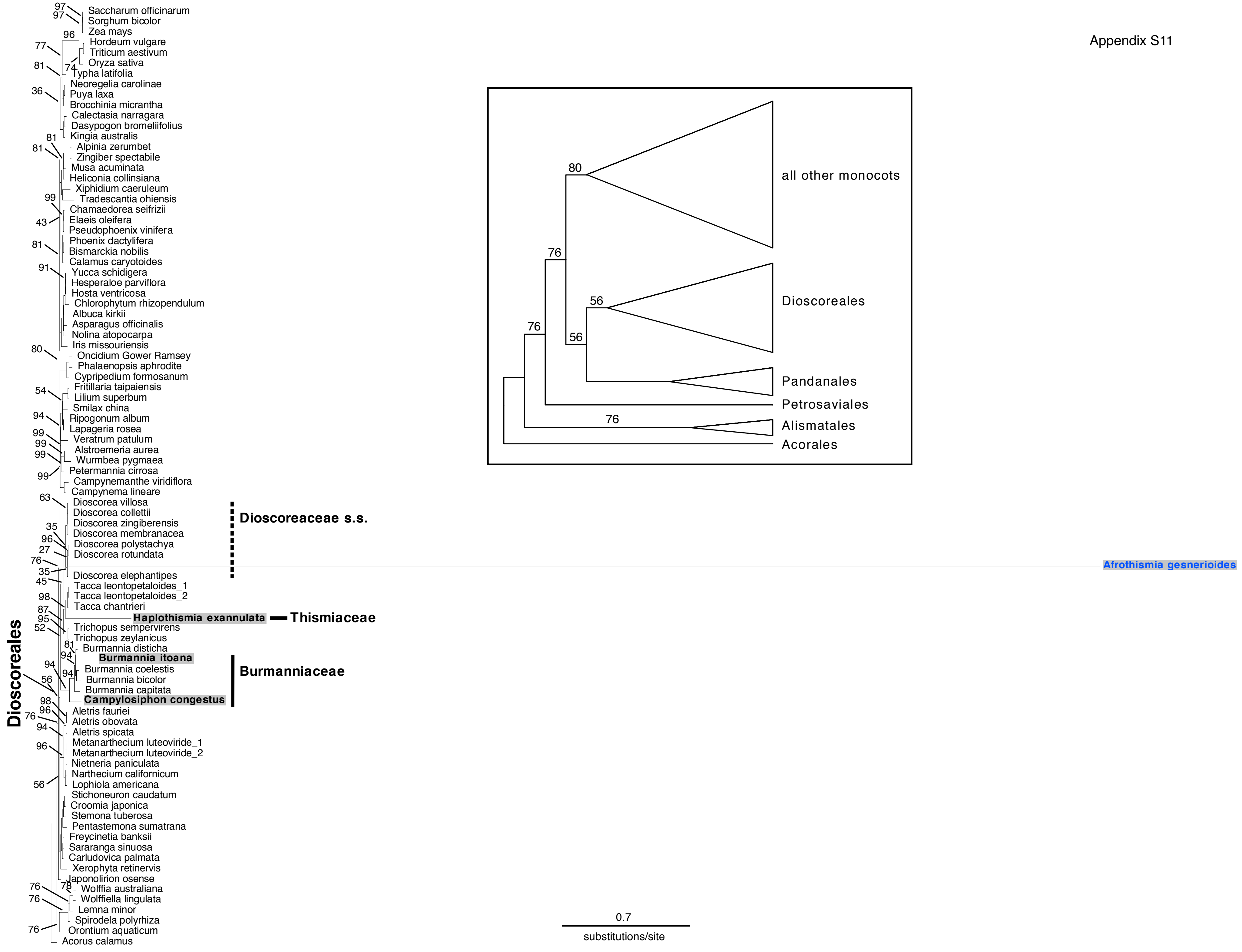

### Appendix S12

Appendix S12

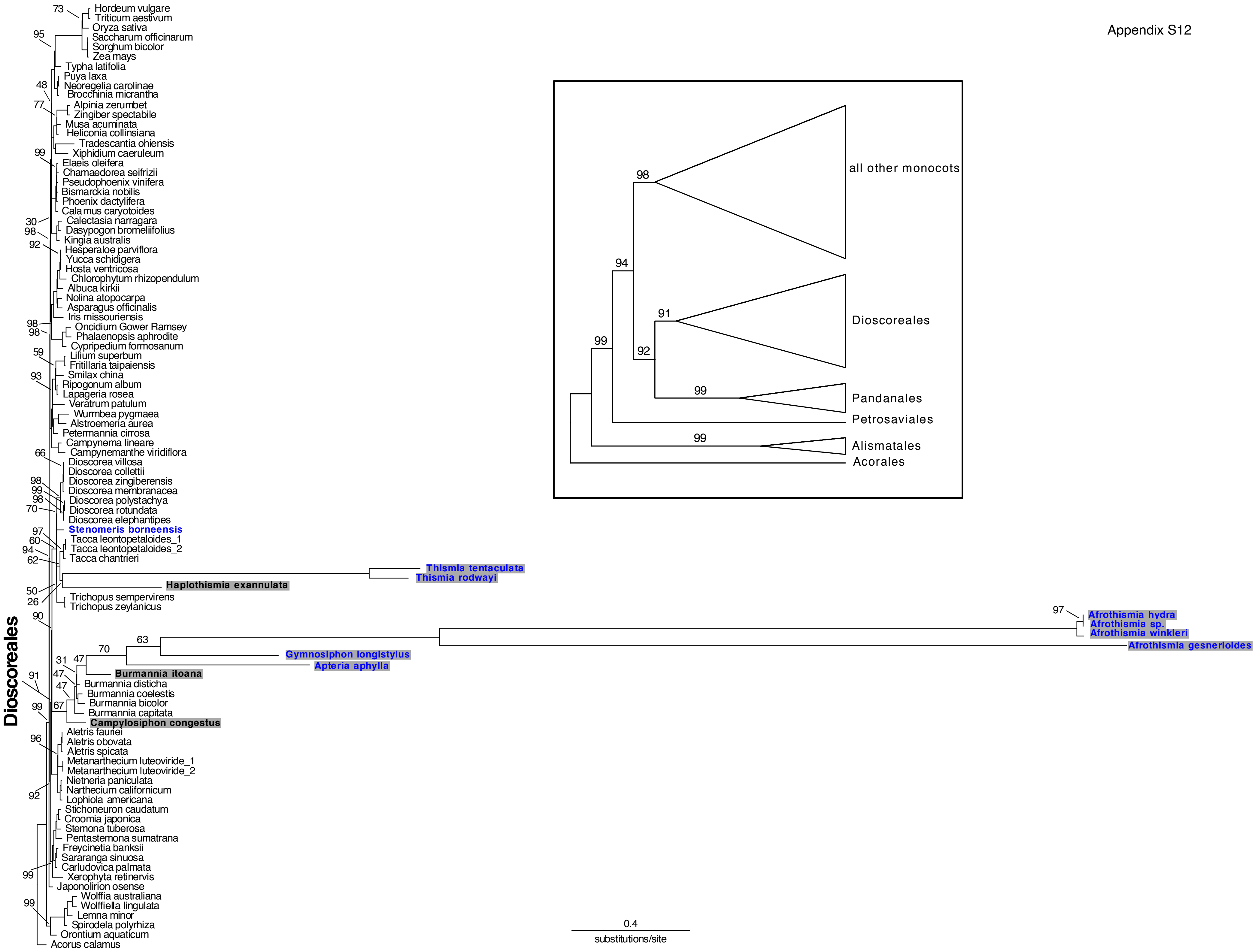

### Appendix S13

Appendix S13

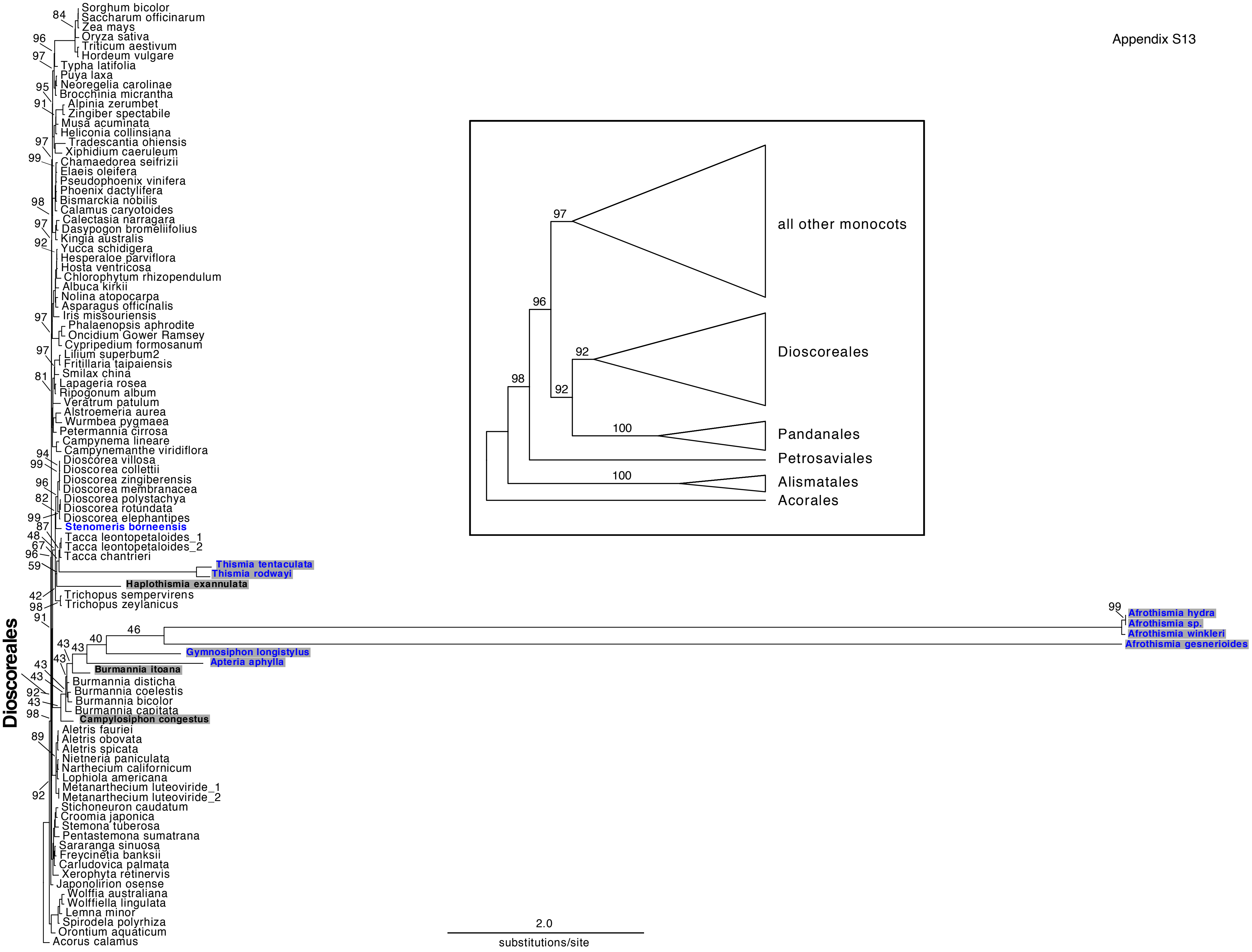

### Appendix S14

Appendix S14

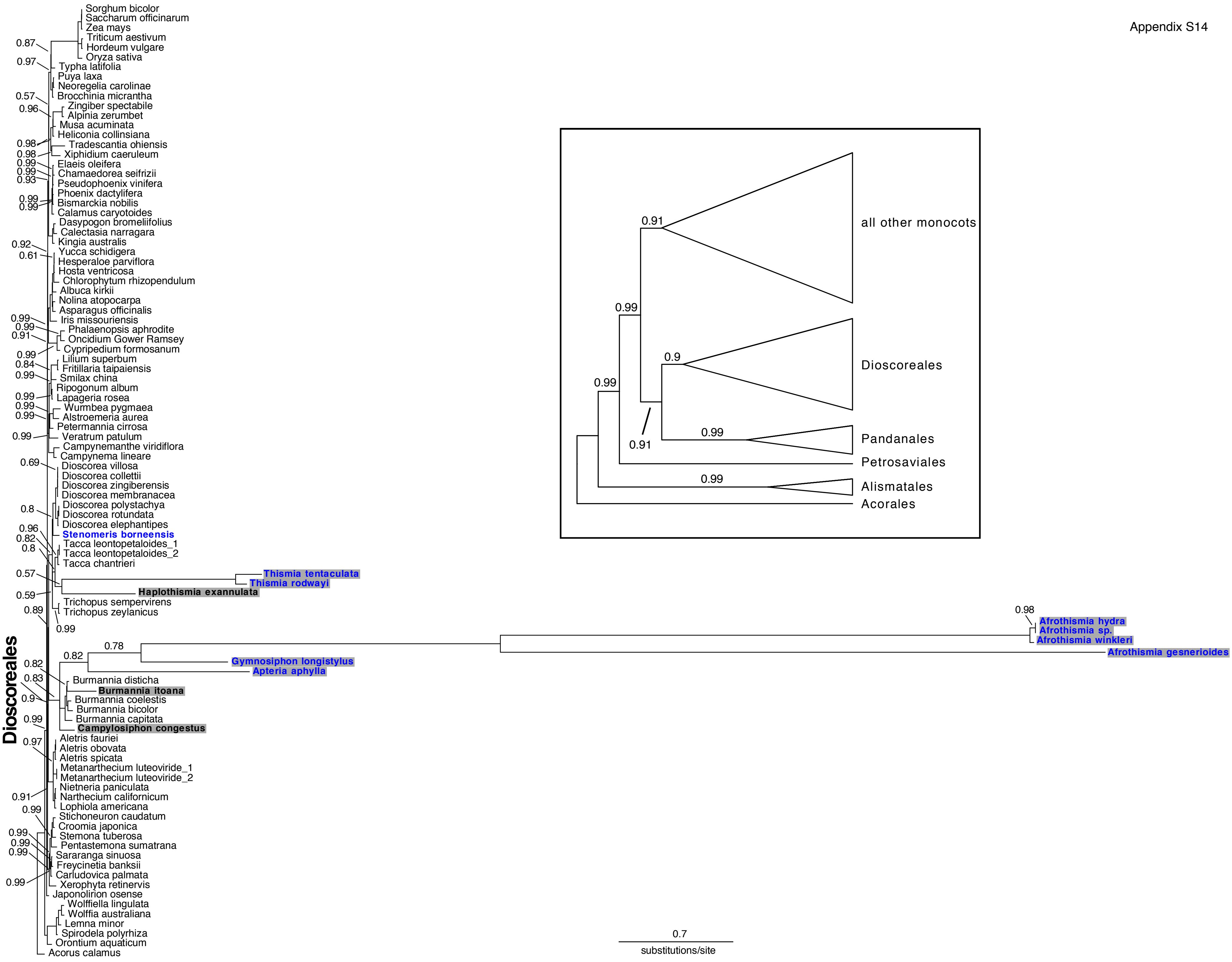

### Appendix S15

Appendix S15

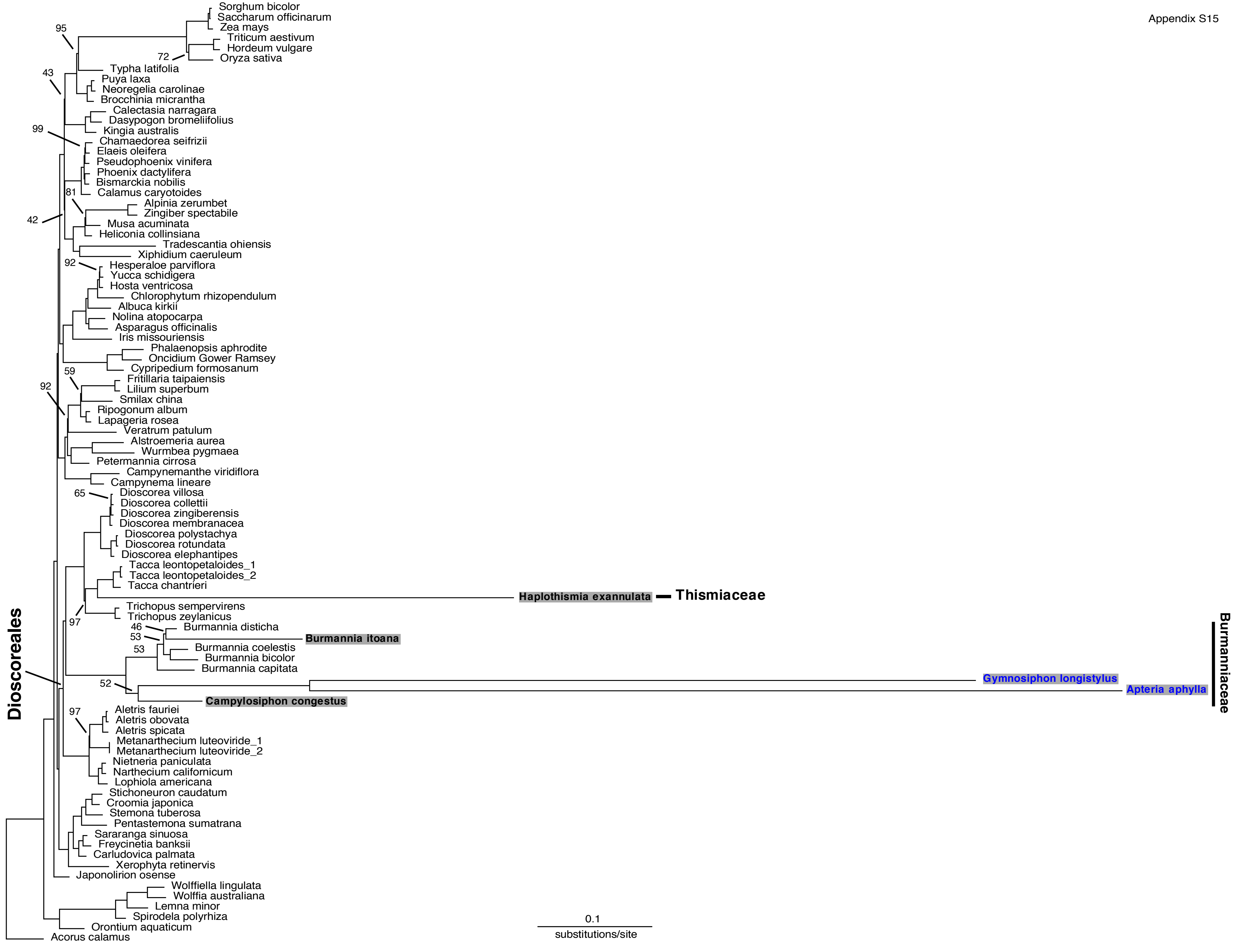

### Appendix S16

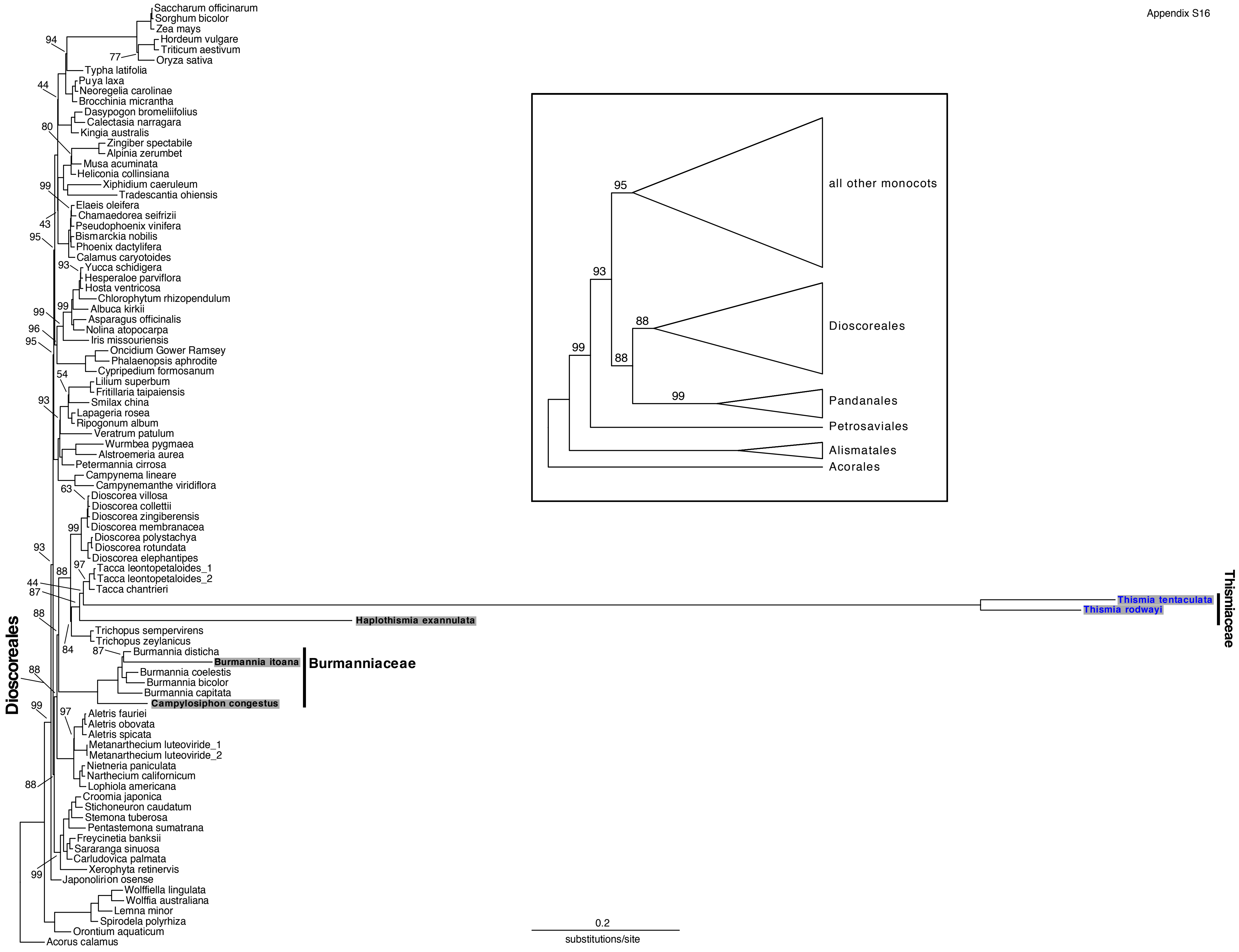

### Appendix S17

Appendix S17

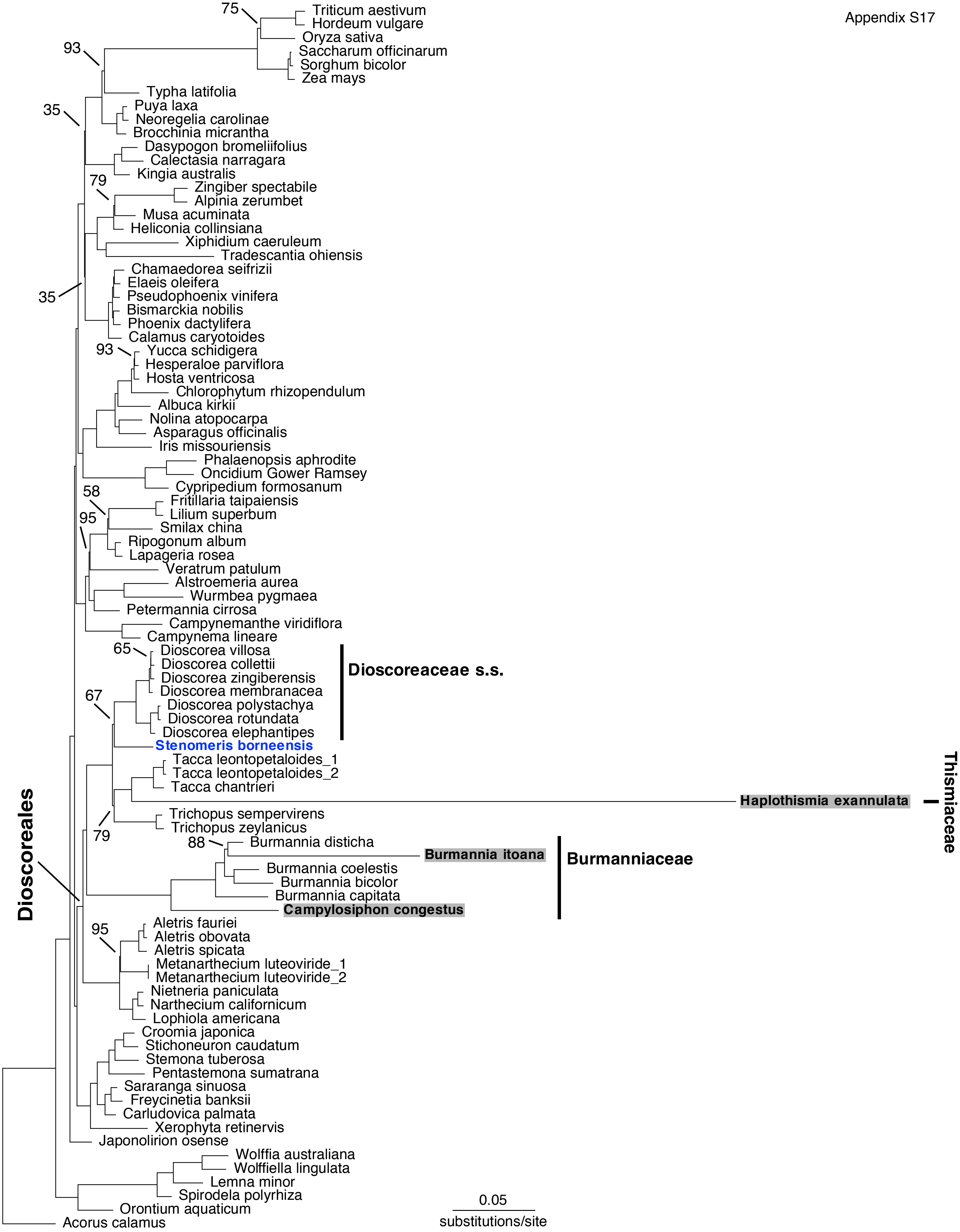
